# Identification of UCB-9721 as a potent inhibitor of MyoA, the essential class XIV myosin motor of apicomplexan parasites

**DOI:** 10.64898/2026.06.18.733251

**Authors:** Anne K Snyder, Filomena Tedesco, Anne Kelsen, Eddie Wehri, Binod Nepal, Jose Teixeira, Emmett Dews, Sachie Kanatani, Kyle Kasprzak, Jonathan Oliva, Kaesi Morelli, Samantha Beck Previs, Bruno Martorelli di Genova, Fran Sverdrup, Martin J Boulanger, Photini Sinnis, Christopher D Huston, Sandhya Kortagere, David M Warshaw, Julia Schaletzky, Nicholas J Westwood, Gary E Ward

## Abstract

The virulence of *Toxoplasma gondii* and other apicomplexan parasites relies on a unique form of cellular motility driven by MyoA, an unconventional class XIV myosin motor protein. To identify new chemical probes for investigating the molecular mechanisms of parasite motility, we screened over 50,000 small molecules for inhibitors of *T. gondii* MyoA (TgMyoA). The top hit from the screen, UCB-9721, is almost 40-fold more potent as an inhibitor of TgMyoA actin-activated ATPase activity than the previously described TgMyoA inhibitor, KNX-002, and 45-fold more potent at inhibiting parasite motility, with no detectable toxicity towards mammalian cells. UCB-9721 also inhibited the motility and/or growth of the related apicomplexan parasites *Plasmodium falciparum, Cryptosporidium parvum,* and *Babesia duncani,* suggesting that this compound will be a useful new chemical probe for studying motility and MyoA function in apicomplexan parasites more broadly. While UCB-9721 and KNX-002 were identified independently, they share a similar chemical scaffold. To determine why UCB-9721 is so much more potent than KNX-002 and to inform future development of this inhibitor class, we undertook comparative molecular docking analyses, targeted TgMyoA mutagenesis, and a directed structure-activity relationship analysis. The results identified the sulfonamide group of UCB-9721 and its hydrogen bond interactions with R249, E275 and a stabilized water network within the TgMyoA binding pocket as key to the compound’s increased potency. Further development of UCB-9721, informed by the results presented here, may transform this promising new chemical class into actionable drug development leads against this important group of human and animal pathogens.

## Introduction

Nearly one third of the world’s population is infected with the protozoan parasite *Toxoplasma gondii* (1, 2). Although most infections are subclinical, acute toxoplasmosis can be life threatening in immunocompromised and pregnant individuals. In the immunocompromised, pre-emptive antiparasitic treatment can reduce the risk of toxoplasmic encephalitis; however, serious side effects of these drugs and relapse of infection are common (3–8). The toxic and potentially teratogenic effects of currently available drug regimens also make management of infection in pregnant individuals difficult (6–12). New and better-tolerated drugs are therefore needed for the prevention and treatment of clinical toxoplasmosis.

To address the need for novel therapeutics, a better understanding of the biology of *T. gondii* and the mechanisms underlying parasite virulence is required. One promising area for investigation is parasite motility. *T. gondii* and related apicomplexan parasites (including those that cause malaria, cryptosporidiosis, and babesiosis) employ an unusual form of substrate-dependent gliding motility to invade into and egress from host cells, migrate across biological barriers, and disseminate throughout host tissues (13–22). Motility in apicomplexan parasites is driven, in part, by an unconventional class XIV myosin molecular motor. This motor lies just beneath the parasite plasma membrane and consists of a myosin heavy chain, MyoA, and its associated regulatory and essential light chains (23–25). *T. gondii* MyoA (TgMyoA) is a single-headed motor that is highly conserved amongst apicomplexans but distinctly different from mammalian myosins, as TgMyoA lacks both a conventional tail and several of the conserved residues in mammalian myosins that regulate actomyosin activity (26–29). TgMyoA also displays unique biochemical properties compared to mammalian myosins. The parasite motor has a strikingly low duty-ratio (the proportion of the ATPase cycle time that myosin is strongly bound to actin) of approximately 1% yet moves actin filaments at the same speed as skeletal muscle myosin – a double-headed motor with a 5-fold higher duty ratio (30, 31). In parasites, knocking out *TgMyoA* severely reduces motility, host cell invasion, and host cell egress (32–35). Consequently, parasites that lack TgMyoA are completely avirulent in animal models of infection (35), highlighting the potential of TgMyoA as a drug target and underscoring the need to better understand the mechanisms underlying parasite motility.

We have used high-throughput small-molecule screening to identify novel chemical probes that can be used to study how TgMyoA and motor-associated proteins generate the force necessary for parasite motility. In a previous screen (36), we identified KNX-002 as a micromolar inhibitor of TgMyoA’s actin-activated ATPase activity, parasite motility, and parasite growth in culture. By introducing a mutation into TgMyoA that reduced parasite sensitivity to KNX-002, we were able to demonstrate the specificity of the compound for TgMyoA in our cell-based assays. KNX-002 slowed the progression of disease in a mouse model of infection, demonstrating for the first time that TgMyoA is a druggable target *in vivo* (36). While KNX-002 provided proof-of-principle that TgMyoA can be targeted by small molecule inhibitors, its relatively low potency (IC_50_ for TgMyoA actin-activated ATPase activity = 2.8 µM and IC_50_ for parasite growth = 16.2 µM; ref. (36)) and hepatotoxicity make KNX-002 a less-than-ideal chemical probe for studying TgMyoA function.

We report here the results from an independent high-throughput screen that was undertaken to identify more potent and less toxic inhibitor(s) of TgMyoA. The top hit from the screen – UCB-9721 – was almost 40-fold more potent at inhibiting TgMyoA actin-activated ATPase activity than KNX-002. We showed that UCB-9721 is active not only against *T. gondii* but several other apicomplexan parasites as well, including *Plasmodium falciparum* (the causative agent of severe malaria), *Cryptosporidium parvum* (a significant cause of infant dehydration, malnutrition, and death globally), and *Babesia duncani* (which causes the emerging infectious blood disease babesiosis), suggesting it will be of broad value as a chemical probe of class XIV myosin function in these important human and animal pathogens. Although they were identified independently, UCB-9721 and KNX-002 share a similar chemical scaffold. Using molecular docking, targeted TgMyoA mutagenesis, and directed structure-activity relationship analysis, we identified the sulfonamide group of UCB-9721 as key to this compound’s increased potency compared to KNX-002.

## Results

### Identification of UCB-9721 as an inhibitor of *T. gondii* myosin A

To identify novel inhibitors of TgMyoA, we conducted an unbiased high-throughput screen for compounds that affected TgMyoA’s actin-activated ATPase activity. The top hit from the 50,422-compound screen was UCB-9721 (Figure 1), which inhibited TgMyoA’s actin-activated ATPase activity with an IC_50_ = 0.031 μM (95% confidence interval [95% C.I.] = 0.029-0.033 μM). UCB-9721 is therefore much more potent than KNX-002, the most potent previously described inhibitor of TgMyoA actin-activated ATPase activity (IC_50_ = 2.8 µM; ref. (36)). Since UCB-9721 is not commercially available, a scheme for synthesizing and purifying the compound was developed (see Figure S1A and Supporting Information). The IC_50_ of the re-synthesized UCB-9721 for TgMyoA actin-activated ATPase was 0.047 μM (95% C.I. = 0.042-0.053 μM; Figure S1B), which was comparable to the original library compound (0.031 μM). All subsequent assays used the re-synthesized compound of defined purity (see Supporting Information).

**Figure 1.**
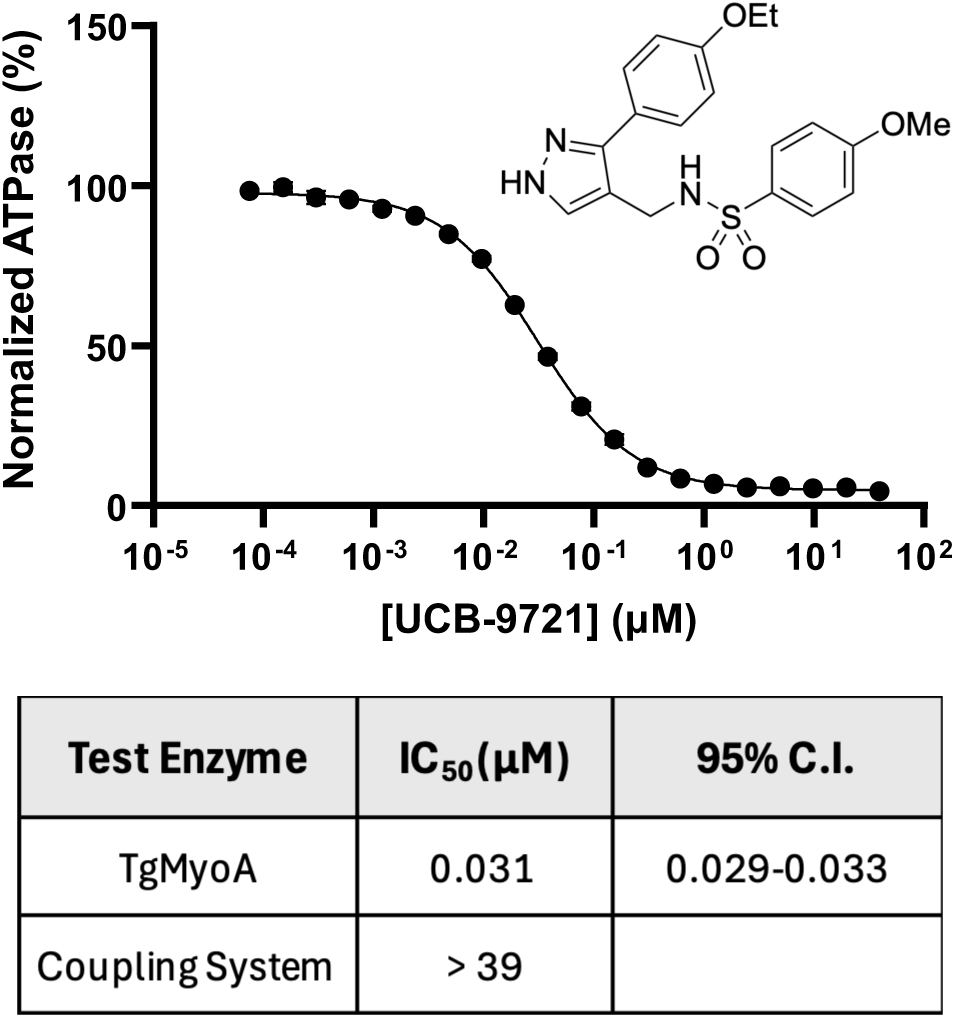
UCB-9721 inhibits TgMyoA with nanomolar potency. Top: Structure of UCB-9721 and dose-dependent inhibition of TgMyoA actin-activated ATPase activity, normalized to DMSO (vehicle) control. The table shows the calculated IC_50_ of UCB-9721 from the screening plates for TgMyoA actin-activated ATPase, with 95% confidence interval (C.I.), and lack of activity against the coupled enzyme system used for the assay, with hexokinase as an unrelated ATPase.

As expected for an inhibitor of TgMyoA actin-activated ATPase activity, UCB-9721 also inhibited TgMyoA motor function in an *in vitro* motility assay (30, 37, 38) that measures the speed at which fluorescently labeled actin filaments are propelled by myosin adsorbed to a glass slide. With increasing concentration of compound, the actin filament speed decreased as more myosin motors were inhibited (Figure 2A and Videos S1A, B). Lower concentrations of compound were required to inhibit actin-activated ATPase activity (Figure 1) than actin filament speed (Figure 2A), because each myosin inactivated by compound in the ATPase assay contributes incrementally to the decrease in activity, whereas filament speed only becomes sensitive to the inhibitor once the density of functional myosin motors on the glass surface falls below the threshold necessary to maintain maximum speed (38). Without knowing the exact myosin surface density, we estimated the effective reduction in functional myosin density caused by UCB-9721 by comparison to serially diluted TgMyoA assayed in the absence of compound (Figure 2D-F). The presence of 2.5 μM UCB-9721 in the assay reduced the actin filament speed to approximately the same extent as a 10-fold myosin dilution (44.5 *vs* 48.5%, respectively; Figures 2A, 2D). This suggests that 2.5 μM UCB-9721 functionally inactivates ∼90% of the myosin present in the assay. In addition to the compound’s effect on speed, the number of filaments moving (expressed as the fraction of the total filaments bound to the coverslip) also decreased with increasing concentration of UCB-9721 (Figure 2B). Combining these two effects into a single “filament motility index” (*i.e.,* the fraction of filaments moving multiplied by the mean speed of the filaments that do move) provides a more complete measure of the overall effect of the compound on motor function in the assay (Figure 2C). Further evidence that UCB-9721 effectively reduces the density of functional motors that keep actin filaments in contact with the surface is the decrease in the absolute number of bound actin filaments with increasing concentrations of UCB-9721 (see numbers at the bottom of the bars in Figure 2B). Taken together, these results identify UCB-9721 as a potent inhibitor of *T. gondii* class XIV myosin activity.

**Figure 2.**
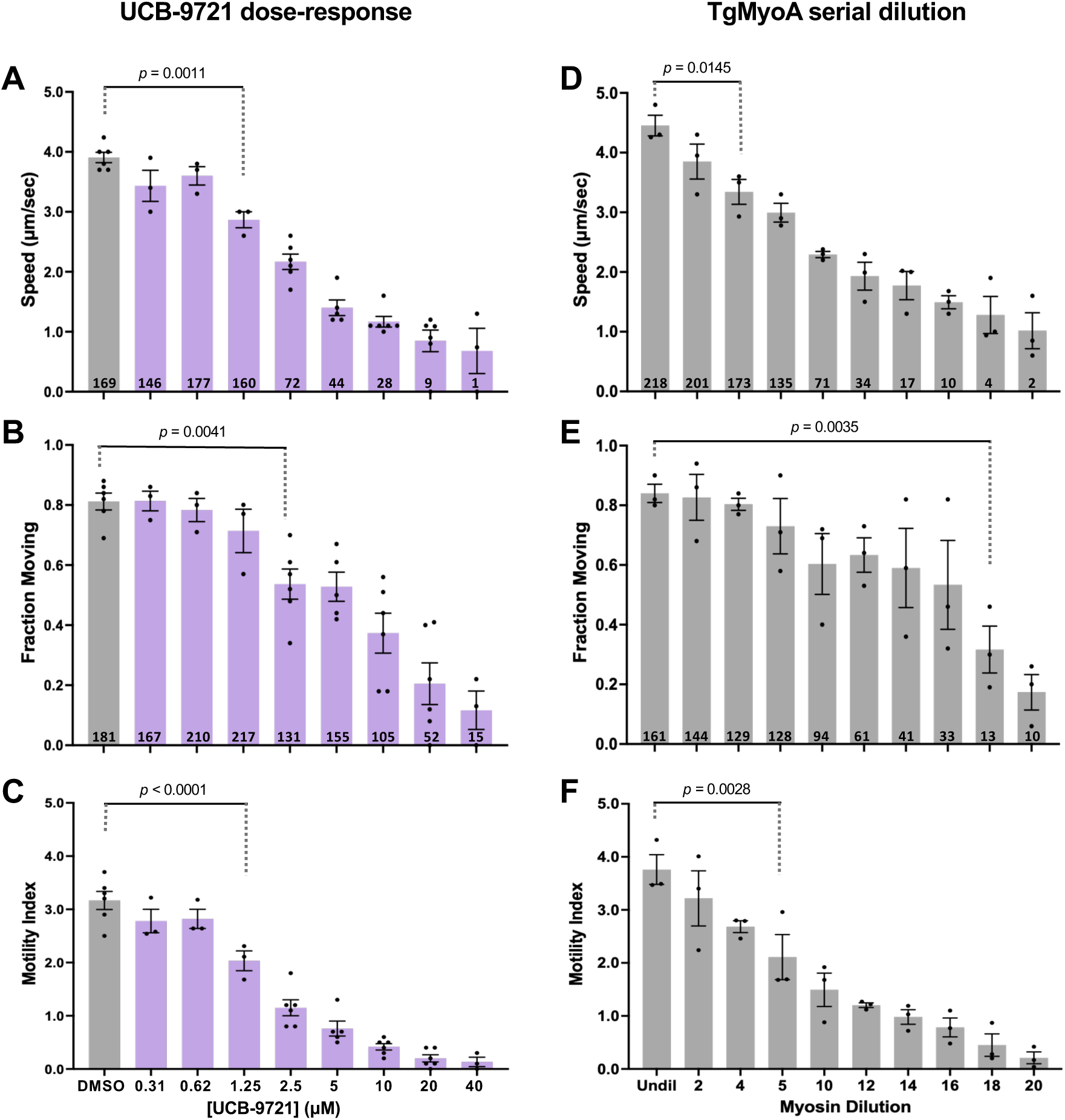
UCB-9721 inhibits *in vitro* motility driven by TgMyoA. **(A-C)**: *in vitro* motility assays, showing (A) the actin filament sliding speed, (B) the fraction of filaments bound to the coverslip that moved, and (C) filament motility index (speed x fraction moving) of TgMyoA treated with various concentrations of UCB-9721 (DMSO = vehicle only). Each data point represents the average from a single biological replicate consisting of 2 technical replicates, for a total of 3-6 biological replicates. Bars indicate the means of the biological replicates ± SEM. (**D-F):** *in vitro* motility assays in which the amount of TgMyoA added to the assay chamber was serially diluted from a starting concentration of 5.6 μg/ml (the concentration used in panels A-C; Undil). Each data point represents the average from a single biological replicate consisting of 2 technical replicates, for a total of 3 biological replicates. Bars indicate the means of the biological replicates ± SEM. The numbers at the bottom of the bars in Panels A and D indicate the average number of tracks, *i.e.*, moving filament trajectories, that were used to calculate speed in each of the biological replicates. The numbers at the bottom of Panels B and E indicate the average total number of filaments (moving plus unmoving) bound to the coverslip in each of the biological replicates. Compound treatments (A-C) or myosin dilution (D-F) were compared to DMSO or undiluted myosin, respectively, by one-way ANOVA with Dunnett’s test for multiple comparisons. Only the lowest concentration of UCB-9721 (A-C) and the least dilute myosin (D-F) showing a statistically significant difference from DMSO or undiluted myosin, respectively, is indicated; all compound concentrations higher than this value or more dilute myosins also resulted in speeds, fraction moving, and motility indices significantly different from the control.

### UCB-9721 inhibits *T. gondii* motility and growth, with no toxicity to host cells

To determine if UCB-9721 inhibits TgMyoA-dependent processes in cell-based assays, we first tested the compound’s effect on parasite motility in a 3D extracellular matrix model. UCB-9721 inhibits the percentage of parasites moving in a dose-dependent manner with an IC_50_ of 0.139 µM (95% C.I. = 0.075-0.247 µM; Figure 3A, B), a 44-fold decrease from the previously reported IC_50_ of KNX-002 in this assay (6.2 µM; ref. (36)).

**Figure 3:**
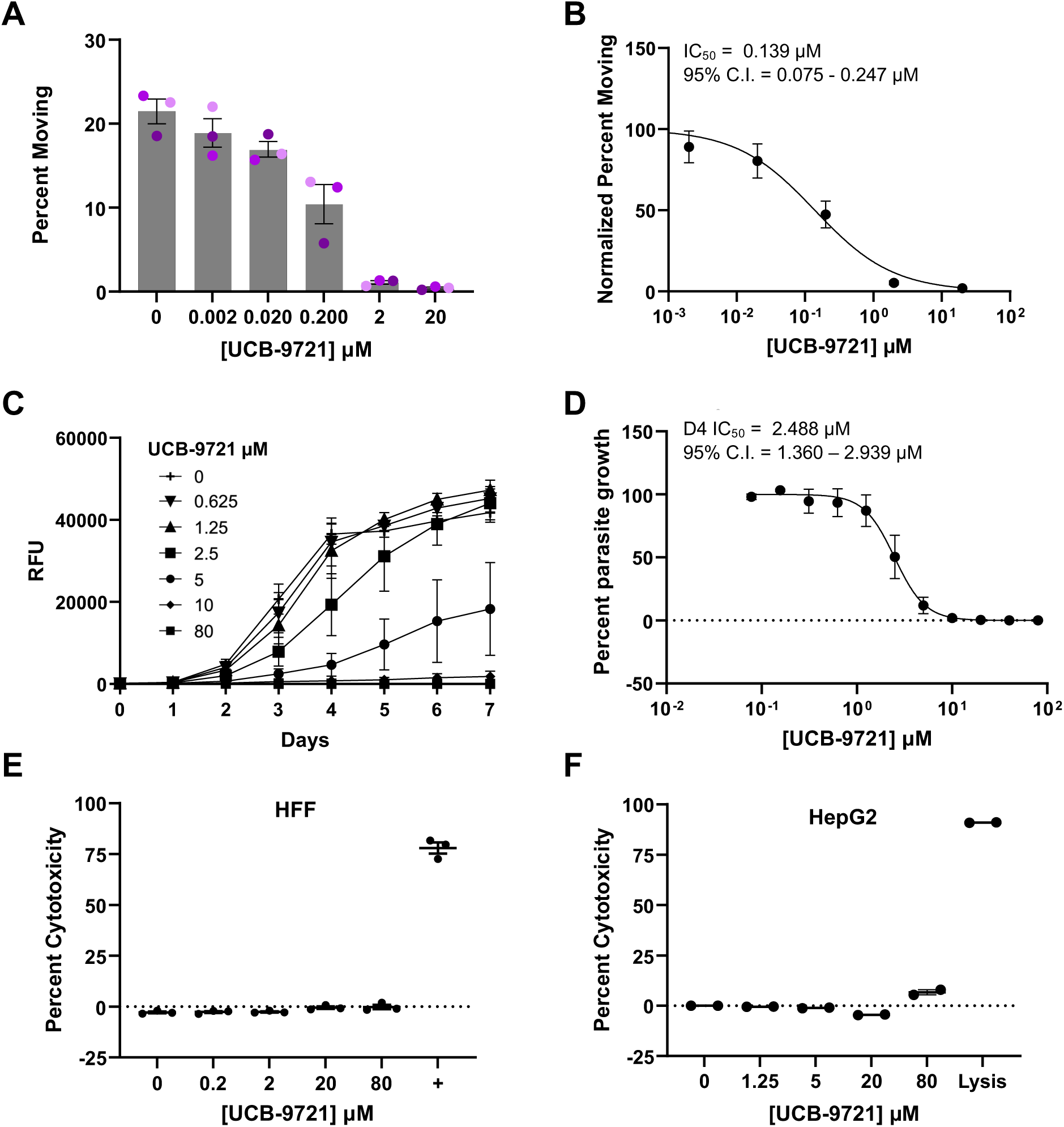
UCB-9721 inhibits 3D motility and growth of *T. gondii* parasites, with no toxicity to host cells. **(A)** Percentage of *T. gondii* tachyzoites moving in a 90-sec 3D motility assay in the presence of different concentrations of UCB-9721. Each data point represents a single biological replicate consisting of 2 technical replicates for a total of three biological replicates; bars indicate the means of the biological replicates ± SEM. **(B)** The data shown in Panel A were normalized to the DMSO control (0 µM) and used to calculate the IC_50_ of UCB-9721 for parasite motility (IC_50_ = 0.139 µM, 95% C.I. = 0.075 - 0.247 µM). **(C)** Growth curves of tdTomato-expressing parasites in the presence of different concentrations of UCB-9721. Fluorescence was measured daily; RFU = relative fluorescence units. The data shown are the mean ± SEM of 4 biological replicates, each consisting of three technical replicates of selected curves from a 12-point compound dilution series. **(D)** The data shown in Panel C on day 4 post infection (D4), normalized to the DMSO control (0 µM) and total inhibition (80 µM), were used to calculate the IC_50_ of UCB-9721 for parasite growth (IC_50_ = 2.488 µM, 95% C.I. = 1.360-2.939 µM); error bars indicate SEM. **(E, F)** Cytotoxicity assay results for human foreskin fibroblasts (HFF; Panel E) and human hepatocytes (HepG2; Panel F) exposed to the indicated concentrations of UCB-9721. The data shown are the mean ± SEM of 2-3 biological replicates, each consisting of 3 technical replicates. The positive control in E, denoted by +, is a known toxic compound, FT093. The positive control in F, “Lysis”, indicates treatment of the parasites with lysis buffer (see Experimental Procedures for details).

As TgMyoA also plays an important role in *T. gondii* growth in culture (32, 33), we tested whether UCB-9721 inhibits parasite growth. UCB-9721 inhibited the growth of *T. gondii* in human foreskin fibroblasts (HFFs) with an IC_50_ of 2.488 µM (95% C.I. = 1.360-2.939 µM, Figures 3C) 3.2-fold less than the IC_50_ of KNX-002 in the same growth assay (8.056 µM, 95% C.I. 6.508 – 9.934 µM; Figure S2). Importantly, UCB-9721 showed no detectable toxicity at the highest concentrations tested to either human foreskin fibroblasts (HFFs), the cell line in which the parasites are cultured (Figure 3E), or human hepatocytes (HepG2), a sensitive cell line commonly used to test cytotoxicity of xenobiotics (Figure 3F; ref. (39)).

### UCB-9721 inhibits the growth and motility of other apicomplexan parasites

Since MyoA and its associated light chains are highly conserved across the apicomplexan phylum (29), we also tested whether UCB-9721 could affect the motility or growth of other apicomplexan parasites, namely *Plasmodium falciparum*, *Cryptosporidium parvum*, and *Babesia duncani*. *P. falciparum* MyoA (PfMyoA) is thought to power the motility of multiple life stages of the malaria parasite and to be particularly important in sporozoites, the highly motile stage initially inoculated into a human host from *Anopheles* mosquitoes (17, 40). UCB-9721 inhibited *P. falciparum* sporozoite motility in a dose-dependent manner (IC_50_ = 1.037 µM [95% C.I. = 0.598-1.838 µM]), as shown by a significant decrease in the quantified pixel area occupied by parasite surface proteins deposited onto the glass coverslip in 2D parasite motility assays (Figure 4A, B). UCB-9721 also inhibited the growth of *B. duncani* in erythrocytes (IC_50_ = 4.049 µM [95% C.I. = 3.741-4.378 µM]; Figure 4C) and was found to be a particularly potent inhibitor of *C. parvum* growth in human ileocecal adenocarcinoma (HCT-8) cells (IC_50_ = 0.438 µM [95% C.I. = 0.341- 0.564 µM]; Figure 4D).

**Figure 4.**
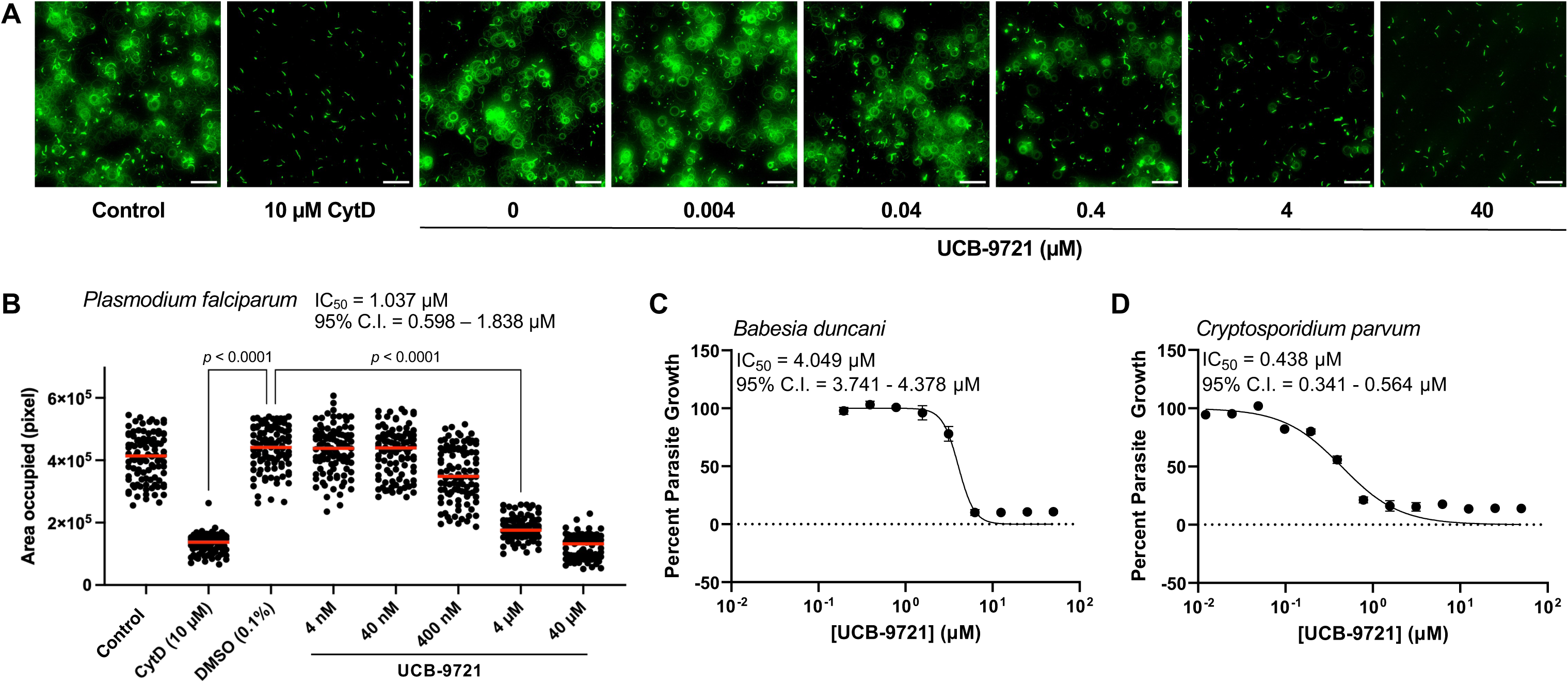
UCB-9721 affects the motility and growth of other Apicomplexan parasites. **(A)** Images of the trails left behind as *Plasmodium falciparum* sporozoites move on glass coverslips in the presence of the indicated concentrations of UCB-9721. Medium with DMSO and cytochalasin D (CytD) served as negative and positive controls, respectively. The trails were visualized by immunofluorescence. Scale bar = 50 µm. **(B)** The pixel area occupied by the fluorescent signal was quantified for 25 images per well (2 wells for each condition); red bars indicate the median of pooled data from two biological replicates (IC_50_ = 1.037 µM, 95% C.I. = 0.598-1.838 µM). All conditions were compared to each other by Kruskal-Wallis test followed by Dunn’s test. **(C)** Dose response data from growth assays of *Babesia duncani* in A+ red blood cells incubated in the presence of different concentrations of UCB-9721. The data shown are the mean ± SD of 2 biological replicates (IC_50_ = 4.054 µM, 95% C.I. = 3.278-4.408 µM). **(D)** Dose response data from growth assays of *Cryptosporidium parvum* in human ileocecal adenocarcinoma cells (HCT8) cells in the presence of different concentrations of UCB-9721. The data shown are the mean ± SD of 2 biological replicates (IC_50_ = 0.4351 µM, 95% C.I. = 0.2557-0.6188 µM).

### Pharmacokinetic properties of UCB-9721

To test whether UCB-9721 could be used to study MyoA function *in vivo*, we analyzed the compound’s pharmacokinetic (PK) properties in mice dosed intraperitoneally (i.p.) with 20 mg/kg (Figure S3). While the compound showed two desirable features for a drug targeting acute or reactivating *Toxoplasma* infections, *i.e.*, good *in vivo* bioavailability (99% at 20mg/kg; Figure S3A) and intrinsic membrane permeability (5.0 x 10^-6^ cm/sec with 92% recovery; Figure S3D), its very short plasma half-life (14-21 minutes) as a result of rapid clearance (3740 ml/hr/kg; Figure S3A) is a major liability for *in vivo* studies. Compound levels within the plasma dropped below the parasite growth assay IC_50_ in less than 90 minutes following i.p. injection (Figure S3B, right-hand graph) and modeling suggested that even increasing the dose 5-fold would be insufficient to overcome the high clearance rate and achieve a reasonable dosing regimen relative to the growth assay IC_50_ (Figure S3C). A second major liability is that penetration of the compound into the brain – the key target tissue for studying and/or treating reactivated infections – was extremely low (Figure S3A, B). Thus, while UCB-9721 will be a useful chemical probe for studying MyoA function in biochemical and cell-based assays, further optimization of its PK properties will be required for studies of TgMyoA function in animal models of toxoplasmosis. To determine which structural features of UCB-9721 are key to its activity and which can be modified in future efforts to improve its PK properties, we next used *in silico* docking and molecular dynamic (MD) simulation to explore the binding mode of UCB-9721 to TgMyoA.

### *in silico* docking reveals a potential rationale for the increased potency of UCB-9721 compared to KNX-002

Although UCB-9721 and KNX-002 were discovered in independent screens of different compound collections, they share a conserved pyrazole-based scaffold and differ only in replacement of the thiophene and cyclopropyl groups of KNX-002 by an ethoxyphenyl moiety and a sulfonamide linker, respectively, in UCB-9721 (Figure 5 A, B). Given the structural similarity between the two compounds, it seemed likely that they occupy the same binding pocket within the TgMyoA motor domain. The KNX-002 binding pocket was originally identified via X-ray crystallographic analysis of KNX-002 bound to PfMyoA (41), which showed that the compound binds to the “inner cleft” of PfMyoA close to the nucleotide binding pocket in the post-rigor state. In comparative molecular docking analyses, UCB-9721 and KNX-002 adopted highly similar binding poses in complex with post-rigor TgMyoA, in which the imidazole ring NH acted as a hydrogen-bond donor to a water molecule coordinated to D471 (PfMyoA D476). Additionally, the methoxyphenyl moiety of both inhibitors was predicted to occupy the same hydrophobic cavity, which is lined by residues L273, E275, E484, F487, and I643 (PfMyoA L271, E273, E482, F485, and I641). The root-mean-square deviation (RMSD), which is used as a measure of the similarity in structure between UCB-9721 and KNX-002 bound to TgMyoA and calculated using the heavy atoms of the maximum common substructure shared by the two inhibitors, was 0.27 Å. This low RMSD value indicates that the two compounds adopt highly similar binding orientations and likely occupy the same binding pocket within TgMyoA (Figure S4).

**Figure 5:**
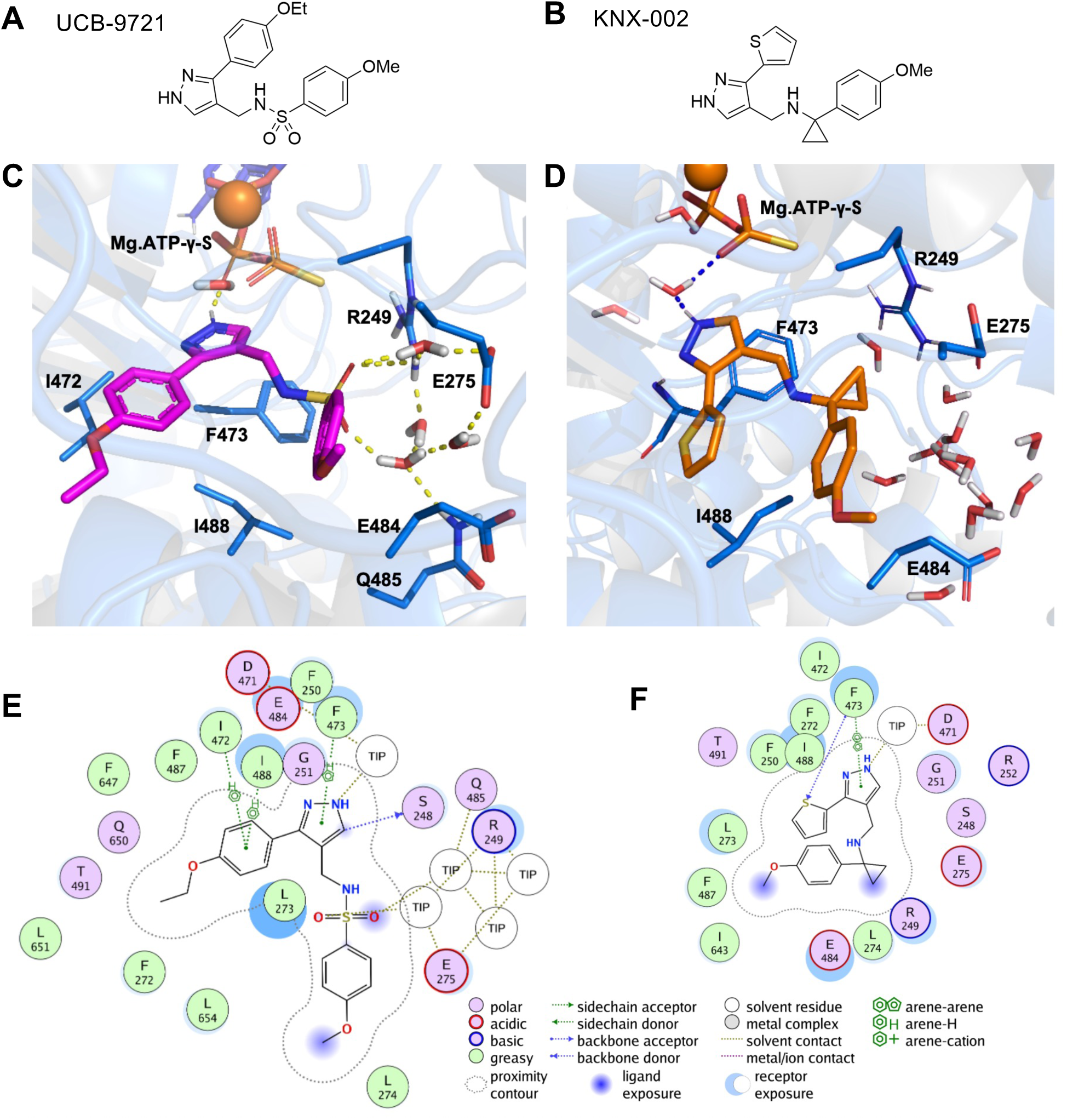
Structural analysis of compound binding modes in TgMyoA. **(A, B)** Chemical structures of UCB-9721 and KNX-002. **(C, D)** Binding poses were predicted by *in silico* docking and equilibrated using explicit-solvent molecular dynamics (MD) simulation. The protein is represented as ribbons and colored blue. Key residue side chains, ligands, and water molecules are depicted as sticks (H, white; O, red; N, blue; S, yellow). Hydrogen bonds and other polar interactions are indicated by yellow dashed lines with distances listed in Å. (C) Three-dimensional binding pose of UCB-9721 (magenta/purple) in the TgMyoA active site. The sulfonamide (–SO₂) group is stabilized by key hydrogen bonding interactions with water molecules coordinated by R249 and E275. (D) Three-dimensional binding pose of KNX-002 (orange) within the TgMyoA active site. **(E,F)** Two-dimensional ligand interaction diagrams for UCB-9721 (E), and KNX-002 (F) were derived from Molecular Operating Environment (MOE). TIP indicates the water molecules. Residues are colored by chemical type, and interaction types are defined in the legend below.

Despite the predicted binding of UCB-9721 in the KNX-002 binding pocket and the high structural similarity between the two compounds, UCB-9721 is almost 40-fold more potent than KNX-002 in the TgMyoA actin-activated ATPase assay and 45-fold more potent in the parasite 3D motility assay. This large difference in potency likely derives from unique interactions of TgMyoA with either the *p-*ethoxyphenyl ring and/or the sulfonamide group of UCB-9721, which are not present in KNX-002. To explore the molecular basis for the enhanced potency of UCB-9721, we carried out molecular dynamics (MD) simulations of KNX002 and UCB-9721 docked to a homology model of TgMyoA in the post-rigor state, in an explicit solvent environment. As described below, the resulting trajectories revealed interactions between the sulfonamide group of UCB-9721 and two TgMyoA residues, R249 and E275, as likely key contributors to the increased potency of UCB-9721.

The distance between the nearest NH hydrogen atom of the guanidinium group of R249 in TgMyoA and one of the sulfonamide oxygens of UCB-9721 was predicted to be 3.2 Å, compared to 3.9 Å between the same R249 hydrogen atom and a carbon atom of the amino-cyclopropyl group in KNX-002 (Figure 5C, D). The MD simulation trajectory revealed direct and water-mediated hydrogen bonds between the R249 guanidinium group and the sulfonamide oxygens (Figure S5A). Results from the trajectory analysis showed that a R249-N–H···O=S hydrogen bond was present in 81.2% of the sampled frames, predicting that R249 forms a persistent H-bond contact with a sulfonamide oxygen throughout the simulation (Figure S5A). In addition, a second N–H of R249 donated a hydrogen bond to a nearby water molecule in 95.8% of frames, while that same bridging water formed a hydrogen bond with a sulfonamide oxygen in 70.3% of frames; this bridged R249···H₂O···sulfonamide interaction was observed in 68.5% of the simulation (Figure S5A). Based on these observations, we conclude that R249 strongly interacts through direct and water-mediated hydrogen bonds with the sulfonamide moiety of UCB-9721.

Similar trajectory analysis was performed to monitor the interactions of E275 and the sulfonamide group in UCB-9721. The analysis showed that the average distance between a bridging water molecule and E275 or this water molecule and the sulfonamide group were 2.4 Å and 2.9 Å, respectively (Figures 5E and S5B). Hydrogen bond analysis further revealed that the bridging water molecule formed hydrogen bonds with either E275 or the sulfonamide group for more than 50% of the simulation time. Together, these results suggest that, in addition to R249, E275 contributes to the stabilization of the UCB-9721 sulfonamide moiety in the binding pocket. The MD simulation also showed that residue Q485 is engaged in the water-mediated interaction network with UCB-9721 within the compound-binding pocket (Figure 5E). These water molecules interact more favorably with the sulfonamide group in UCB-9721 than with the cyclopropyl group of KNX-002 (Figures 5C-E and S5A, B).

Next, we tested experimentally whether these predicted interactions are responsible for the increased potency of UCB-9721 using two complementary approaches: (a) by determining whether mutations introduced into TgMyoA at R249 or E275 reduce its sensitivity to UCB-9721; and (b) by determining which specific substituent(s) on UCB-9721 are responsible for its increased potency through a directed structure-activity relationship (SAR) study.

### Mutational analysis of TgMyoA confirms the importance of R249 for TgMyoA function and is consistent with a role for E275 in UCB-9721 binding

Based on our docking analyses, we predicted that mutating R249 of TgMyoA would lead to reduced sensitivity to UCB-9721. However, the homologous amino acid in *Dictyostelium* myosin II (ddMyoII) contributes to an essential salt-bridge interaction that stabilizes the ATP binding pocket prior to ATP hydrolysis, and a mutation from arginine to alanine at this position rendered the ddMyoII motor non-functional (42). Not surprisingly, therefore, when we introduced an R249A mutation into recombinant TgMyoA (TgMyoA^R249A^), the recombinant protein was completely inactive in the *in vitro* motility assay (Videos S2A, B). Similar levels of soluble TgMyoA^R249A^ and wild-type TgMyoA (TgMyoA^WT^) were produced in our expression system and purified TgMyoA^R249A^ and TgMyoA^WT^ bound similar amounts of actin in the *in vitro* motility assay (Videos S2A, B), suggesting that the lack of activity of TgMyoA^R249A^ in the assay is likely due to a functional defect in the motor rather than to a change in the integrity or stability of the mutant protein.

TgMyoA with an E275A mutation (TgMyoA^E275A^) retained partial function in the *in vitro* motility assay, translocating actin filaments at ∼25% the speed of the wild-type motor (compare DMSO samples in Figure 6A, D and Videos S2A, C). This partial function enabled us to test the prediction that E275 plays a role in the binding of UCB-9721. The addition of UCB-9721 to wild-type TgMyoA (TgMyoA^WT^) inhibited the speed of the moving filaments in a dose-dependent manner (Figures 2 and 6A). In contrast, the slower filament speeds generated by TgMyoA^E275A^ were completely insensitive to addition of up to 40 μM UCB-9721 (Figure 6D). The compound reduced the fraction of filaments moved by both TgMyoA^WT^ and TgMyoA^E275A^ in a dose-dependent manner (Figure 6B, E). However, when speed and fraction moving were combined into the filament motility index, TgMyoA^WT^ was clearly more sensitive to UCB-9721 treatment than TgMyoA^E275A^ (Figure 6C, F): 2.5 μM UCB-9721 reduced the motility index of TgMyoA^WT^ by more than 50%, whereas at least 20 μM UCB-9721 was needed to achieve a 50% reduction in the motility index of TgMyoA^E275A^. These data are consistent with the modeling predictions for the role of E275 in the binding of UCB-9721 to TgMyoA.

**Figure 6:**
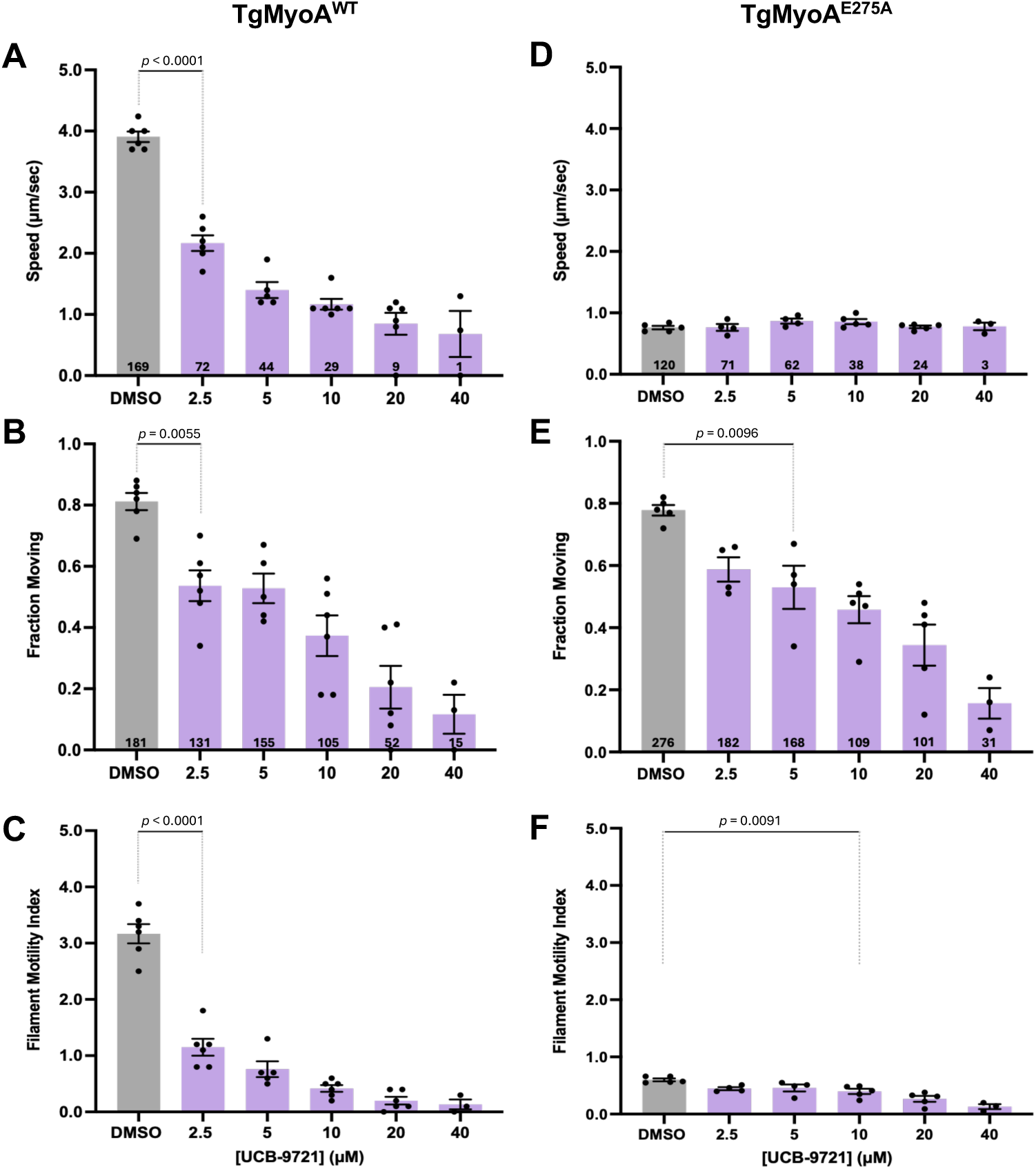
TgMyoA^E275^ is less sensitive to UCB-9721 than TgMyoA^WT^. *in vitro* motility assays, showing the actin sliding speed (top panels), fraction of actin filaments moving (middle panels), and corresponding filament motility index (fraction moving × mean speed; bottom panels) of TgMyoA^WT^ (panels A-C) and TgMyoA^E275^ (D-F) treated with various concentrations of UCB-9721, as indicated (DMSO = vehicle only). Each data point represents the average from a single biological replicate consisting of 2 technical replicates, for a total of 3-6 biological replicates; bars indicate the means of the biological replicates ± SEM. The TgMyoA^WT^ data in panels A-C are a subset of the same data shown in Figure 2A-C, selected here for direct comparison to the TgMyoA^E275^ data in panels D-F. Compound treatments were compared to DMSO vehicle by one-way ANOVA with Dunnett’s test for multiple comparisons. Only the lowest concentration showing a statistically significant difference from vehicle is indicated; all UCB-9721 concentrations higher than this value yield results that are also significantly different from the control. Numbers at the bottom of the bars indicate the average number of trajectories (A, D) or filaments (B, E) measured at each compound concentration.

### Directed SAR identifies the sulfonamide group of UCB-9721 as critical for its potency

In parallel with the mutational studies described above, structure-activity relationship analysis was used to inform on key interactions mediating the binding of UCB-9721 to TgMyoA. Eleven structural analogs of UCB-9721 (seven from the original screening collection and four synthesized) were assayed for their effect on TgMyoA activity (see Table S1 for the full list; key compounds are highlighted in Table 1). Based on the compounds from the screening collection it was concluded that, in general, substitutions in the sulfonamide aryl group lead to decreased inhibition of actin-activated ATPase activity (UCB-82-823, UCB-82-815, UCB-82-817, UCB-82-825, UCB-82-821, Table S1). Functional groups at the *p-*position are tolerated better than *o-*and *m-* substitutions (compare UCB-82-821 *m*-CF_3_ and UCB-82-817 *p*-CF_3_ IC_50_s; also decreased activity of UCB-82-825 *o-*ethyl, Table S1). Two of the synthesized analogs provided additional insight into the nature of the pocket occupied by the *p-*substituted aryl ring bound to the pyrazole, as the removal of a -CH_2_ unit from the O-ethoxy substituent in UCB-9721 decreased its activity (compare VEST16 to UCB-9721, Table S1, Figure S6). This effect is even more pronounced in VEST17, in which -CH_2_ unit from the O-ethoxy substituent was removed and the sulfonamide was replaced with an amino cyclopropyl group (Table S1, Figure S6).

**Table 1:**
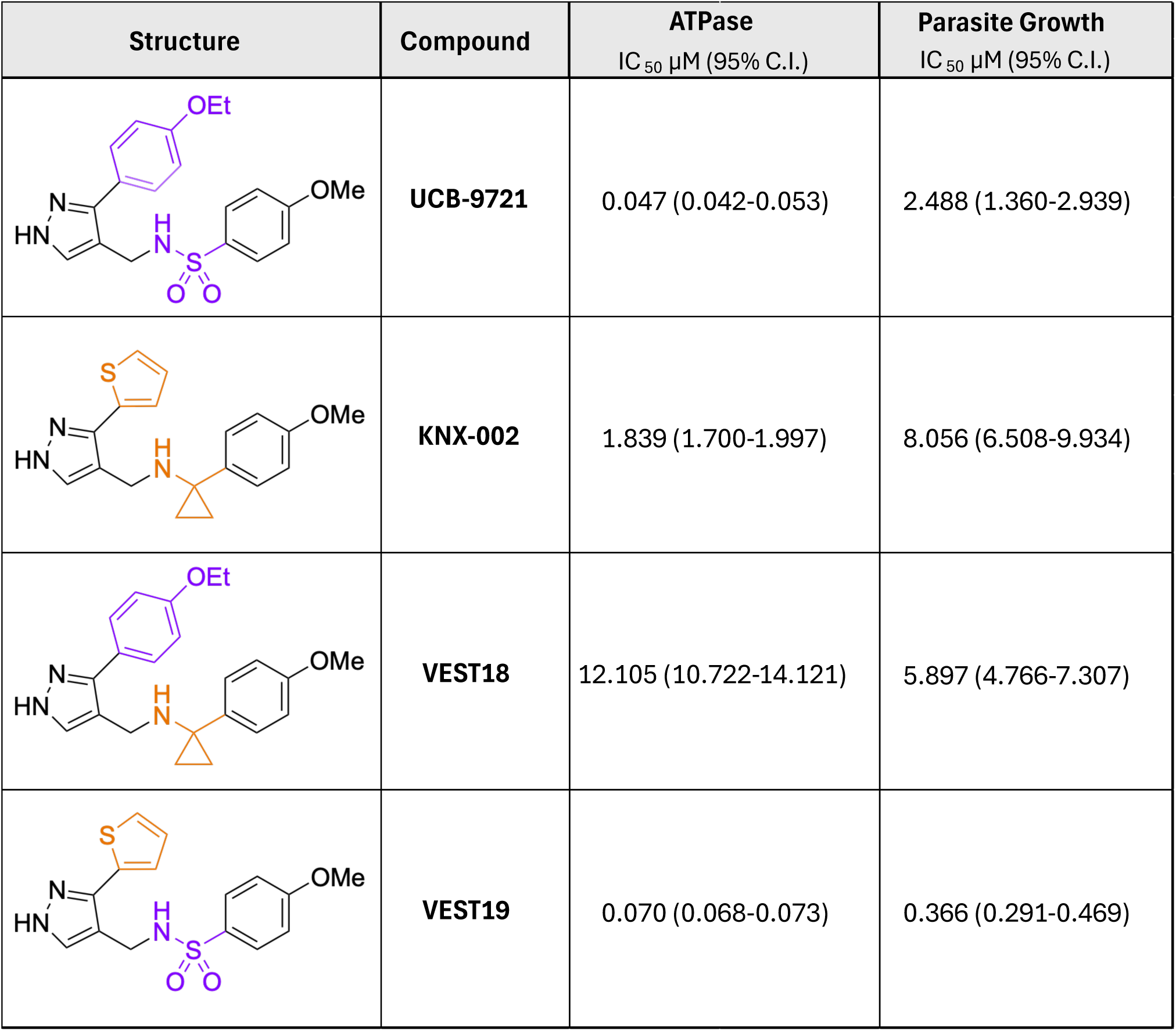
The increased potency of UCB-9721 compared to KNX-002 is due to the cyclopropyl to sulfonamide replacement. . Table shows chemical structures of compounds with colors denoting unique features of UCB-9721 (purple) and KNX-002 (orange). Also shown are each compound’s IC_50_ for actin-activated ATPase and parasite growth, with 95% confidence intervals.

| Structure | Compound | ATPase<br>IC<sub>50</sub> μM (95% C.I.) | Parasite Growth<br>IC<sub>50</sub> μM (95% C.I.) |
| --- | --- | --- | --- |
|  | UCB-9721 | 0.047 (0.042-0.053) | 2.488 (1.360-2.939) |
|  | KNX-002 | 1.839 (1.700-1.997) | 8.056 (6.508-9.934) |
|  | VEST18 | 12.105 (10.722-14.121) | 5.897 (4.766-7.307) |
|  | VEST19 | 0.070 (0.068-0.073) | 0.366 (0.291-0.469) |

The greatly reduced activity of VEST17 compared to VEST16 supported the importance of the sulfonamide group to the potency of UCB-9721. To determine which portion(s) of UCB-9721 are responsible for its increased potency compared to KNX-002, we synthesized two analogs of KNX-002: (a) VEST18, in which the thiophene ring of KNX-002 was replaced by the *p-*ethoxyphenyl ring of UCB-9721 and (b) VEST19, in which the amino-cyclopropyl group in KNX-002 was replaced by the sulfonamide of UCB-9721. A comparison of TgMyoA actin-activated ATPase activity and *T. gondii* growth in the presence of KNX-002 and VEST18 (Table 1) showed clearly that the increased potency of UCB-9721 is not due to its ethoxyphenyl ring. A comparison of actin-activated ATPase inhibition by KNX-002, UCB-9721, and VEST19 (Table 1) revealed that the increased potency of UCB-9721 can instead be attributed to the sulfonamide group, which is consistent with the modeling predictions presented in Figure 5. VEST19, which combines the sulfonamide of UCB-9721 with the thiophene ring of KNX-002, is a particularly interesting analog for two reasons. First, while the activity of VEST19 against TgMyoA actin-activated ATPase activity is comparable to that of UCB-9721, VEST19 is 7-fold more potent than UCB-9721 in the *T. gondii* growth assay (Table 1). VEST19 is therefore the most effective TgMyoA inhibitor we have thus far identified. Second, we recently learned that VEST19 is identical to KNX-115, a compound independently discovered by Trivedi, Spudich, and colleagues at Kainomyx, Inc., as a potent inhibitor of *P. falciparum* MyoA (ref. (43); discussed further below).

## Discussion

### UCB-9721 as a chemical probe for studying the motility of apicomplexan parasites

While a role for TgMyoA in apicomplexan parasite motility has been well established, significant mechanistic questions remain. These include: how the motor is organized within the parasite pellicle; where and how the force generated by the intracellular motor is transmitted to the surrounding substrate to generate forward motion (44); how motor activity is regulated to generate the periodic acceleration/deceleration and complex 3D trajectories observed both *in vitro* and *in vivo* (45); and whether the force-generating capacity of the motor changes depending on biological/environmental context (*e.g.,* migration along two dimensional endothelial surfaces vs invasion of host cells) (46–48). Small-molecule inhibitors provide a powerful approach for studying rapid and dynamic processes in parasite biology (49, 50), such as motility, because the timing of exposure to the compound can be stringently controlled and compensatory adaptations that can arise in response to genetic manipulation are avoided. Chemical probes exhibiting activity in one experimental system may also be used in cross-species studies, serving as a useful way to study less experimentally tractable organisms (in the case of apicomplexan parasites, this includes *Cryptosporidium* and *Babesia*). Finally, potent and specific inhibitors of proteins critical to the parasite’s lytic cycle, such as MyoA, can serve as the starting point for drug development.

In this study, we identified and characterized a potent new small-molecule inhibitor of TgMyoA – UCB-9721. This compound inhibits TgMyoA actin-activated ATPase activity, the ability of the recombinant motor to translocate actin filaments *in vitro*, and parasite motility and growth in cell-based assays. While UCB-9721 is structurally related to a previously identified inhibitor of TgMyoA (KNX-002), UCB-9721 was approximately 40-fold more potent in recombinant myosin- and parasite-based assays and significantly more potent in assays measuring growth within host cells. We used a combination of molecular docking, targeted TgMyoA mutagenesis, and directed SAR analysis to determine that the source of this increased potency is the sulfonamide group present in UCB-9721 but not in KNX-002. Finally, our demonstration that UCB-9721 inhibits the motility and growth of several other apicomplexan parasites in addition to *T. gondii* suggest it will be a valuable chemical probe for studying class XIV myosin function and motility in apicomplexan parasites in general.

### UCB-9721, KNX-002, KNX-115 and the compound-binding pocket in TgMyoA

In a 2022 preprint, Trivedi *et al* reported that a derivative of KNX-002 (KNX-115) was 20-fold more potent than KNX-002 at inhibiting recombinant PfMyoA actin-activated ATPase activity (51). A followup manuscript from this same group revealed that the structure of KNX-115 contains the same sulfonamide group we showed to be critical for UCB-9721’s improved potency (43). In fact, KNX-115 is identical to compound VEST19, which was synthesized in our SAR series by combining the thiophene ring from KNX-002 and the sulfonamide from UCB-9721 (Table 1). We found that the activity of VEST19/KNX-115 against TgMyoA actin-activated ATPase activity was comparable to that of UCB-9721, but VEST19/KNX-115 was several-fold more potent in the cell-based growth assay, suggesting that the combined thiophene ring and sulfonamide group in VEST19/KNX-115 confer improved host cell permeability or stability. VEST19/KNX-115 has been shown to trap *Plasmodium falciparum* MyoA in a state that binds weakly to actin (43, 51). Our observation that UCB-9721 decreased the number of actin filaments bound to the TgMyoA-coated coverslip in the *in vitro* motility assay (Figure 2B, numbers at base of bars) suggests that UCB-9721 similarly impacts binding affinity of TgMyoA for actin.

The modeling and mutagenesis studies presented here argue that UCB-9721 binds to TgMyoA within the homologous pocket that X-ray crystallography identified as the binding site of KNX-002 and VEST19/KNX-115 within post-rigor PfMyoA (41, 43, 51). This compound-binding pocket is adjacent to the ATP binding site, and while the amino acids within the pocket are largely conserved between apicomplexan MyoA homologs (91% similarity), they diverge significantly in mammalian myosins (60% similarity) (27) offering a potential explanation for the high degree of specificity this class of compounds shows for MyoA over mammalian myosins (36, 43, 51). The high degree of sequence identity/similarity seen in the binding pocket residues of the different apicomplexan MyoAs also suggests that the structure of TgMyoA in this region is important for class XIV myosin motor function. The TgMyoA^T130A^ mutation that we previously demonstrated provides partial resistance to KNX-002 is located in the transducer domain far away from the KNX-002 binding pocket, and the chemical mutagenesis strategy employed in that study to generate resistance to KNX-002 did not result in any viable parasites containing mutations within the compound-binding pocket (36). Similarly, when Trivedi *et al* subjected malaria parasites to weak selective pressure with VEST19/KNX-115, none of the resulting PfMyoA mutations that conferred drug resistance were located within the compound-binding site; all were allosteric mutations elsewhere in the motor domain (43). Combined, these results strongly suggest that the residues within the compound-binding pocket are important for class XIV myosin motor function. The lack of motor activity of recombinant TgMyoA^R249A^ and the significantly reduced activity of TgMyoA^E275A^ (Figure 6) are consistent with the functional importance of residues within this compound-binding pocket.

As in our studies with UCB-9721, Trivedi *et al* found that VEST19/KNX-115 inhibits the growth of multiple apicomplexan parasites (43). This observation, together with mapping of the same compound binding site and establishment of the importance of the sulfonamide group highlight both the complementary nature of these two independent studies and the broad-based value of this class of compounds as chemical probes for studying MyoA function in apicomplexan parasites.

### Prospects for *in vivo* applications

While there is no universally accepted target product profile for drugs to treat acute or reactivating *T. gondii* infections, key PK properties would include a plasma half-life long enough to support a reasonable dosing regimen, and good penetration into the tissues where reactivation takes place (muscle, eye, and especially the brain). UCB-9721 currently falls short on both of these criteria, despite its excellent bioavailability and high intrinsic membrane permeability. Future medicinal chemistry efforts will therefore be directed towards improving the high clearance rates and poor tissue exposure of UCB-9721. The short plasma half-life of UCB-9721 may be due to glucuronidation on one of the nitrogens of the pyrazole ring (52), as has been observed with other compounds containing a similar substituent (*e.g.*, (53)). However, given the predicted importance of the pyrazole ring for binding to MyoA (Figure 5C, E and refs. (41, 43)), the compound will most likely not tolerate substitution on the pyrazole nitrogens. It may therefore be necessary to design a prodrug that releases the active compound exclusively in the parasite. Alternatively, UCB-9721 and its active analogs could be used to design a ligand-based 3D pharmacophore, which would then be screened *in silico* for predicted inhibitors lacking the pyrazole group, an approach with which we (BN, SK) have extensive experience.

While the suboptimal PK properties of UCB-9721 in mice were disappointing, different PK characteristics will be necessary for drugs to treat the diseases caused by different apicomplexan parasites: toxoplasmosis is primarily a disease of the muscle, retina, and brain; malaria is a disease of the blood; and cryptosporidiosis is an infection of the small intestinal epithelium. Without further development, the PK data presented here suggest that UCB-9721 will be of limited use for treating either *Toxoplasma* or *Plasmodium* infections *in vivo*. However, the compound’s solubility and permeability profile (Figure S3D) is comparable to that of other compounds that are efficacious against cryptosporidiosis *in vivo* (54), where slow absorption as a result of suboptimal solubility or permeability is thought to maximize the concentration of compound in the intestinal epithelium. Future studies will therefore evaluate the efficacy of UCB-9721 in a mouse model of cryptosporidiosis. It will also be of interest to determine whether VEST19/KNX-115 has better plasma or tissue exposure than UCB-9721 due to the presence of the thiophene ring in VEST19, which our data show confers improved activity in cell-based *Toxoplasma* growth assays (Table 1, Figure S6) – although it should be noted that thiophene-containing compounds are frequently associated with unacceptable levels of toxicity *in vivo* (55). Finally, there is currently intense interest in using drug-impregnated bed nets to target malaria parasites within the mosquito vector as a means of reducing malaria transmission (56). We have shown that UCB-9721 is a potent inhibitor of the motility of insect salivary gland sporozoites; whether this class of compounds could be delivered via treated bed nets and achieve a sufficient half-life and tissue distribution within the mosquito to impact transmission will require further study.

### Summary

The data presented here, in our previous study (36) and in the work of others (41, 43, 51) collectively demonstrate that UCB-9721 and the related compound VEST19/KNX-115 are promising new chemical probes for studying the function of class XIV myosins *in vitro* and in cell-based assays. Our previous study demonstrated that TgMyoA is a druggable target *in vivo* (36), and the data presented here will help guide further development of this class of compounds to explore their value as novel antiparasitic compounds for use in treating infections *in vivo*.

## Experimental Procedures

### High-throughput Screen

UCB-9721 was identified through a high-throughput screen of 50,422 compounds at the UC Berkeley Drug Discovery Center within the Center for Neglected and Emerging Diseases at the University of California Berkeley. The screen was designed to identify inhibitors of the actin-activated ATPase activity of TgMyoA as previously described using an enzymatically coupled steady state assay coupling MyoA ATPase function to NADH oxidation (36). Hits were also screened for activity against the coupling system using an unrelated ATPase, hexokinase/Glucose.

### Compound synthesis and use

Compounds were synthesized as described in Supporting Information and stored at -20°C in DMSO in the dark until use. In all experiments with compounds, an equal final amount of DMSO (vehicle) was added to each sample within the experiment; final DMSO concentrations in *T. gondii* growth and 3D motility assays were 0.2% v/v.

### Cell and Parasite Culture

Human foreskin fibroblasts (HFFs; SCRC-1041, ATCC, Manassas, VA) were grown to confluence in Dulbecco’s Modified Eagle’s Medium (DMEM) (Life Technologies, Carlsbad, CA) containing 10% v/v heat-inactivated fetal bovine serum (FBS) (Life Technologies), 10 mM HEPES (pH 7), and 100 units/ml penicillin and 100 μg/ml streptomycin, as previously described (57). Once HFFs were confluent, the medium was changed to DMEM with 10 mM HEPES (pH 7), 100 units/ml penicillin, 100 μg/ml streptomycin, and 5% v/v FBS. Parasites were propagated by serial passage in confluent HFFs.

HepG2 cells were provided by Tom Jetton (UVM) and passaged in DMEM (Life Technologies) containing 10% v/v heat-inactivated fetal bovine serum (FBS) (Life Technologies), 10 mM HEPES (pH 7), and 100 units/ml penicillin and 100 μg/ml streptomycin. These cells were filtered post trypsinization through a 40 µm filter (Thermo Fisher Scientific, Waltham, MA, USA) during each passage.

### *In vitro* motility assays

Expression and purification of recombinant TgMyoA and determination of filament speeds in *in vitro* motility assays were performed as previously described (36). To determine the fraction of actin filaments moving in a given video recording, both the total number of fluorescent actin filaments in the first frame of the video (T_f_) and the total number of filaments exhibiting directed motion that persisted for at least 5 μm (T_m_) were counted manually using IMARIS 10.2.0 (Oxford Instruments Andor, Belfast, Northern Ireland). Fraction moving for that video was calculated as T_m_ /T_f_.

### Parasite Motility Assays

#### *T. gondii* 3D motility

Imaging chambers were prepared as previously described (45). Parasites were isolated by scraping, syringe releasing, and filtering through a 3-μM Whatman Nuclepore filter (Millipore Sigma, St. Louis MO). Parasites were then pelleted at 1,000 *x g* for 2 min and resuspended in live cell imaging solution (LCIS: Ringer’s solution with glucose to 20 mM: 155 mM NaCl, 2 mM KCl, 2 mM CaCl_2_, 1 mM MgCl_2_, 3 mM NaH_2_PO_4_, 10 mM HEPES, 20 mM glucose, pH 7.2) with 0.5 mg/mL Hoechst 33342. Parasites, LCIS containing compound or vehicle control, and Matrigel were mixed at a ratio of 1:3:3 and pipetted into a flow cell. Concentrations of compound listed are final concentrations within the flow cell with a constant DMSO concentration of (0.2% v/v). Flow cells were then incubated for 7 min at 27°C and 3 min at 35°C. Imaging was carried out at at 35°C on a Nikon Eclipse TE300 epifluorescence microscope (Nikon Instruments, Melville, New York, USA) using a 20× PlanApo λ objective and a NanoScanZ piezo Z stage insert (Prior Scientific, Rockland, Massachusetts, USA). 1,024 pixel × 384 pixel images were captured using an iXON Life 888 EMCCD camera (Andor Technology, Belfast, Northern Ireland) driven by NIS Elements software v.5.11, including the Illumination Sequence acquisition module (Nikon Instruments, Melville NY), and excited using a pE-4000 LED illumination system (CoolLED, Andover, England). Images were captured for 80 s. The camera was set to trigger mode, no binning, readout speed of 30 MHz, conversion gain of 3.8×, and an EM gain of 300.

All 3D motility assays captured image stacks consisting of 41 z-slices, captured 1 μm apart with a 16 ms exposure time. The data were analyzed using Imaris ×64 v. 9.2.0 (Bitplane AG, Zurich, Switzerland). The ImarisTrack module tracked the parasite nuclei using an estimated spot volume of 3.0 × 3.0 × 6.0 μm. A maximum distance of 6.0 μm and maximum gap size of 2 were applied to the tracking algorithm. A track duration filter of 10 s was used to avoid tracking artifacts and a 2 μm displacement filter was used to eliminate parasites moving by Brownian motion (45).

#### *Plasmodium falciparum* sporozoite 2D motility

*P. falciparum* NF54 blood stage cultures were propagated in O^+^ erythrocytes at a 4% hematocrit in RPMI 1640 supplemented with 2.1mM L-glutamine, 25mM HEPES, 0.72 mM hypoxanthine, 0.21% w/v sodium bicarbonate, and 10% v/v heat inactivated human serum (Interstate Blood Bank) and maintained at 37°C in a glass candle jar. Gametocyte cultures were initiated at 0.5% asexual blood stage parasitemia and maintained by changing the media daily for 17 days without the addition of fresh blood to promote gametocytogenesis as previously described (58). Stage V gametocytes at 0.3% gametocytemia in 40% hematocrit were fed to *Anopheles stephensi* mosquitoes using a glass membrane feeder for 30 min. Infected mosquitoes were maintained at 25°C and 80% humidity and were provided with 10% w/v sucrose solution for 19 days, after which mosquito salivary glands were dissected in Hanks’ Balanced Salt Solution (HBSS) and manually homogenized to release sporozoites.

Motility assays were performed as previously described (59). Briefly,15,000 freshly dissected *P. falciparum* salivary gland sporozoites in 50 µl of HBSS/BSA pH 7.4 were mixed 1:1 with UCB-9721 in HBSS and pre-incubated for 30 minutes at 20°C and then transferred to a 96 well glass bottom plate (Greiner, Monroe NC) coated with 5 µg/ml of mAb 2A10 which is specific for the PfCSP repeat region. The plate was centrifuged for 3 minutes at 300 × g and incubated for 1 hour at 37°C. Wells were fixed in 4% paraformaldehyde in PBS, blocked with 1% BSA in PBS (pH 7.4) and stained with biotinylated mAb 2A10 in 1% BSA in PBS (pH 7.4) for 1 hour at room temperature, followed by detection with Alexa Fluor 488 streptavidin (Invitrogen, Carlsbad CA) diluted at 1:500 in PBS for 1 hour at room temperature. Samples were preserved in a glycerol / PBS solution (ratio, 9:1) at 4°C and imaging was performed on 25 positions per well (5 x 5,500 µm apart) by using ImageXpress Micro XLS Widefield high-content analysis system (Molecular Devices, San Jose CA) with 40X Plan fluor objective. Quantification of the area occupied by trails were determined by using Cell Profiler software (version 3.0.0) (60). All conditions were compared to each other by Kruskal-Wallis test followed by Dunn’s test. The IC_50_ of *P. falciparum* 2D motility was quantified from the means from each biological replicate normalized to the BSA negative control condition and the cytochalasin D (CytD) positive control using a non-linear regression in GraphPad Prism.

### Growth Assays

#### Toxoplasma gondii growth assays

Growth in the presence of compound(s) was measured as previously described (36), with the minor modifications described below. Tachyzoites expressing tdTomato (1 × 10^5^) were preincubated for 5 minutes with DMSO or serial dilutions of compound (with a DMSO concentration of 0.2% v/v in all samples), in DMEM with 10 mM HEPES (pH 7), 100 units/ml penicillin and 100 μg/ml streptomycin, and 1% v/v FBS. The parasites and medium were then added to individual wells of a black-sided, clear bottom 384-well plate containing a confluent monolayer of HFF (SCRC-1041) cells and incubated at 37°C. The fluorescent signal was quantified daily for 7 days (excitation 530/25 nm, emission 590/35 nm) on a Biotek synergy 2 plate reader (Agilent, Santa Clara, CA). The IC_50_ of growth in inhibitory compounds was quantified from the first day of growth plateau of the DMSO control condition after parasite inoculation (D4) using a non-linear regression in GraphPad Prism.

#### Babesia duncani growth assays

*Babesia* growth assays were performed as described in (61). Briefly, *B. duncani* were provided by the Mamoun lab at Yale University and cultured in complemented DMEM/F12 (cDMEM; 20% heat-inactivated FBS (Sigma-Aldrich), 1 x HT media supplement (Sigma-Aldrich), 1 x L-glutamine (Gibco, Grand Island NY), 1 x antibiotic-antimycotic (Gibco), 1 x gentamicin (Gibco)) in fresh A+ blood from the University of Vermont Medical Center Blood Blank. Fresh A+ blood was washed three times with PSG+ (Puck’s Saline Glucose Medium, 2% w/v D-glucose, 1 x antibiotic-antimycotic solution (Gibco)) and compounds to be tested were prepared in a 96-well plate by serial dilution in cDMEM. A 5% parasitemia, 10% hematocrit parasite stock was prepared in cDMEM. Equal volumes of parasite stock and compound were combined in 384-well plates (50 μL total with final 5% hematocrit, 2.5% parasitemia, 0.25% DMSO). Vehicle (DMSO) and the known inhibitor WR99210 (25 µM) (Sigma-Aldrich) were used as negative and positive controls for inhibition, and four technical replicates were included for each condition. Plates were incubated for 48 hours at 37°C in a modular incubator with 2% O_2_, 5% CO_2_, and 93% N_2_, after which 25 μL of PBS with 0.6% Triton-X and 22.5 μM propidium iodide (PI) were added. 24 hours later, fluorescence was measured (excitation 535, emission 615) using a Biotek synergy 2 plate reader (Agilent). GraphPad Prism version 10.0.2 (GraphPad Software, San Diego, CA) was used to generate dose-response curves and graphs.

#### Cryptosporidium parvum growth assays

*Cryptosporidium* growth assays were performed as previously described (62, 63). Briefly, *C. parvum* oocysts (Iowa strain) were obtained from Bunch Grass Farm (Deary, ID) and stored in phosphate-buffered saline (PBS) with penicillin and streptomycin at 4°C for up to 5 months prior to use. HCT-8 cells were grown to 90% confluence in 384-well plates and infected with oocysts that had been induced for excystation using 10 mM hydrochloric acid (10 min, 37°C) followed by 2 mM sodium taurocholate (Sigma-Aldrich) in PBS (10 min, 16°C). Different concentrations of compound were added to the HCT-8 cells 3 hrs after each well was inoculated with 5500 *C. parvum* oocysts. The plates were then incubated for 45 hr at 37°C. Wells were fixed and stained with fluorescein-labeled *Vicia villosa* lectin (Vector laboratories) to visualize parasitophorous vacuoles and Hoechst 33258 to image nuclei. Images were acquired using a Nikon Eclipse TE2000 epifluorescence microscope (Nikon, USA) with an automated stage programmed to acquire a 3 x 3 tiled image of the center of each well using an Exi Blue fluorescence microscopy camera (QImaging, Canada) and a 20x objective (NA 0.45). Nuclei and parasite counts were analyzed using custom macros on ImageJ. IC_50_ values were calculated using nonlinear regression analysis for dose-response (four parameters) with a top constraint set to 100.

### Cell Toxicity Assays

#### CellTox Green

HepG2 cells were seeded at 1 × 10^5^ cells/mL at 50 µL/well in a 96-welll plate. Cells were 80% confluent at time of assay when the medium was replaced with DMEM containing 10% v/v FBS and serial dilutions of compound or vehicle while maintaining a consistent DMSO concentration (0.2% v/v). Cells were incubated with compound at 37°C for 72 h. Toxicity was measured using the CellTox Green Cytotoxicity Assay (Promega, Madison, Wisconsin, USA) and the included controls. Fluorescence was quantified after 72 hr on Biotek synergy 2 plate reader (Agilent).

#### CellTox 96-well non-radioactive assay

HFF cells were seeded at 1 × 10^5^ cells/mL in a 96-well plate. Once cells reached confluency, the medium was replaced with DMEM containing 1% v/v FBS and serial dilutions of compound or vehicle while maintaining a consistent DMSO concentration (0.2% v/v), to match the growth assay conditions. Cells were incubated with compound at 37°C for 72 h. Toxicity was measured through the CytoTox 96 Non-Radioactive Cytotoxicity Assay (Promega, Madison, Wisconsin, USA) using kit provided controls. Absorbance was quantified after 72 hr (A:490 nm) on Biotek synergy 2 plate reader (Agilent, Santa Clara, CA).

### Homology Modeling and Molecular Docking

The three-dimensional structure of *T. gondii* MyoA (PDB ID: 6DUE, ref. (27)) was homology modeled using the crystal structure of KNX-002-bound PfMyoA (PDB ID: 8CDQ, ref. (41)) as a template using Molecular Operating Environment (MOE) 2022.02 (Chemical Computing Group ULC, Montreal, Canada). The modeled structure was energy minimized, and ionizable residues were adjusted for pH 7. Mg²⁺-bound ATP and the water molecules involved in KNX-002 binding were included in the modeled structure. UCB-9721 and KNX-002 were docked to the binding cavity lined by residues E275, R249, E484, S248, F487, F272, G251, T491, L273, F473 and I472 using GOLD docking software (64). The ligands were energy minimized prior to docking using AMBER force-field parameters (ff10) (65). For each ligand, the 30 best binding poses were reviewed and the pose with the highest docking score was selected for further analysis.

### Molecular Dynamics Simulation

The docked complexes were placed in a water box with dimensions 126 × 126 × 126 Å³ using the CHARMM-GUI (66). The protein complex was neutralized with 0.15 M KCl. Molecular dynamics (MD) simulations were performed using NAMD software (67). The complex was equilibrated for 1 ns with a 2-fs timestep while restraining the ligand, protein backbone atoms, ATP, Mg²⁺, and the water molecules bound to the ligand at 303.15 K. These restraints were subsequently removed for the production run, which was continued for an additional 20 ns with a 2-fs timestep. The CHARMM36m (68) force-field was used for the protein, and the TIP3P (69) model was used for water molecules. The forcefield parameters for the ligands were generated using CHARMM General Force Field (CGenFF) program (70). All MD simulations were carried out in the NPT ensemble using the Hoover thermostat (71).

### Pharmacokinetics

CBA/J mouse, 18-24 grams, were dosed IP at 5 ml/kg with 20 mg/kg UCB-9721 in PBS containing 30% polyethylene glycol 400. Animals were euthanized with CO_2_ at 0.25, 0.5, 1, 2, 4, 6, 8 and 24 hours; two animals per time point. To obtain IV pk data, CBA/J mouse, 19-26 grams, were dosed at 5ml/kg with 1 mg/kg with UCB-9721 in 5:40:55 DMSO:glycerol formal:0.9% saline. Animals were euthanized with CO_2_ at 0.083, 0.25, 0.5, 1, 2, 4, 6, and 8 hours; two animals per time point. Blood samples were collected via cardiac puncture and transferred to Mini Collect heparin tubes (Greiner). Brain and lung samples were collected on dry ice for drug tissue exposure levels. Liver samples were collected in 10% formalin. A satellite group of 6 mice were dosed with 5mg/kg UCB-9721 in the same vehicle. Blood was collected in Mini Collect heparin tubes via cheek bleeds at 0.25, 1, and 4 hours. Terminal bleeds were collected after CO_2_ euthanasia and cardiac puncture at 0.5, 2, and 6 hours. Brain and lung tissue samples were collected for analysis of drug levels. Samples were stored on wet ice (<4 hours) before being centrifuged at 2000*xg* for 10 minutes to isolate plasma. Plasma was collected and stored at -80°C until analysis by LC-MS/MS.

Plasma samples were diluted, if necessary, to bring them into the standard curve range (1–1000 ng/ml). Samples were crashed in 3 volumes of acetonitrile with internal standard (enalapril) at 200ng/ml final concentration. Samples were vortexed and centrifuged at 14000rpm for 7 minutes. 80ul supernatants were transferred into titer plates and sealed for liquid chromatography-tandem mass spectrometry.

A benchtop binary Sciex ExionLC AC Pump equipped with AC autosampler (20uL), ExionLC controller and AC column oven interfaced with a Sciex 4500 mass spectrometer (MS) was used in this study. Mass spectrometry parameters for individual target analyte and internal standards (IS) were optimized by direct infusion. The following MRM transitions were monitored: target analyte UCB-9721 m/z: 388.038>189; Enalapril (IS) 376.0>91.2; Chloridazon (IS) 221.943>104.0.

The target compound UCB-9721 and ISs were separated using a gradient elution profile containing 0.1% formic acid in both water (A) and acetonitrile (B) mobile phase with a Waters XBridge 3.5μm (50 mm length × 2.1 mm ID) C18 analytical column at room temperature. These samples were passed through the analytical column at a constant flow rate of 0.4 ml/min using gradient elution profile that starts 10%B up to 1.2min, 95%B at 1.2 -3.0 min, 10%B at 3.01 min and held the composition for 1.0 min to re-equilibrate the system. MS data acquisition was performed with Analyst software (version 1.7.3) using positive mode for the target compound UCB-9721 and ISs.

For plasma, the standard curve was analyzed using a least-square linear regression with 1/x*x weighting. Reference standards were excluded from use in calculating the standard curve if the back calculated concentration was >20% different. Pharmacokinetic parameters were calculated using noncompartmental analysis in WinNonLin (Certara). IP dose projections were calculated with standard PK equations with the assumption that an IP dose would be similar to a PO dose with 100% bioavailability.

### Solubility and permeability (*in vitro* absorption) testing

Solubility and permeability of UCB-9721 were determined by Eurofins Discovery Services LLC (St. Charles MO).

#### Solubility

A chromatogram of UCB-9721 (200 µM) along with a UV/VIS spectrum with labeled absorbance maxima, was generated. Aqueous solubility (µM) was then determined by comparing the peak area of the principal peak in a calibration standard (200 µM) containing organic solvent (methanol/water, 60/40, v/v) with the peak area of the corresponding peak in a buffer sample. Chromatographic purity (%) was defined as the peak area of the principal peak relative to the total integrated peak area in the HPLC chromatogram of the calibration standard. A chromatogram of the calibration standard of the test compound, along with a UV/VIS spectrum with labeled absorbance maxima, was generated.

#### Parallel Artificial membrane Permeability Assay (PAMPA)

PAMPA assays were carried out at Eurofins Discovery (St. Charles, MO). UCB-9721 and reference compounds (ketoprofen, phenytoin and furosemide) were diluted from 10 mM DMSO stock solution to a single final concentration into donor solution (Prisma HT buffer; PION, Billerica, MA). The donor solution was placed in contact with the acceptor buffer (Acceptor Sink buffer-7.4; PION) at pH 6.5 with the membrane in between. The sandwich was incubated for 4 hrs at ambient temperature. Samples were then analyzed by HPLC-MS/MS using selected reaction monitoring. The apparent permeability coefficient (P_app_) of the test compound and its recovery were calculated as follows:

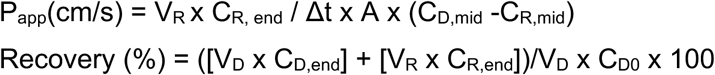

where A = the surface area of the cell monolayer (0.40 cm^2^), C = concentration of the test compound, expressed as peak area, D = donor and R = receiver. 0, mid, and end denote time zero, mid-point, and end of the incubation. Δt is the incubation time and V is the volume of the donor or receiver. The reference compounds yield log Pe (10^-6^ cm/s) values that cover the typical range of the assay from high (phenytoin), mid (ketoprofen) to low (furosemide) permeability.

## Supporting information

This article contains supporting information.

## Acknowledgements

This work was supported by the Drug Discovery Center within the Center of Emerging and Neglected Diseases at the University of California, Berkeley, and by National Institutes of Health (NIH) grants AI139201 (GEW), GM141743 (DMW), S10OD032206 (FS), AI178535 (PS), and AI180804 (SK). AKS was partially supported through NIH T32 grant AI055402 (GEW, PI), and FT was funded by the School of Chemistry at the University of St Andrews. We thank James Spudich and Kainomyx for making these studies possible through a Continuing Research Agreement between Kainomyx and the Ward laboratory at the University of Vermont, established for Dr. Ward’s studies on *Toxoplasma*. We acknowledge that the present work expands upon the studies permitted under that agreement and appreciate their recognition of that expanded scope. We also thank Drs. Godfree Mlambo and Abhai Tripathi, and Chris Kizito, the team of the parasitology and insectary core facilities at the Johns Hopkins Malaria Research Institute for their outstanding work and Bloomberg Philanthropies for their support of these facilities. We also acknowledge the Johns Hopkins School of Medicine Microscopy Facility (MicFac) and the grants supporting the High Content Imager R01GM28007 and R01GM66817.

The authors declare that they have no conflicts of interest with the contents of this article.

