## Supporting information for "Identification of UCB-9721 as a potent inhibitor of MyoA, the essential class XIV myosin motor of apicomplexan parasites"

Table of Contents:

| | |
|---|---|
| Page SI-2 | Figure S1 |
| Page SI-3 | Figure S2 |
| Pages SI-4 to SI-5 | Figure S3 |
| Page SI-6 | Figure S4 |
| Pages SI-7 to SI-8 | Figure S5 |
| Page SI-9 | Table S1 |
| Page SI-10 | Figure S6 |
| Page SI-11 | Supplemental Video legends |
| Pages SI-12 to SI-29 | Supplemental Methods: Compound synthesis |
| Pages SI-30 to SI-38 | Analytical characterization of synthesized compounds: NMR spectra |

**A**

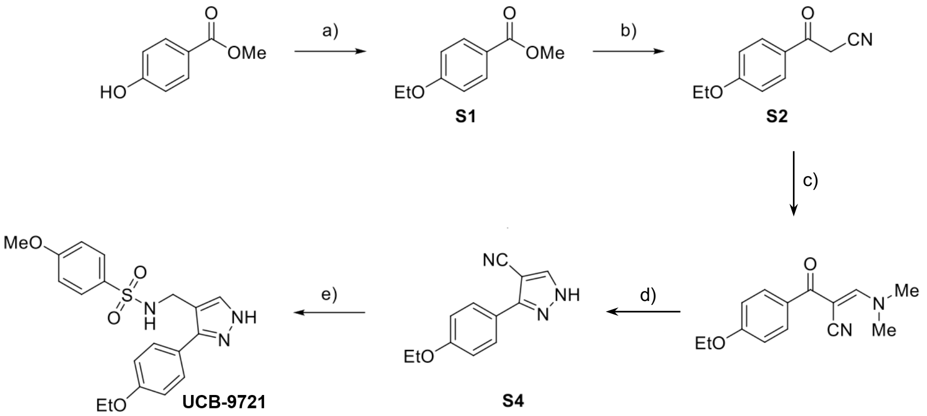


**B**

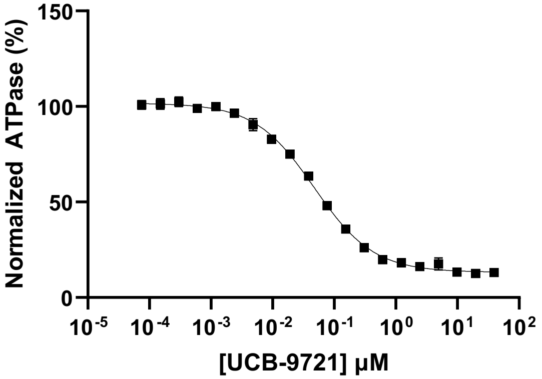


| Test Enzyme | Compound Source | $IC_{50}$ (μM) | 95% C.I. |
|---|---|---|---|
| TgMyoA | Resynthesized | 0.047 | 0.042-0.053 |

**<u>Figure S1</u>. Synthesis of UCB-9721 and inhibitory activity of the synthesized compound. (A)** General synthetic scheme for UCB-9721; see pages SI-12 to SI-38 for details of synthesis and analytical characterization. Reagents and conditions: a) Reagents and conditions a) ethyl bromide (3.00 eq.), $K_2CO_3$ (3.00 eq.), acetone, reflux, 16 hrs, 94%; b) i) $CH_3CN$ (1.10 eq.), n-BuLi (1.15 M in hexane, 1.10 eq.), THF, – 78 °C, 1 hr; ii) **S1**, THF, r.t., 16 hrs, 74%; c) DMF-DMA (1.10 eq.), toluene, r.t., 16 hrs, 84%; d) $NH_2NH_2/H_2O$ (64 wt%, 1.50 eq.), MeOH, r.t., 16 hrs, 89%; e) i) $LiAlH_4$ (2.4 M in THF, 6.00 eq.), THF, 0 – 40 °C, 48 h; ii) amine **S5**, 4-methoxybenzenesulfonyl chloride (1.05 eq.), i-$Pr_2NEt$ (1.05 eq.), $CH_2Cl_2$, – 20 °C – r.t., 16 hrs, 52% over two steps. **(B)** Dose-dependent inhibition of TgMyoA actin-activated ATPase activity by resynthesized UCB-9721, normalized to DMSO (vehicle) control. The table shows the calculated $EC_{50}$, with 95% confidence interval (C.I.).

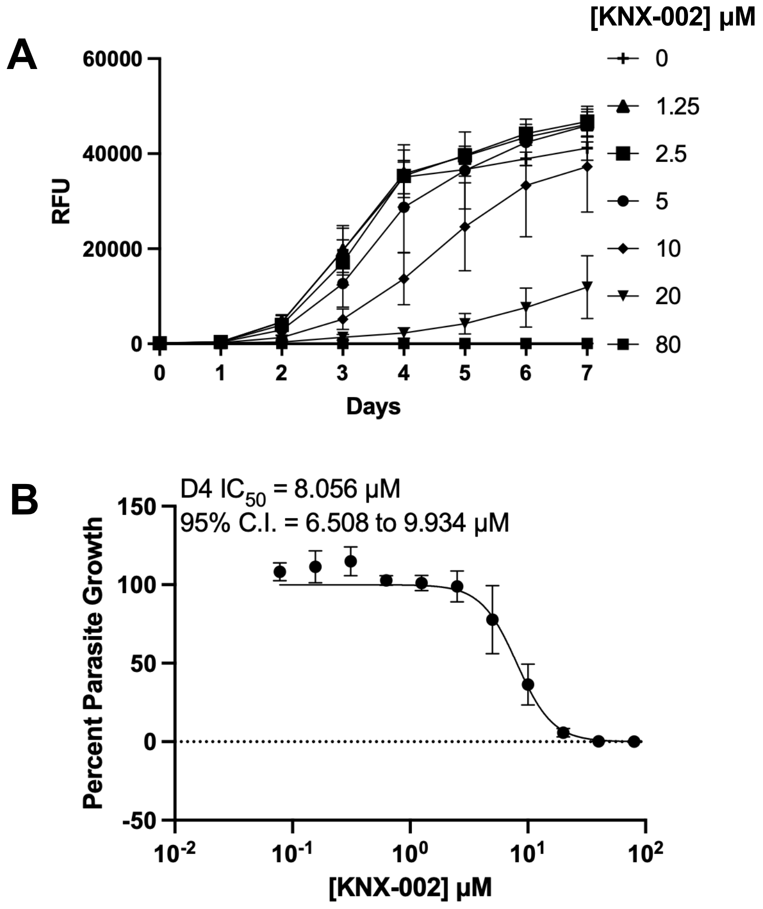


**Figure S2: KNX-002 inhibits parasite growth. (A)** Selected growth curves from a 12-point compound dilution series of tdTomato-expressing parasites in the presence of the indicated concentrations of KNX-002. Fluorescence was measured daily post infection; RFU = relative fluorescence units. Data shown are the mean ± SEM of 4 biological replicates, each consisting of three technical replicates, all done in parallel with the UCB-9721 growth inhibition assays shown in Figure 3C. **(B)** $IC_{50}$ curve and 95% C.I. calculated from growth assay data in **(A)** on day 4 post infection (D4), normalized to the DMSO control (0 μM) and total inhibition (80 μM) with KNX-002; error bars indicate SEM.

**A**

| Matrix | Route | Dose (mg/kg) | Rsq | Half-life (hr) | Tmax (hr) | Cmax (ng/mL) | AUClast (hr·ng/mL) | AUCINF (hr·ng/mL) | AUC % Extrap | Cl (mL/hr/kg) | Vss (mL/kg) | F (%) | Tissue:Plasma Cmax (%) | Tissue:Plasma AUC (%) |
|---|---|---|---|---|---|---|---|---|---|---|---|---|---|---|
| Plasma | IV | 1 | 0.945 | 0.23 | 0.083 | 856.6 | 268.8 | 269.6 | 0.3 | 3709.4 | 881.5 | — | — | — |
| Brain | IV | 1 | — | — | 0.083 | 17.9 | 4.3 | — | — | — | — | — | 2.1 | — |
| Lung | IV | 1 | 0.987 | 0.162 | 0.083 | 305.4 | 79.8 | 81 | 1.5 | — | — | — | 35.7 | 30 |
| Plasma | IP | 20 | 0.827 | 0.35 | 0.25 | 14642 | 5346.7 | 5348.8 | 0.017 | 3740 | N/A | 99.2 | — | — |
| Brain | IP | 20 | — | — | 0.25 | 188.4 | 88.9 | — | — | — | — | — | 1.3 | — |
| Lung | IP | 20 | 1 | 0.194 | 0.25 | 2068.5 | 974.3 | 975.8 | 0.157 | — | — | — | 14.1 | 18.2 |
| Plasma | IP | 5 | 0.95 | 0.22 | 0.25 | 2402 | 922.4 | 924.5 | 0.19 | 5408.1 | N/A | 68.6 | — | — |

**B**

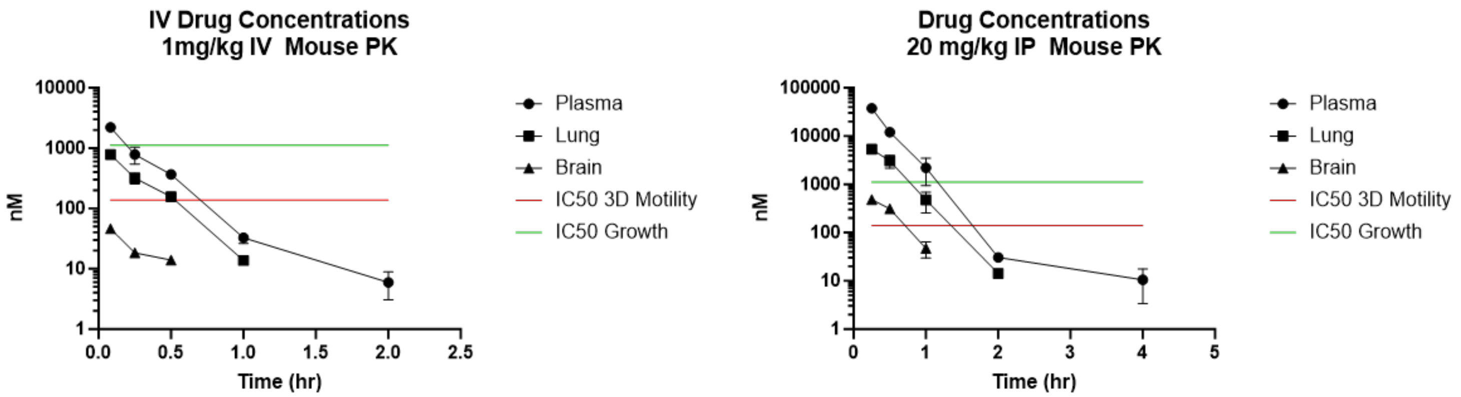


**C**

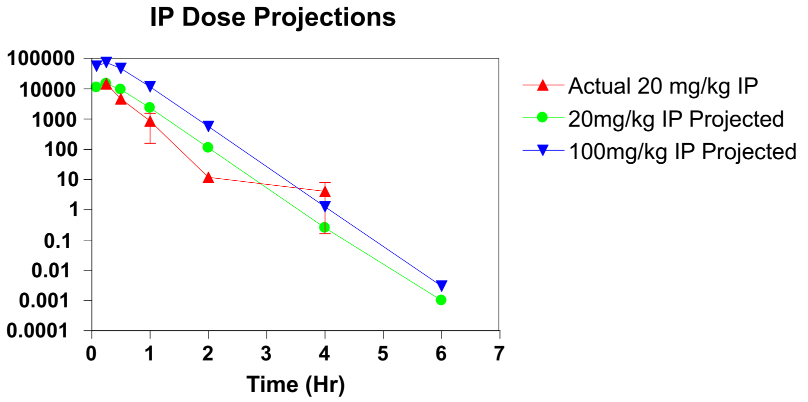


**D**

| *in vitro* PK assays | | |
|---|---|---|
| | Permeability $10^{-6}$ cm/sec (stdev) | Percent Recovery |
| PAMPA pH 6.5 | 5.00 (0.16) | 92% |

| Aqueous Solubility | Solubitlity uM (stdev) |
|---|---|
| PBS pH 7.4 | 26.7 (2.95) |
| PBS pH 6.5 | 19.0 (1.48) |

**Figure S3 (previous page): Pharmacokinetic (PK) and *in vitro* ADME properties of UCB-9721.** (A) Summary table of *in vivo* PK parameters from CBA/J mice calculated in WinNonLin using noncompartmental analysis: Rsq = $R^2$. (B) Graphs from and IV 1mg/kg and IP 20mg/kg pharmacokinetic studies in mice. For comparison to $IC_{50}$s from *T. gondii* growth and motility assays, drug levels (ng/ml) were converted to nM. For tissues, 1 gram of tissue was treated as 1 gram of water. (C) IP dose projections (20mg/kg [green] and 100mg/kg [blue]) were made using standard PK equations for predicting plasma exposures from an oral dose assuming 100% bioavailability and a similar $C_{max}$ observed in the 20mg/kg IP PK study (red). (D) Results of *in vitro* ADME assays. Top panel is Parallel Artificial Membrane Permeability Assay (PAMPA) assay. The bottom panel shows aqueous solubility at pH 6.5 or pH 7.4, each measured in duplicate; means and standard deviations are shown.

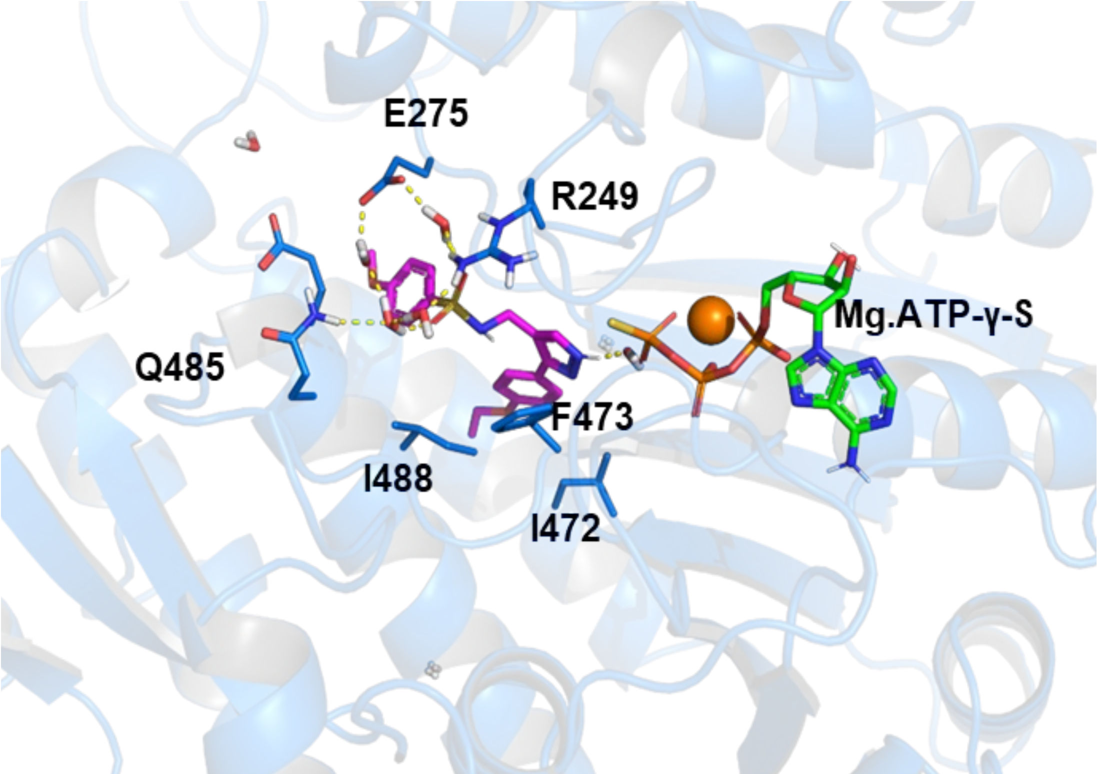


**<u>Figure S4</u>: *in silico* docking predicts UCB-9721 binding to TgMyoA at a site adjacent to but distinct from the nucleotide binding site.** The binding mode of UCB-9721 to TgMyoA predicted by *in silico* docking and equilibrated with 20 ns molecular dynamics (MD) simulation is shown. The highlighted structures are UCB-9721 (magenta/purple), TgMyoA (blue ribbons), bound nucleotide (colored by atom) and water added to the simulation (red and white). Important TgMyoA side chains, ligands and water molecules are depicted as sticks and labeled.

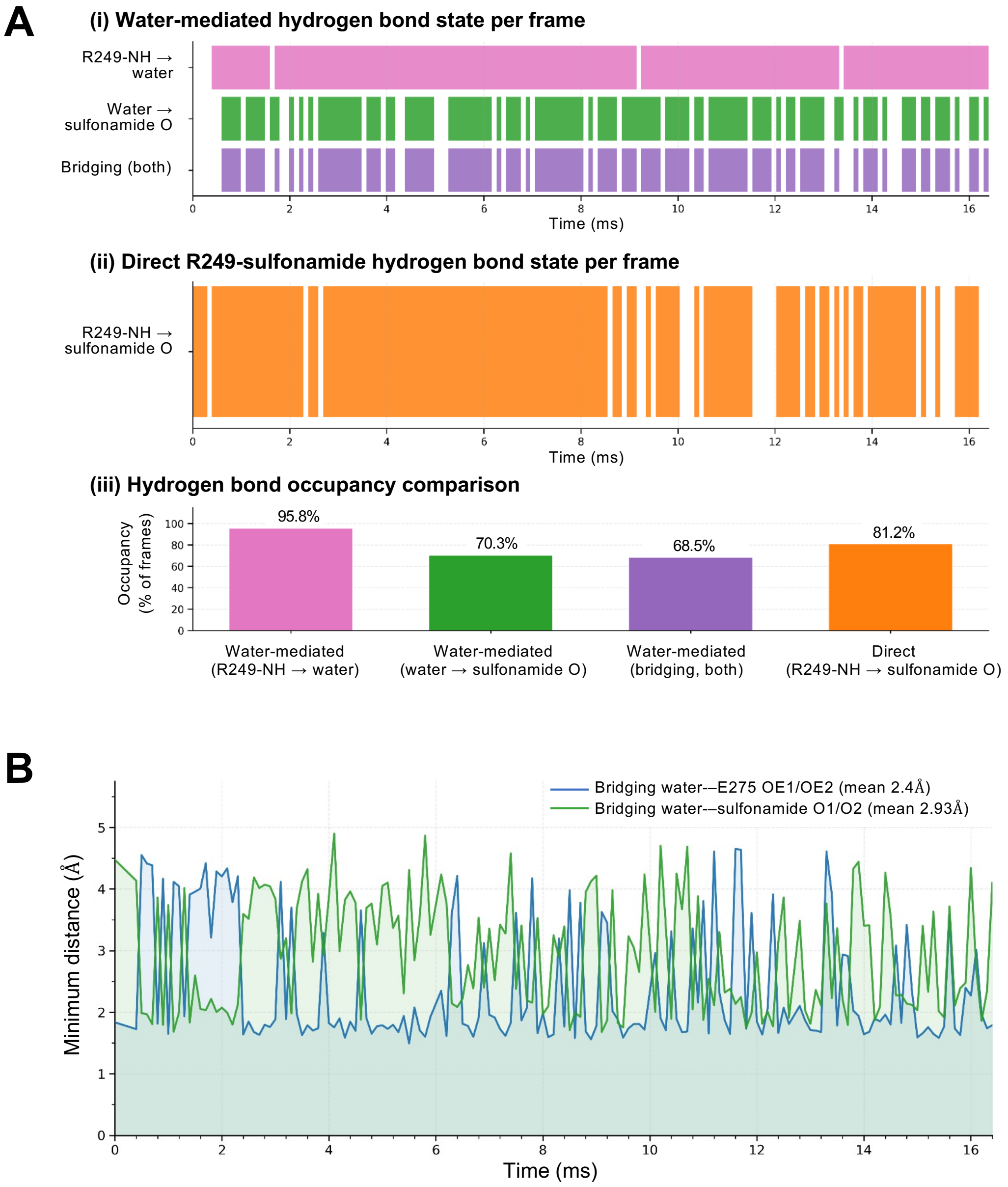
A
(i) Water-mediated hydrogen bond state per frame
R249-NH → water
Water → sulfonamide O
Bridging (both)
Time (ms)
(ii) Direct R249-sulfonamide hydrogen bond state per frame
R249-NH → sulfonamide O
Time (ms)
(iii) Hydrogen bond occupancy comparison
Occupancy (% of frames)
95.8%
70.3%
68.5%
81.2%
Water-mediated (R249-NH → water)
Water-mediated (water → sulfonamide O)
Water-mediated (bridging, both)
Direct (R249-NH → sulfonamide O)
B
Bridging water-–E275 OE1/OE2 (mean 2.4Å)
Bridging water-–sulfonamide O1/O2 (mean 2.93Å)
Minimum distance (Å)
Time (ms)

**<u>Figure S5 (previous page)</u>: (A)** Analysis of the R249A-sulfonamide interaction**.** Direct and water-mediated hydrogen bonds between R249 and the sulfonamide oxygens along the 16.4 ns MD trajectory. (i) Per-frame state of the three water-mediated modes: R249-N–H → water (pink, 95.8%), water → sulfonamide O (green, 70.3%), and both bonds simultaneously present, defining a fully bridged R249···$H_2O$···sulfonamide configuration (purple, 68.5%). (ii) Per-frame state of the direct R249-N–H···O–S hydrogen bond (orange, 81.2%). (iii) Occupancies of the four modes (% of all frames). Hydrogen bonds were assigned using D···A ≤ 3.5 Å and D–H···A ≥ 150° (distance-only for the direct bond) **(B)** Fluctuation of the distances between the bridging water molecule and the side-chain oxygen atoms of E275, as well as the sulfonamide oxygen atoms, during a 16 ns MD simulation. The short mean distances indicate that the bridging water molecule interacts strongly and simultaneously with both E275 and the sulfonamide.

**Table S1: Effects of the UCB-9721 analogs on TgMyoA actin-activated ATPase activity and parasite growth.**

| Structure | Compound | ATPase $IC_{50}$ (µM) (95% C.I.) | Structure | Compound | ATPase $IC_{50}$ (µM) (95% C.I.) | Parasite Growth $IC_{50}$ µM (95% C.I.) |
|---|---|---|---|---|---|---|
| 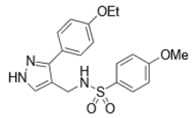 | **UCB-9721** **from screen** | 0.031 (0.029 - 0.032) | 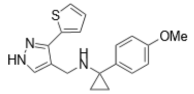 | **KNX-002** | 1.839 (1.700 - 1.997) | 8.056 (6.508-9.934) |
| **Compound Screen** | | | **Synthesized Compounds** | | | |
| 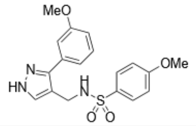 | **UCB-82-813** | <0.076 | 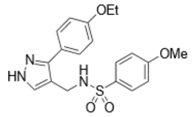 | **UCB-9721** **resynthesized** | 0.047 (0.042 - 0.053) | 2.488 (1.360-2.939) |
| 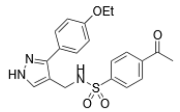 | **UCB-82-823** | 0.258 (0.168 - 0.335) | 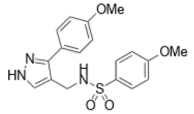 | **VEST16** | 0.382 (0.359 - 0.407) | 9.212 (7.035-11.928) |
| 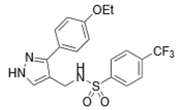 | **UCB-82-817** | 0.414 (0.277 - 0.544) | 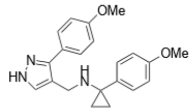 | **VEST17** | >39 | 29.993 (16.739-69.701) |
| 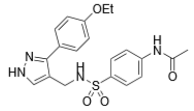 | **UCB-82-815** | 0.751 (0.601-0.899) | 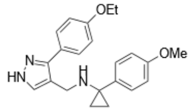 | **VEST18** | 12.105 (10.722 - 14.121) | 5.897 (4.766-7.307) |
| 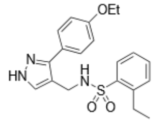 | **UCB-82-825** | 2.196 (1.852 - 2.644) | 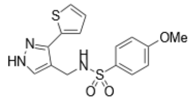 | **VEST19** | 0.070 (0.068 - 0.073) | 0.366 (0.291-0.469) |
| 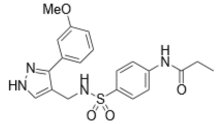 | **UCB-82-796** | 3.089 (2.558 - 3.814) | | | | |
| 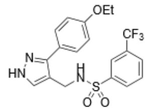 | **UCB-82-821** | 5.718 (4.299 - 9.812) | | | | |

#### Supplemental video legends

**<u>Videos S1A, S1B</u>: Effect of UCB-9721 on TgMyoA *in vitro* motility.** (Video S1A): *in vitro* motility assay done in the presence of DMSO (vehicle only); average filament speed 4.5 μm/sec. (Video S1B): *in vitro* motility assay done in the presence of 5μM UCB-9721; average filament speed 1.1 μm/sec.

**<u>Videos S2A-C</u>: *in vitro* motility assays with wild-type and mutant myosins.** (Video S2A): $TgMyoA^{WT}$, average filament speed 4.6 μm/sec. (Video S2B): $TgMyoA^{R249A}$, average filament speed 0.0 μm/sec. (Video S2C): $TgMyoA^{E275A}$, average filament speed 0.6 μm/sec)

### Supporting Information – Compound synthesis

#### General Considerations

All reagents were purchased from Aldrich (UK), Alfa Aesar, Apollo Scientific or Fluorochem, and used without further purification. All reactions requiring dry conditions were conducted in flame-dried glassware under an inert atmosphere ($N_2$), using standard vacuum line techniques in oven-dried glassware. Dry solvents ($CH_2Cl_2$, DMF, THF) were obtained after passing through an alumina column (Mbraun SPS-800). Standard syringe techniques were used for dry addition of liquids.

Room temperature (r.t.) refers to 20 – 25 °C. Temperatures of 0 °C, –20 °C, and –78 °C were obtained using ice/water, methanol/water/liquid $N_2$, and ethyl acetate/liquid $N_2$ baths, respectively. Reaction involving heating were performed using DrySyn blocks and a contact thermocouple.

Nuclear magnetic resonance (NMR) spectra were recorded on a Bruker Advance II 400 ($^1$H 400; $^{13}$C 101 MHz) or Bruker Ascend 500 ($^1$H 500; $^{13}$C 126 MHz). $^{13}$C NMR spectra were taken with a DEPTQ pulse sequence. Chemical shifts are expressed as δ in units of ppm. Multiplicities are described using the following abbreviations: bs=broad signal, s=singlet, d=doublet, dd=doublet of doublets, t=triplet, q=quartet, and m=multiplet and the J couplings are reported in Hz. NMR spectra were processed using MestReNova. Peaks were assigned with the aid of the two-dimensional NMR spectroscopic techniques COSY (2D $^1$H-$^1$H correlated spectroscopy), HMBC (2D $^1$H-$^{13}$C heteronuclear multiple-bond correlation spectroscopy), and HSQC (2D $^1$H-$^{13}$C heteronuclear single quantum coherence) if necessary. Column chromatography was performed using Davisil® silica (40–63 µm, 230–400 mesh). Thin layer chromatography was performed on pre-coated glass plates (Silica Gel 60A, Fluorochem) and visualised under UV light (254 nm) or by staining with $KMnO_4$ and vanillin. Infrared spectra were recorded on a Shimadzu IR Affinity$^{-1}$ Fourier transform IR spectrophotometer as thin films. Absorption maxima are reported in wavenumbers (cm$^{-1}$) IR. Melting points were recorded on an Electrothermal 9100 melting point apparatus, (*dec*) refers to decomposition. Mass spectrometry data were acquired by Mrs Caroline Horsburgh in the University of St Andrews School of Chemistry Mass Spectrometry Service.

#### General Scheme for the Synthesis of UCB-9721

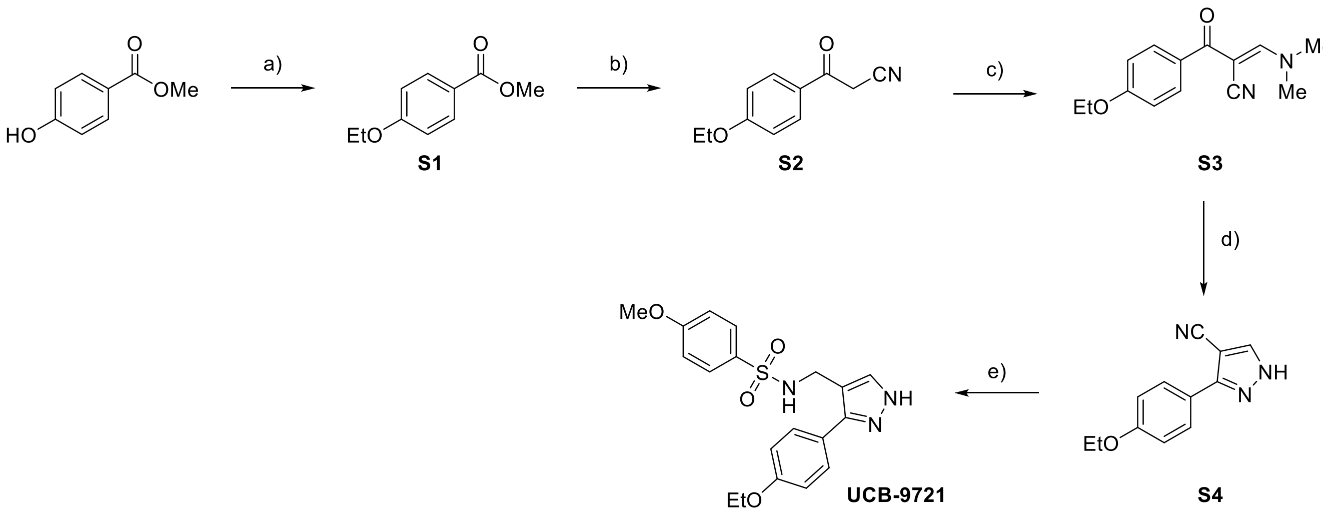


**Scheme 1.1** *Reagents and conditions*: a) ethyl bromide (3.00 eq.), $K_2CO_3$ (3.00 eq.), acetone, reflux, 16 h, 94%; b) i) $CH_3CN$ (1.10 eq.), *n*-BuLi (1.15 M in hexane, 1.10 eq.), THF, – 78 °C, 1 h; ii) **1**, THF, r.t., 16 h, 74%; c) DMF-DMA (1.10 eq.), toluene, r.t., 16 h, 84%; d) $NH_2NH_2 \cdot H_2O$ (64 wt%, 1.50 eq.), MeOH, r.t., 16 h, 89%; e) $LiAlH_4$ (2.4 M in THF, 6.00 eq.), THF, 0 – 40 °C, 48 h; ii) 4-methoxybenzenesulfonyl chloride (1.05 eq.), *i*-$Pr_2$Net (1.05 eq.), $CH_2Cl_2$, – 20 °C – r.t., 16 hrs, 52% over two steps.

#### Methyl 4-ethoxybenzoate (S1)[1]

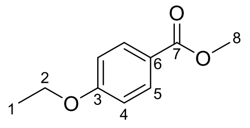


Commercially available methyl 4-hydroxybenzoate (20.0 g, 131 mmol, 1.00 eq.) was dissolved in acetone (250 mL). $K_2CO_3$ (54.4 g, 394 mmol, 3.00 eq.) and ethyl bromide (29.2 mL, 394 mmol, 3.00 eq.) were added. The reaction mixture was stirred at reflux for 16 hours. After cooling to room temperature, the reaction mixture was filtered, rinsing with acetone (20 mL) to remove the excess of $K_2CO_3$. The filtrate was concentrated *in vacuo*, and the crude mixture was dissolved in EtOAc (300 mL). The organic phase was washed with 1 M NaOH (3 × 100 mL), dried ($MgSO_4$), filtered, and concentrated *in vacuo* to give **S1** (22.3 g, 124 mmol, 94%) as a pale-yellow solid.

**M.P.** 35-36 °C (*lit.* 35-36 °C).[1]

**$\delta_H$** (500 MHz, $CDCl_3$) 7.97 (2 H, d, *J* 9.1, 2 × H-5), 6.89 (2 H, d, *J* 9.1, 2 × H-4), 4.08 (2 H, q, *J* 7.0, 2 × H-2), 3.88 (3 H, s, 3 × H-8), 1.43 (3 H, t, *J* 7.0, 3 × H-1).

Analytical data were in agreement with the literature.[1]

#### 3-(4-Ethoxyphenyl)-3-oxopropanenitrile (S2)[2]

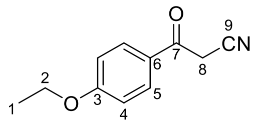


*n*-BuLi solution (1.15 M in hexane, 2.65 mL, 3.05 mmol, 1.10 eq.) was added dropwise at – 78 °C to a solution of freshly distilled acetonitrile[3] (0.16 mL, 3.05 mmol, 1.10 eq.) and dry THF (3.05 mL), and the reaction mixture was stirred at – 78 °C for 1 hour. Then, a solution of methyl 4-ethoxybenzoate (**S1**) (0.500 g, 2.77 mmol, 1.00 eq.) in dry THF (3.70 mL) was added dropwise at – 78 °C. The resulting mixture was stirred for 16 hours at room temperature. After cooling to 0 °C, the reaction mixture was quenched with aqueous 2 M HCl (7 mL) and the aqueous layer was extracted with $Et_2O$ (3 × 30 mL). The combined organic extracts were washed with brine (1 × 15 mL), dried ($MgSO_4$), filtered, and concentrated *in vacuo*. The crude product was purified by silica gel column chromatography (0 – 35% EtOAc in hexane) to give **S2** (0.390 g, 2.06 mmol, 74%) as a pale-yellow powder.

**M.P.** 124-125 °C (*lit*. 119.5-121.4 °C).[2]

**$\delta_H$** (500 MHz, $CDCl_3$) 7.88 (2 H, d, *J* 8.9, 2 × H-5), 6.95 (2 H, d, *J* 8.8, 2 × H-4), 4.12 (2 H, q, *J* 7.0, H-2), 4.01 (2 H, s, 2 × H-8), 1.45 (1 H, t, *J* 7.0, 3 × H-1).

**$\delta_C$** (126 MHz, $CDCl_3$) 185.5 (C-7), 164.3 (C-3), 131.1 (C-5), 127.2 (C-9), 114.9 (C-4), 114.3 (C-6), 64.2 (C-2), 29.1 (C-8), 14.7 (C-1).

Analytical data were in agreement with the literature.[2]

#### (*E*)-3-(dimethylamino)-2-(4-ethoxybenzoyl)acrylonitrile (S3)[4]

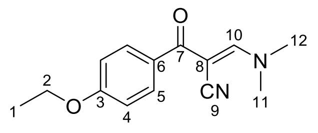


3-(4-Ethoxyphenyl)-3-oxopropanenitrile (**S2**) (0.927 g, 4.90 mmol, 1.00 eq.) was dissolved in dry toluene (15 mL), and *N,N*-Dimethylformamide dimethyl acetal (DMF-DMA) (0.716 mL, 5.39 mmol, 1.10 eq.) was added dropwise. The reaction mixture was stirred for 16 hours at room temperature, and then concentrated *in vacuo*. The crude product was purified by silica gel column chromatography (0 – 4% MeOH in $CH_2Cl_2$) to give **S3** (1.01 g, 4.13 mmol, 84%) as a pale-yellow solid.

**M.P.** 108-109 °C (*lit*. 110-115 °C).[4]

**IR $\nu_{max}$** (film) $cm^{-1}$ 2980, 2359, 2341, 2193, 1647, 1601, 1290, 1254, 1175.

**$\delta_H$** (500 MHz, $CDCl_3$) 7.86 (1 H, s, H-10), 7.78 (2 H, d, *J* 8.8, 2 × H-5), 6.86 (2 H, d, *J* 8.7, 2 × H-4), 4.03 (2 H, q, *J* 6.9, 2 × H-2), 3.40 (3 H, s, 3 × H-11 or H-12), 3.19 (3 H, s, 3 × H-11 or H-12), 1.38 (3 H, t, *J* 7.0, H-1).

**$\delta_C$** (126 MHz, $CDCl_3$) 188.8 (C-7), 161.8 (C-3), 159.3 (C-10), 130.7 (C-6), 130.6 (C-5), 120.7 (C-9), 113.7 (C-4), 79.2 (C-8), 63.6 (C-2), 48.1 (C-11 or C-12), 38.9 (C-11 or C-12), 14.7 (C-1).

**HRMS (ESI$^+$)** m/z calculated for $C_{14}H_{16}N_2O_2Na$ $[M+Na]^+$ 267.1104, found 267.1107.

#### 3-(4-Ethoxyphenyl)-1H-pyrazole-4-carbonitrile (S4)

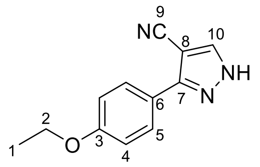


(*E*)-3-(Dimethylamino)-2-(4-ethoxybenzoyl)acrylonitrile (**S3**) (2.26 g, 9.25 mmol, 1.00 eq.) was dissolved in methanol (90 mL), and hydrazine monohydrate (64 wt% $NH_2NH_2$ in $H_2O$, 1.05 mL, 13.9 mmol, 1.50 eq.) was added. The reaction mixture was stirred for 16 hours at room temperature, and then concentrated *in vacuo*. The resulting crude reaction mixture was purified by silica gel column chromatography (0 – 2% MeOH in $CH_2Cl_2$) to give **S4** (1.75 g, 8.23 mmol, 89%) as a pale-yellow solid.

**M.P.** 118-119 °C.

**IR $\nu_{max}$** (film) $cm^{-1}$ 2978, 2230, 1612, 1250, 1182, 732.

**$\delta_H$** (500 MHz, $CDCl_3$) 7.89 (1 H, s, H-10), 7.72 (2 H, d, *J* 8.8, 2 × H-5), 6.94 (2 H, d, *J* 8.8, 2 × H-4), 4.06 (2 H, q, *J* 7.0, 2 × H-2), 1.44 (3 H, t, *J* 7.0, 3 × H-1).

**$\delta_C$** (126 MHz, $CDCl_3$) 160.7 (C-3), 152.0 (C-7), 140.9 (C-10), 128.3 (C-5), 119.9 (C-6, determined through HMBC), 115.2 (C-4), 114.7 (C-9), 89.0 (C-8), 63.8 (C-2), 14.8 (C-1).

#### HRMS (ESI$^+$) m/z calculated for $C_{12}H_{11}N_3ONa$ $[M+Na]^+$ 236.0800, found 236.0790.*N*-((5-(4-Ethoxyphenyl)-1H-pyrazol-4-yl)methyl)-4-methoxybenzenesulfonamide (UCB-9721)

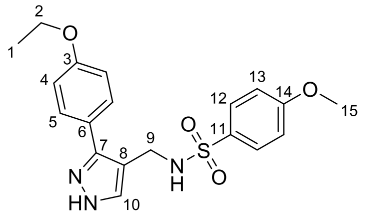


**UCB-9721**

Synthesized in two steps *via* (3-(4-Ethoxyphenyl)-1H-pyrazol-4-yl)methanamine

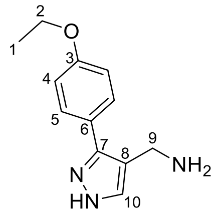


3-(4-Ethoxyphenyl)-1H-pyrazole-4-carbonitrile (**S4**) (0.614 g, 2.88 mmol, 1.00 eq.) was dissolved in dry THF (40 mL), and $LiAlH_4$ solution (2.0 M in THF, 8.64 mL, 17.3 mmol, 6.00 eq.) was added dropwise at 0 °C. The reaction mixture was stirred at 40 °C for 48 hours. The reaction was cooled to 0 °C, quenched by slow addition of cold EtOH (20 mL) followed by saturated aqueous $NH_4Cl$ solution (15 mL). The suspension was stirred for 2 hours at 0 °C, and then concentrated *in vacuo*. The resulting mixture was diluted with aqueous 1 M NaOH (15 mL), and the aqueous layer was extracted with $CH_2Cl_2$ (3 × 30 mL). The combined organic extracts were washed with brine (2 × 15 mL), dried ($MgSO_4$), filtered, and concentrated *in vacuo* to give the corresponding amine, which was used in the next step without any further purification.

**$^1$H NMR analysis of amine**

**$\delta_H$** (400 MHz, $CDCl_3$) 7.57 (1 H, s, H-10), 7.49 (2 H, d, *J* 8.7, 2 × H-5), 6.94 (2 H, d, *J* 8.7, 2 × H-4), 4.06 (2 H, q, *J* 7.0, 2 × H-2), 3.89 (2 H, s, 2 × H-9), 1.43 (2 H, t, *J* 7.0, 3 × H-1).

The resulting amine (2.88 mmol, 1.00 eq.) was dissolved in dry $CH_2Cl_2$ (35 mL), and DIPEA (0.53 mL, 3.02 mmol, 1.05 eq.) was added dropwise at – 20 °C. The resulting mixture was stirred for 15 minutes, and 4-methoxybenzenesulfonyl chloride (0.625 g, 3.02 mmol, 1.05 eq.) was added. The reaction mixture was stirred at room temperature for 16 hours. The resulting mixture was quenched with saturated aqueous $NaHCO_3$ solution (15 mL), and diluted with $CH_2Cl_2$ (30 mL). The organic phase was washed with saturated aqueous $NaHCO_3$ solution (3 × 15), dried ($MgSO_4$), filtered, and concentrated *in vacuo*. The resulting crude was purified by silica gel column chromatography (0 – 10% MeOH in $CH_2Cl_2$) to give **UCB-9721** (0.581 g, 1.50 mmol, 52% over two steps) as a white powder.

**Analytical characterization for UCB-9721**

**M.P.** 128-129 °C.

**IR $\nu_{max}$** (film) $cm^{-1}$ 3279, 2927, 1597, 1496, 1100, 832.

**$\delta_H$** (500 MHz, $CDCl_3$) 7.75 (2 H, d, *J* 8.9, 2 × H-12), 7.33 (1 H, s, H-10), 7.27 (2 H, d, *J* 8.8, 2 × H-5), 6.93 (2 H, d, *J* 8.9, 2 × H-13), 6.81 (2 H, d, *J* 8.8, 2 × H-4), 5.21 (1 H, brs, NH), 4.09 (2 H, d, *J* 4.4, 2 × H-9), 4.05 (2 H, q, *J* 7.0, 2 × H-2), 3.88 (3 H, s, 3 × H-15), 1.45 (3 H, t, *J* 7.0, 3 × H-1).

**δ$_C$** (126 MHz, $CDCl_3$) 165.1 (C-14), 159.3 (C-3), 145.5 (C-7), 136.1 (C-10), 131.3 (C-11), 129.5 (C-12), 128.8 (C-5), 115.0 (C-4), 114.4 (C-13), 112.7 (C-8), 122.7 (C-6), 63.6 (C-2), 57.1 (C-15), 38.6 (C-9), 15.5 (C-1).

**HRMS (ESI$^+$)** m/z calculated for $C_{19}H_{21}N_3O_4SNa$ $[M+Na]^+$ 410.1151, found 410.1149.

#### General Scheme for the Synthesis of VEST16

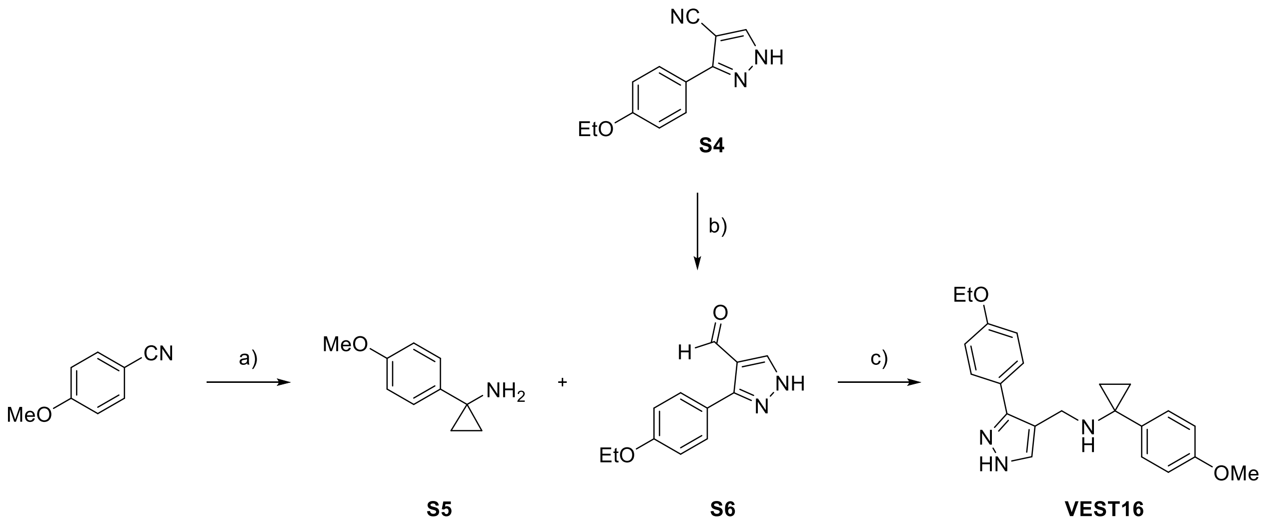


**Scheme S1** *Reagents and conditions*: a) i) Ti(IV) isopropoxide (1.10 eq.), EtMgBr (3.0 M in $Et_2O$, 2.20 eq.), $Et_2O$, – 78 °C – r.t., 2 hrs; ii) $BF_3{\cdot}Et_2O$ (2.00 eq.), 0 °C – r.t., 2 hrs, 35%; b) DIBAL-H (1.2 M in toluene), $CH_2Cl_2$, – 20 °C – r.t., 3 hrs, 59%; c) i) **S5** (1.00 eq.), **S6** (1.15 eq.), THF, r.t., 16 hrs; ii) $NaBH_4$ (1.30 eq.), MeOH, 0 °C – r.t, 3 hrs, 48% over two steps.

##### 1-(4-Methoxyphenyl)cyclopropan-1-amine (S5)[5]

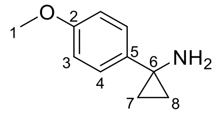


Commercially available 4-methoxybenzonitrile (0.250 g, 1.88 mmol, 1.00 eq.) and titanium(IV) isopropoxide (0.618 mL, 2.07 mmol, 1.10 eq.) were dissolved in $Et_2O$ (10 mL), and ethyl magnesium bromide solution (3.0 M in $Et_2O$, 1.38 mL, 4.14 mmol, 2.20 eq.) was added at – 78 °C. The resulting mixture was stirred at room temperature for 2 hours. Then, boron trifluoride diethyl etherate (0.464 mL, 3.76 mmol, 2.00 eq.) was added at 0 °C, and the reaction was stirred for 2 hours at room temperature. The resulting mixture was quenched with 1 M $HCl_{(aq)}$ and 1 M $NaOH_{(aq)}$. The aqueous layer was extracted with $Et_2O$ (3 × 30 mL). The combined organic extracts were dried ($MgSO_4$), filtered, and concentrated *in vacuo*. The crude reaction mixture was purified by silica gel column chromatography (0 – 65% EtOAc in hexane) to give **S5** (0.0730 g, 0.448 mmol, 35%) as a colorless oil.

**$\delta_H$** (400 MHz, $CDCl_3$) 7.25 (2 H, d, *J* 9.0, 2 × H-4), 6.90 – 6.81 (2 H, d, *J* 9.0, 2 × H-3), 3.80 (3 H, s, 3 × H-1), 1.02 – 0.98 (2 H, m, 1 × H-7, 1 × H-8), 0.93 – 0.89 (2 H, m, 1 × H-7, 1 × H-8).

Analytical data are in agreement with the literature.[5]

#### 3-(4-Ethoxyphenyl)-1H-pyrazole-4-carbaldehyde (S6)[6]

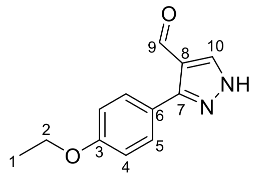

3-(4-Ethoxyphenyl)-1H-pyrazole-4-carbonitrile (**S4**) (0.100, 0.469 mmol, 1.00 eq.) was dissolved in dry $CH_2Cl_2$, and a DIBAL-H solution (1.2 M in toluene, 1.17 mL, 1.41 mmol, 3.00 eq.) was added dropwise at – 20 °C. The reaction mixture was stirred at room temperature for 3 hours. After cooling to 0 °C, the reaction mixture was quenched by slow addition of cold MeOH (3 mL) followed by a saturated aqueous solution of Rochelle salt (5 mL). The resulting mixture was stirred for 30 minutes at – 20 °C, and then concentrated *in vacuo*. The resulting crude reaction mixture was diluted with water (10 mL), and the aqueous layer was extracted with EtOAc (3 × 20 mL). The combined organic extracts were dried ($MgSO_4$), filtered, and concentrated *in vacuo*. The resulting solid was purified by silica gel column chromatography (0 – 2% MeOH in $CH_2Cl_2$) to give **S6** (0.0600 g, 0.277 mmol, 59%) as a white solid.

**M.P.** 162-163 °C (*lit.* 163-164 °C).[6]

**$\delta_H$** (500 MHz, $CDCl_3$) 9.98 (1 H, s, H-9), 8.15 (1 H, s, H-10), 7.59 (2 H, d, *J* 8.8, 2 × H-5), 7.02 (2 H, d, *J* 8.8, 2 × H-4), 4.13 (2 H, q, *J* 7.0, 2 × H-2), 1.49 (3 H, t, *J* 7.0, 3 × H-1).

Analytical data were in agreement with the literature.[6]

#### *N*-((3-(4-Ethoxyphenyl)-1H-pyrazol-4-yl)methyl)-1-(4-methoxyphenyl)cyclopropan-1-amine (VEST16)

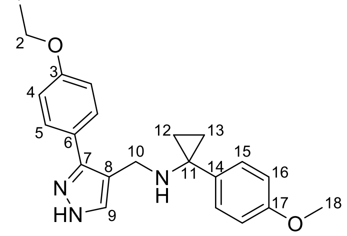

Synthesized in two steps *via* (*E*)-1-(3-(4-Ethoxyphenyl)-1H-pyrazol-4-yl)-N-(1-(4-methoxyphenyl)-cyclopropyl)methanimine

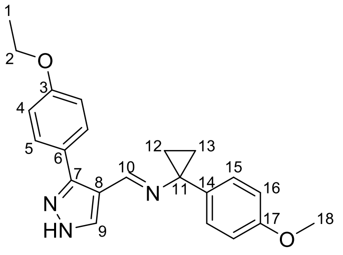


3-(4-ethoxyphenyl)-1H-pyrazole-4-carbaldehyde (**S6**) (0.0500 g, 0.231 mmol, 1.15 eq.) and 1-(4-methoxyphenyl)cyclopropan-1-amine (**S5**) (0.0330 g, 0.198 mmol, 1.00 eq.) were dissolved in dry THF (3 mL). The resulting mixture was stirred for 16 hours at room temperature, and then concentrated *in vacuo* to obtain imine, which was used immediately into without any further purification.

**$^1$H NMR analysis of imine**

**$\delta_H$** (500 MHz, $CDCl_3$) 8.00 (1 H, s, H-9), 7.82 (1 H, s, H-10) 7.28 (2 H, d, *J* 8.8, 2 × H-5), 7.24 (2 H, d, *J* 8.8, 2 × H-15), 6.90 – 6.82 (4 H, m, 2 × H-4, 2 × H-16), 4.05 (2 H, q, *J* 7.0, 2 × H-2), 3.80 (3 H, s, 3 × H-18), 1.41 – 1.36 (2 H, m, 1 × H-12, 1 × H-13), 1.25 – 1.21 (2 H, m, 1 × H-12, 1 × H-13).

The resulting imine was dissolved in dry MeOH (3 mL), and $NaBH_4$ (10 mg, 0.257 mmol, 1.30 eq.) was added at 0 °C. The reaction mixture was stirred at room temperature for 3 hours. The reaction mixture was cooled to 0 °C and quenched with saturated aqueous $NH_4Cl$ solution (5 mL), and water (5 mL). The aqueous layer was extracted with EtOAc (3 × 15 mL). The combined organic extracts were dried ($MgSO_4$), filtered, and concentrated *in vacuo*. The crude reaction mixture was purified by silica gel column chromatography (0 – 10% MeOH in $CH_2Cl_2$) to give **VEST16** (0.0350 g, 0.0960 mmol, 48% over two steps) as a white powder.

**Analytical characterization for VEST16**

**M.P.** 130-131 °C.

**IR $\nu_{max}$** (film) $cm^{-1}$ 2924, 2802, 2357, 1612, 1514, 1263, 1246, 779.

**$\delta_H$** (500 MHz, $CDCl_3$) 7.49 (1 H, s, H-9), 7.43 (2 H, d, *J* 8.7, 2 × H-5), 7.29 (2 H, d, *J* 7.9, 2 × H-15), 6.92 – 6.84 (4 H, m, 2 × H-4, 2 × H-16), 4.08 (2 H, q, *J* 7.0, 2 × H-2), 3.84 (3 H, s, 3 × H-18), 3.67 (2 H, s, 2 × H-10), 1.47 (3 H, t, *J* 7.0, 3 × H-1), 1.04 – 0.99 (2 H, m, 1 × H-12, 1 × H-13), 0.91 – 0.86 (2 H, m, 1 × H-12, 1 × H-13).

**δC** (126 MHz, $CDCl_3$) 158.8 (C-3), 158.3 (C-17), 144.7 (C-7, determined through HMBC), 135.8 (C-9, determined through HSQC), 135.0 (C-14), 129.4 (C-15), 128.7 (C-5), 123.7 (C-6), 116.3 (C-8), 114.6 (C-4), 113.6 (C-16), 63.5 (C-2), 55.3 (C-18), 42.2 (C-11), 40.1 (C-10), 14.9 (C-12, C-13), 14.8 (C-1).

**HRMS (ESI⁺)** m/z calculated for $C_{22}H_{26}O_2N_3$ $[M+H]^+$ 364.2025, found 364.2019.

#### General Scheme for the Synthesis of VEST17

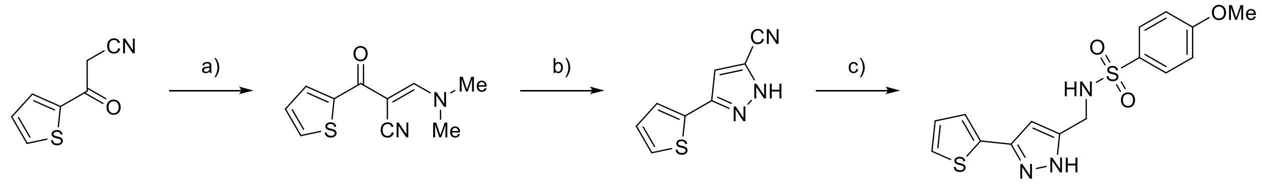


**S7** **S8** **VEST17**

**Scheme 1.3** a) DMF-DMA (1.10 eq.), r.t., 16 hrs, 73%; b) $NH_2NH_2 \cdot H_2O$ (64 wt%, 1.50 eq.), MeOH, r.t., 16 hrs, 83%; c) i) $LiAlH_4$ (2.4 M in THF, 6.00 eq.), THF, 0 – 40 °C, 48 hrs; ii) 4-methoxybenzenesulfonyl chloride (1.05 eq.), *i*-$Pr_2NEt$ (1.05 eq.), $CH_2Cl_2$, – 20 °C – r.t., 16 hrs, 24% over two steps.

#### (*E*)-3-(Dimethylamino)-2-(thiophene-2-carbonyl)acrylonitrile (S7)[7]

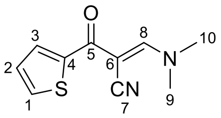


Commercially available 3-oxo-3-(thiophen-2-yl)propanenitrile (1.00 g, 6.61 mmol, 1.00 eq.) was dissolved in dry toluene (22 mL), and *N,N*-Dimethylformamide dimethyl acetal (DMF-DMA) (0.970 mL, 7.28 mmol, 1.10 eq.) was added dropwise. The reaction mixture was stirred for 16 hours at room temperature, and then concentrated *in vacuo*. The crude product was purified by silica gel column chromatography (0 – 10% MeOH in $CH_2Cl_2$) to give **S7** (1.00 g, 4.85 mmol, 73%) as a pale-yellow solid.

**M.P.** 153-154 °C (*lit.* 153-154 °C).[8]

**IR** $\nu_{max}$ (film) $cm^{-1}$ 3099, 2193, 1631, 1556, 1514, 1401, 1300, 727

**δH** (500 MHz, $CDCl_3$) 8.19 (1 H, dd, *J* 3.9, 1.1, H-3), 8.03 (1 H, s, H-8), 7.57 (1 H, dd, *J* 5.0, 1.0, H-1), 7.10 (1 H, dd, *J* 5.0, 3.9, H-2), 3.48 (3 H, s, 3 × H-9 or 3 × H-10), 3.29 (3 H, s, 3 × H-9 or 3 × H-10).

**δC** (126 MHz, $CDCl_3$) 179.5 (C-5), 159.9 (C-8), 144.0 (C-4), 132.6 (C-1), 132.1 (C-3), 128.1 (C-2), 120.8 (C-7), 77.9 (C-6), 48.5 (C-9 or C-10), 39.2 (C-9 or C-10).

**HRMS (ESI+)** m/z calculated for $C_{10}H_{10}ON_2SNa$ $[M+Na]^+$ 229.0412, found 229.0403.

#### 3-(Thiophen-2-yl)-1H-pyrazole-5-carbonitrile (S8)[8]

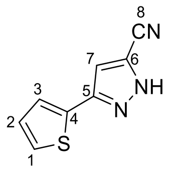

(*E*)-3-(Dimethylamino)-2-(thiophene-2-carbonyl)acrylonitrile (**S7**) (0.500 g, 2.42 mmol, 1.00 eq.) was dissolved in methanol (25 mL), and hydrazine monohydrate (64 wt% $NH_2NH_2$ in $H_2O$, 0.275 mL, 13.9 mmol, 1.50 eq.) was added. The reaction mixture was stirred for 16 hours at room temperature, and then concentrated *in vacuo*. The resulting crude oil was purified by silica gel column chromatography (0 – 5% MeOH in $CH_2Cl_2$) to give **S8** (0.351 g, 2.01 mmol, 83%) as a pale-yellow solid.

**M.P.** 204-205 °C (*lit*. 205-206 °C).[8]

**$\delta_H$** (500 MHz, DMSO-$d_6$) 8.59 (1 H, s, H-7), 7.67 (1 H, d, *J* 5.1, H-1), 7.63 (1 H, d, *J* 3.5, H-3), 7.23 – 7.18 (1 H, m, H-2).

Analytical data are in agreement with the literature.[8]

#### 1-(4-Methoxyphenyl)-*N*-((3-(thiophen-2-yl)-1H-pyrazol-5-yl)methyl)methanesulfonamide (VEST17)

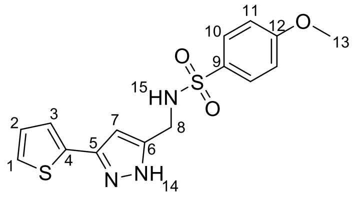

Synthesized in two steps via (3-(Thiophen-2-yl)-1H-pyrazol-4-yl)methanamine

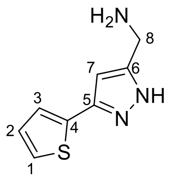

3-(Thiophen-2-yl)-1H-pyrazole-5-carbonitrile (**S8**) (0.200 g, 1.14 mmol, 1.00 eq.) was dissolved in dry THF (10 mL), and $LiAlH_4$ solution (2.4 M in THF, 2.85 mL, 6.84 mmol, 6.00 eq.) was added dropwise at 0 °C. The reaction mixture was stirred at 40 °C for 48 hours. The reaction was cooled to 0 °C, quenched by slow addition of cold EtOH (5 mL) followed by saturated aqueous $NH_4Cl$ solution (5 mL). The suspension was stirred for 2 hours at 0 °C, and then concentrated *in vacuo*. The resulting mixture was diluted with aqueous 1 M NaOH (10 mL), and the aqueous layer was extracted with $CH_2Cl_2$ (3 × 30 mL). The combined organic extracts were washed with brine (2 × 15 mL), dried ($MgSO_4$), filtered, and concentrated *in vacuo* to give the resulting amine, which was used in the next step without any further purification.

**$^1$H NMR analysis for amine**

**$\delta_H$** (400 MHz, $CDCl_3$) 7.51 (1 H, s, H-7), 7.31 – 7.29 (2 H, m, H-1, H-3), 7.09 (1 H, dd, *J* 4.9, 3.8, H-2), 3.97 (2 H, app. s, 2 × H-8).

The resulting amine (1.14 mmol, 1.00 eq.) was dissolved in dry $CH_2Cl_2$ (10 mL), and *i*-$Pr_2NEt$ (0.18 mL, 1.01 mmol, 1.05 eq.) was added dropwise at – 20 °C. The resulting mixture was stirred for 15 minutes, and 4-methoxybenzenesulfonyl chloride (0.208 g, 1.01 mmol, 1.05 eq.) was added. The reaction mixture was stirred at room temperature for 16 hours. The resulting mixture was quenched with saturated aqueous $NaHCO_3$ solution (10 mL), and diluted with $CH_2Cl_2$ (20 mL). The organic phase was washed with saturated aqueous $NaHCO_3$ solution (3 × 15 mL), dried ($MgSO_4$), filtered, and concentrated *in vacuo*. The resulting crude product was purified by silica gel column chromatography (0 – 10% MeOH in $CH_2Cl_2$) to give **VEST17** (0.0980 g, 0.270 mmol, 24% over two steps) as a white powder.

**Analytical characterization for VEST17**

**M.P.** 211-213 °C.

**IR $\nu_{max}$** (film) $cm^{-1}$ 3501, 3065, 2915, 2851, 1740, 1577, 1495, 1304, 1253, 1147, 1093, 1024, 829, 740.

$\delta_H$ (500 MHz, DMSO-$d_6$) 12.79 (1 H, s, NH-14), 7.77 (2 H, d, *J* 8.5, 2 × H-10), 7.72 (1 H, t, *J* 5.8, NH-15), 7.62 (1 H, s, H-7), 7.44 (1 H, d, *J* 5.1, H-1), 7.23 – 7.19 (1 H, m, H-3), 7.12 (1 H, d, *J* 8.5, 2 × H-11), 7.04 (1 H, t, *J* 4.4, H-2), 3.94 (2 H, d, *J* 5.4, 2 × H-8), 3.85 (3 H, s, 3 × H-13).

**$\delta_C$** (126 MHz, DMSO-$d_6$) 162.7 (C-12), 143.9 (C-5), 136.5 (C-4), 132.1 (C-9), 130.8 (C-7), 129.4 (C-10), 128.1 (C-2), 125.4 (C-1), 124.9 (C-3), 114.8 (C-11), 112.9 (C-6), 56.1 (C-13), 37.8 (C-8).

**HRMS (ESI$^+$)** m/z calculated for $C_{15}H_{15}O_3N_3S_2$ $[M+Na]^+$ 372.0447, found 372.0440.

#### General Scheme for the Synthesis of VEST18

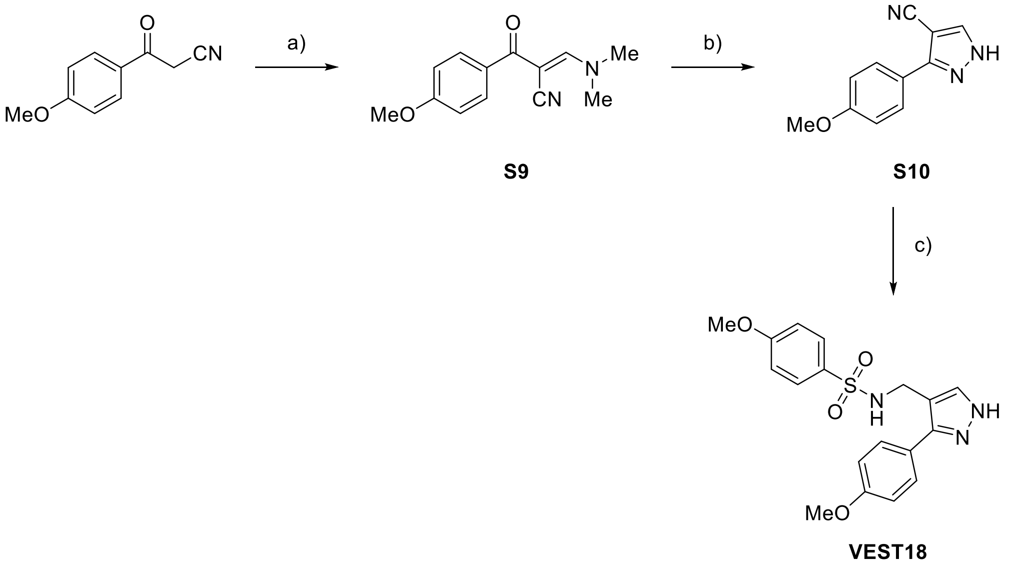


**Scheme S2** *Reagents and conditions* a) DMF-DMA (1.10 eq.), r.t., 16 hrs, 72%; b) $NH_2NH_2$ $H_2O$ (64 wt%, 1.50 eq.), MeOH, r.t., 16 hrs, 81%; c) i) $LiAlH_4$ (2.4 M in THF, 6.00 eq.), THF, 0 – 40 °C, 48 hrs; ii) 4-methoxybenzenesulfonyl chloride (1.05 eq.), *i*-$Pr_2NEt$ (1.05 eq.), $CH_2Cl_2$, – 20 °C – r.t., 16 hrs, 36% over two steps.

#### (*E*)-3-(Dimethylamino)-2-(4-methoxybenzoyl)acrylonitrile (S9)[9]

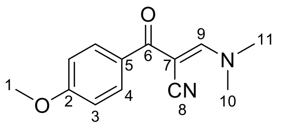

Commercially available 3-(4-methoxyphenyl)-3-oxopropanenitrile (1.50 g, 10.3 mmol, 1.00 eq.) was dissolved in dry toluene (34 mL), and *N,N*-dimethylformamide dimethyl acetal (DMF-DMA) (1.51 mL, 11.3 mmol, 1.10 eq.) was added dropwise. The reaction mixture was stirred for 16 hours at room temperature, and then concentrated *in vacuo*. The crude product was purified by silica gel column chromatography (0 – 5% MeOH in $CH_2Cl_2$) to give **S9** (1.70 g, 7.38 mmol, 72%) as a pale-yellow solid.

**M.P.** 90-91 °C (*lit.* 127-128 °C).[9]

**IR** $\nu_{max}$ (film) $cm^{-1}$ 2922, 2360, 2191, 1645, 1600, 1508, 1423, 1350, 1288, 1201, 1172, 765.

$\delta_H$ (500 MHz, $CDCl_3$) 7.93 (1 H, s, H-9), 7.83 (2 H, d, *J* 8.8, 2 × H-4), 6.91 (2 H, d, *J* 8.8, 2 × H-3), 3.84 (3 H, s, 1, 3 × H-1), 3.46 (3 H, s, 3 × H-10 or H-11), 3.27 (3 H, s, 3 × H-10 or H-11).

$\delta_C$ (126 MHz, $CDCl_3$) 189.0 (C-6), 162.5 (C-2), 159.5 (C-9), 131.0 (C-5), 130.7 (C-4), 120.9 (C-8), 113.4 (C-3), 79.3 (C-7), 55.5 (C-1) 48.3 (C-10 or C-11), 37.9 (C-10 or C-11).

**HRMS (ESI$^+$)** m/z calculated for $C_{13}H_{14}O_2N_2Na$ [M+Na]$^+$ 253.0953, found 253.0945.

#### 5-(4-Methoxyphenyl)-1H-pyrazole-4-carbonitrile (S10)[10]

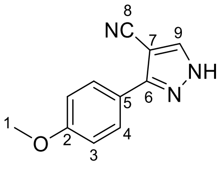

(*E*)-3-(dimethylamino)-2-(4-methoxybenzoyl)acrylonitrile (**S9**) (1.65 g, 7.17 mmol, 1.00 eq.) was dissolved in methanol (70 mL), and hydrazine monohydrate (64 wt% $NH_2NH_2$ in $H_2O$, 0.819 mL, 10.8 mmol, 1.50 eq.) was added. The reaction mixture was stirred for 16 hours at room temperature, and then concentrated *in vacuo*. The resulting crude was purified by silica gel column chromatography (0 – 2% MeOH in $CH_2Cl_2$) to give **S10** (1.16 g, 5.81 mmol, 81%) as a pale-yellow solid.

**M.P** 148-149 °C (*lit.* 150-152 °C).[10]

**$\delta_H$** (500 MHz, $CDCl_3$) 11.04 (1 H, s, NH), 7.98 (1 H, s, H-9), 7.81 (2 H, d, *J* 8.4, 2 × H-4), 7.07 – 7.01 (2 H, m, 2 × H-3), 3.81 (3 H, s, 3 × H-1)

Analytical data are in agreement with the literature.[10]

#### 4-Methoxy-*N*-((5-(4-methoxyphenyl)-1H-pyrazol-4-yl)methyl)benzenesulfonamide (VEST18)

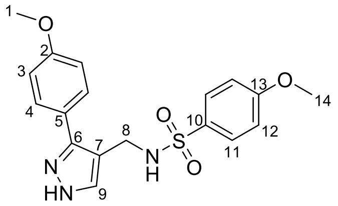

Synthesized in two steps via (3-(4-Methoxyphenyl)-1H-pyrazol-4-yl)methanamine

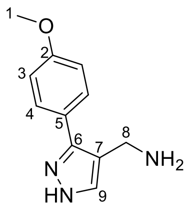

**S12**

3-(4-Methoxyphenyl)-1H-pyrazole-4-carbonitrile (**S10**) (0.250 g, 1.25 mmol, 1.00 eq.) was dissolved in dry THF (20 mL), and $LiAlH_4$ solution (2.4 M in THF, 3.14 mL, 7.50 mmol, 6.00 eq.) was added dropwise at 0 °C. The reaction mixture was stirred at 40 °C for 48 hours. The reaction was cooled to 0 °C, quenched by slow addition of cold EtOH (10 mL) followed by saturated aqueous $NH_4Cl$ solution (10 mL). The suspension was stirred for 2 hours at 0 °C, and then concentrated *in vacuo*. The resulting mixture was diluted with 1 M NaOH (15 mL), and the aqueous layer was extracted with $CH_2Cl_2$ (3 × 30 mL). The combined organic extracts were washed with brine (2 × 15 mL), dried ($MgSO_4$), filtered, and concentrated *in vacuo* to give the amine which was used in the next step without any further purification.

**$^1$H NMR analysis for amine**

**$\delta_H$** (400 MHz, $CDCl_3$) 7.57 (1 H, s, H-9), 7.51 (2 H, d, *J* 8.7, 2 × H-4), 6.95 (2 H, d, *J* 8.7, 2 × H-3), 3.90 (2 H, s, 2 × H-8), 3.84 (3 H, s, 3 × H-1).

The resulting amine (1.25 mmol, 1.00 eq.) was dissolved in dry $CH_2Cl_2$ (20 mL), and *i*-$Pr_2NEt$ (0.228 mL, 1.31 mmol, 1.05 eq.) was added dropwise at – 20 °C. The resulting mixture was stirred for 15 minutes, and 4-methoxybenzenesulfonyl chloride (0.270 g, 1.31 mmol, 1.05 eq.) was added. The reaction mixture was stirred at room temperature for 16 hours. The resulting mixture was quenched with saturated aqueous $NaHCO_3$ solution (15 mL), and diluted with $CH_2Cl_2$ (30 mL). The organic phase was washed with saturated aqueous $NaHCO_3$ solution (3 × 15 mL), dried ($MgSO_4$), filtered, and concentrated *in vacuo*. The resulting crude was purified by silica gel column chromatography (0 – 10% MeOH in $CH_2Cl_2$) to give **VEST18** (0.150 g, 0.401 mmol, 36% over two steps) as a white powder.

**Analytical characterization for VEST18**

**M.P.** 141-142 °C.

**IR $\nu_{max}$** (film) $cm^{-1}$ 3267, 2950, 2361, 2341,1597, 1496, 1301, 1257, 1150, 833.

**$\delta_H$** (500 MHz, $CDCl_3$) 7.76 (2 H, d, *J* 8.9, 2 × H-11), 7.37 (1 H, s, H-9), 7.30 (2 H, d, *J* 8.8, 2 × H-4), 6.95 (2 H, d, *J* 8.9, 2 × H-12), 6.85 (2 H, d, *J* 8.8, 2 × H-3), 4.96 (1 H, t, *J* 5.1, NH), 4.09 (2 H, d, *J* 5.1, 2 × H-8), 3.87 (3 H, s, 3 × H-14), 3.82 (3 H, s, 3 × H-1).

**$\delta_C$** (126 MHz, $CDCl_3$) 163.1 (C-13), 160.3 (C-2), 145.1 (C-6), 136.0 (C-9), 131.2 (C-10), 129.5 (C-11), 128.9 (C-4), 122.9 (C-5), 114.5 (C-3), 114.4 (C-12), 112.7 (C-7), 55.8 (C-14), 55.4 (C-1), 36.8 (C-8).

**HRMS (ESI$^+$)** m/z calculated for $C_{18}H_{19}N_3O_4SNa$ $[M+Na]^+$ 396.0994, found 396.0987.

#### General Scheme for the Synthesis of VEST19

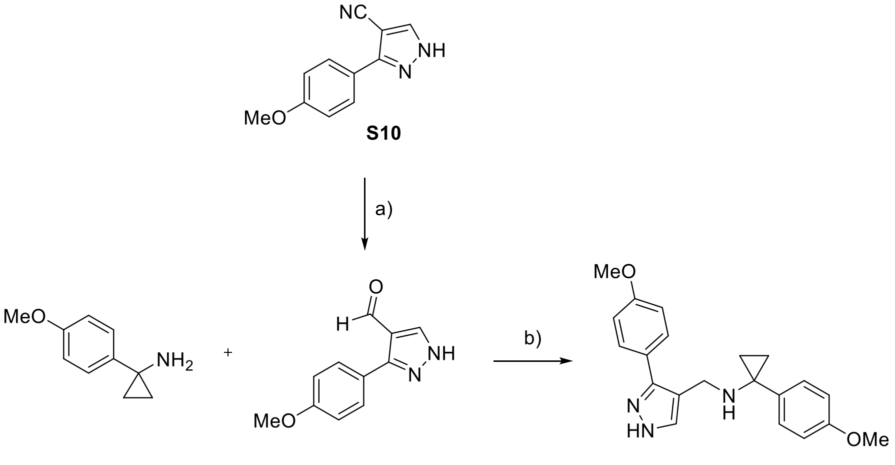


**Scheme S3** *Reagents and conditions* a) DIBAL-H (1.2 M in toluene, 6.00 eq.), $CH_2Cl_2$, – 20 °C – r.t., 3 hrs, 76%; b) i) **S13** (1.30 eq.), THF, r.t., 16 hrs; ii) $NaBH_4$ (eq.), MeOH, 0 °C – r.t, 30 mins, 56% over two steps.

#### 3-(4-Methoxyphenyl)-1H-pyrazole-4-carbaldehyde (S13)[11]

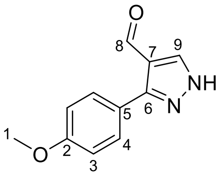


3-(4-Methoxyphenyl)-1H-pyrazole-4-carbonitrile (**S10**) (0.100, 0.502 mmol, 1.00 eq.) was dissolved in dry $CH_2Cl_2$, and DIBAL-H solution (1.2 M in toluene, 2.50 mL, 3.01 mmol, 6.00 eq.) was added dropwise at – 20 °C. The reaction mixture was stirred at room temperature for 3 hours. After cooling to 0 °C, the reaction mixture was quenched by slow addition of cold MeOH (3 mL) followed by a saturated aqueous solution of Rochelle salt (5 mL). The resulting mixture was stirred for 30 minutes at – 20 °C, and then concentrated *in vacuo*. The resulting crude reaction mixture was diluted with water (10 mL), and the aqueous layer was extracted with EtOAc (3 × 20 mL). The combined organic extracts were dried ($MgSO_4$), filtered, and concentrated *in vacuo*. The resulting crude was purified by silica gel column chromatography (0 – 2% MeOH in $CH_2Cl_2$) to give **S13** (0.0780 g, 0.386 mmol, 76%) as a white solid.

**M.P.** 167-168 °C (*lit.* 168-169 °C).[11]

**$\delta_H$** (500 MHz, DMSO-$d_6$) 9.86 (1 H, s, H-8), 8.28 (1 H, s, H-9), 7.79 – 7.73 (2 H, m, 2 × H-4), 7.09 – 7.03 (2 H, m, 2 × H-3), 3.81 (3 H, s, 3 × H-1).

Analytical data were in agreement with the literature.[11]

##### 1-(4-Methoxyphenyl)-*N*-((3-(4-methoxyphenyl)-1H-pyrazol-4-yl)methyl)cyclopropan-1-amine (VEST19)

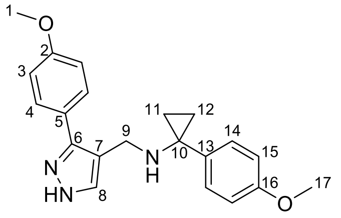


Synthesized in two steps via (*E*)-1-(3-(4-Ethoxyphenyl)-1H-pyrazol-4-yl)-N-(1-(4-methoxyphenyl)cyclopropyl)methanimine

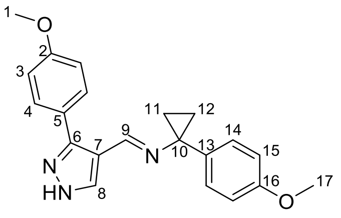


3-(4-Ethoxyphenyl)-1H-pyrazole-4-carbaldehyde (**S13**) (0.0400 mg, 0.245 mmol, 1.00 eq.) and 1-(4-methoxyphenyl)cyclopropan-1-amine (**S5**) (0.0644 g, 0.318 mmol, 1.30 eq.) were dissolved in dry THF (3 mL). The resulting mixture was stirred for 16 hours at room temperature, and then concentrated *in vacuo* to give the imine, which was used immediately in the next step without any further purification.

**$^1$H NMR analysis of imine**

**$\delta_H$** (500 MHz, $CDCl_3$) 8.00 (1 H, s, H-8), 7.82 (1 H, s, H-9), 7.29 (2 H, d, *J* 8.7, 2 × H-4), 7.25 (2 H, d, *J* 4.9, 2 × H-14), 6.91 – 6.82 (4 H, m, 2 × H-3, 2 × H-15), 3.83 (3 H, s, 3 × H-1), 3.80 (3 H, s, 3 × H-17), 1.41 – 1.38 (2 H, m, 1 × H-11, 1 × H-12), 1.25 – 1.22 (1 H, m, 1 × H-11, 1 × H-12).

The resulting imine was dissolved in dry MeOH (3 mL), and $NaBH_4$ (12 mg, 0.319 mmol, 1.30 eq.) was added at 0 °C. The reaction mixture was stirred at room temperature for 30 minutes. The reaction mixture was cooled to 0 °C and quenched with saturated aqueous $NH_4Cl$ solution (5 mL), and water (5 mL). The aqueous layer was extracted with EtOAc (3 × 15 mL). The combined organic extracts were dried ($MgSO_4$), filtered, and concentrated *in vacuo*. The crude product was purified by silica gel column chromatography (0 – 10% MeOH in $CH_2Cl_2$) to give **S14** (0.0480 g, 0.137 mmol, 56% over two steps) as a white powder.

**Analytical characterization for VEST19**

**M.P.** 158-159 °C.

**IR $\nu_{max}$** (film) $cm^{-1}$ 3301, 3150, 2920, 2349, 1603, 1516, 1436, 1288, 1202, 1170, 1033, 820.

**$\delta_H$** (500 MHz, $CDCl_3$) 7.48 (1 H, s, H-8), 7.41 (2 H, d, *J* 8.3, 2 × H-4), 7.28 (2 H, d, *J* 2.8, 2 × H-14), 6.90 – 6.81 (4 H, m, 2 × H-3, 2 × H-15), 3.84 (3 H, s, 3 × H-1), 3.82 (3 H, s, 3 × H-17), 3.65 (2 H, s, 2 × H-9), 1.04 – 0.98 (2 H, m, 1 × H-11, 1 × H-12), 0.93 – 0.82 (2 H, m, 1 × H-11, 1 × H-12).

**$\delta_C$** (126 MHz, $CDCl_3$) 159.6 (C-2), 158.5 (C-16), 144.9 (C-6, determined through HMBC), 135.7 (C-8, determined through HSQC), 134.9 (C-13), 129.6 (C-14), 128.8 (C-4), 124.0 (C-5), 116.3 (C-7), 114.2 (C-3), 113.8 (C-15), 55.4 (C-1), 55.4 (C-17), 42.3 (C-10), 40.2 (C-9), 14.8 (C-11, C-12).

**HRMS ($ESI^+$)** m/z calculated for $C_{21}H_{24}O_2N_3$ $[M+H]^+$ 350.1863, found 350.1862.

### NMR spectra

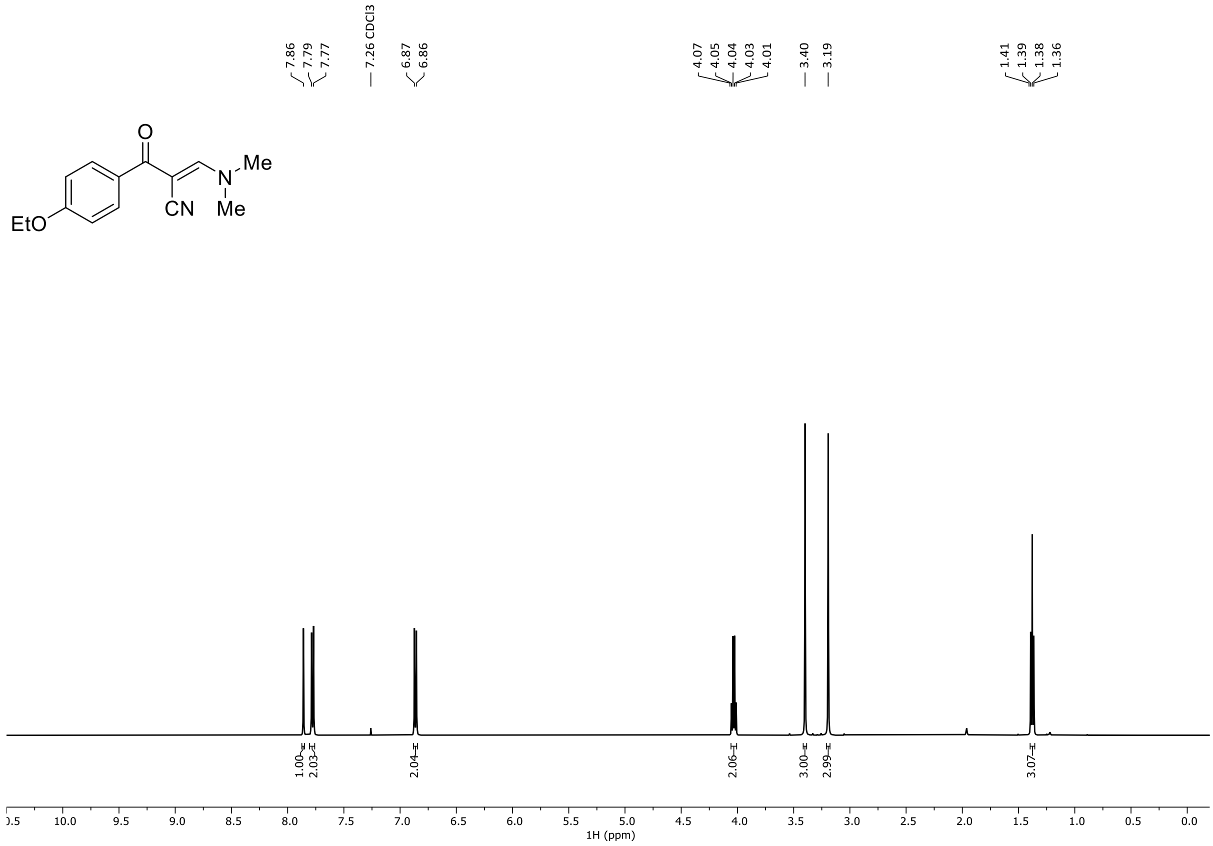
7.86
7.79
7.77
7.26 CDCl3
6.87
6.86
4.07
4.05
4.04
4.03
4.01
3.40
3.19
1.41
1.39
1.38
1.36
O
Me
N
CN
Me
EtO
1.00
2.03
2.04
2.06
3.00
2.99
3.07
1H (ppm)

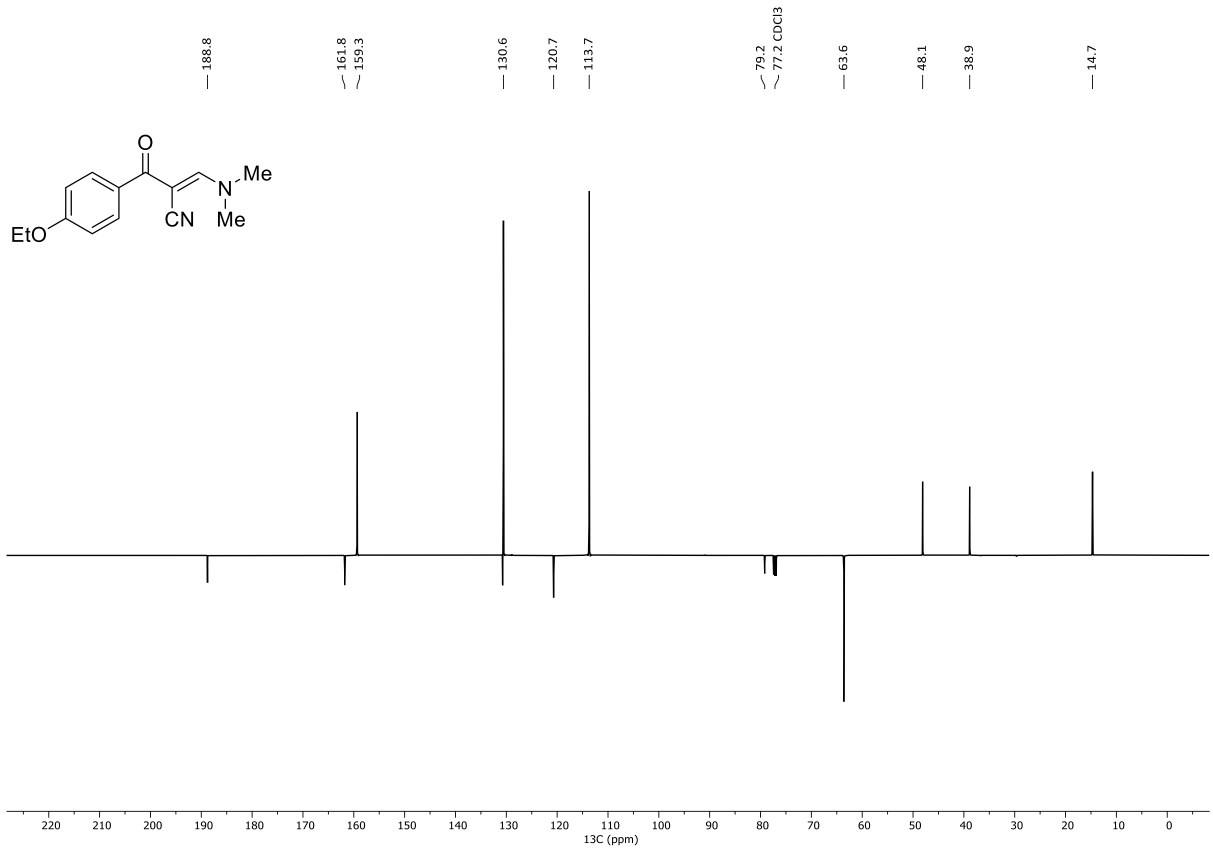
188.8
161.8
159.3
130.6
120.7
113.7
79.2
77.2 CDCl3
63.6
48.1
38.9
14.7
O
Me
N
CN
Me
EtO
13C (ppm)

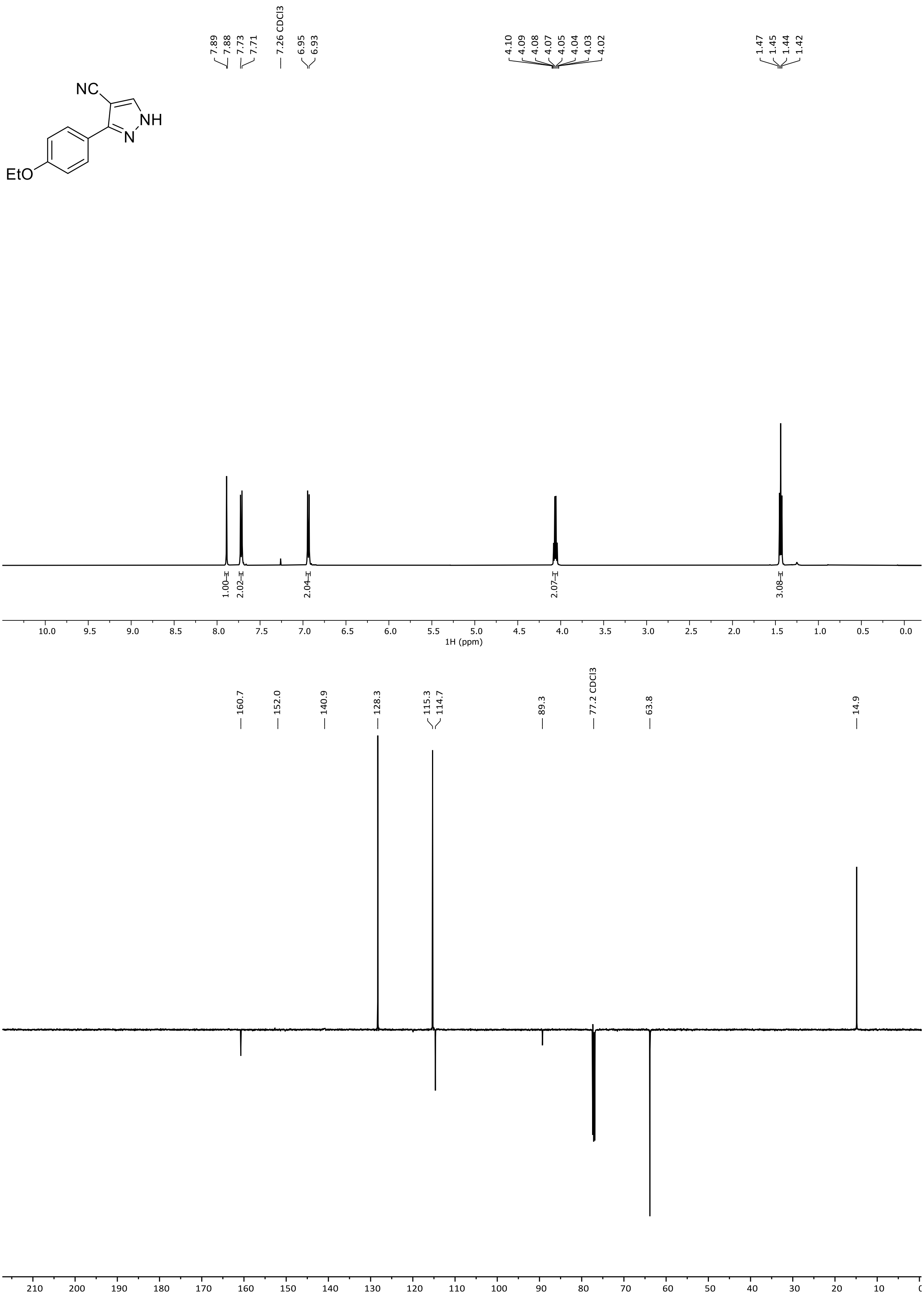
NC
NH
N
EtO
7.89
7.88
7.73
7.71
7.26 CDCl3
6.95
6.93
4.10
4.09
4.08
4.07
4.05
4.04
4.03
4.02
1.47
1.45
1.44
1.42
1.00
2.02
2.04
2.07
3.08
1H (ppm)
160.7
152.0
140.9
128.3
115.3
114.7
89.3
77.2 CDCl3
63.8
14.9
13C (ppm)

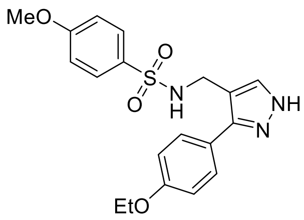
MeO
S
O
O
N
H
NH
N
EtO

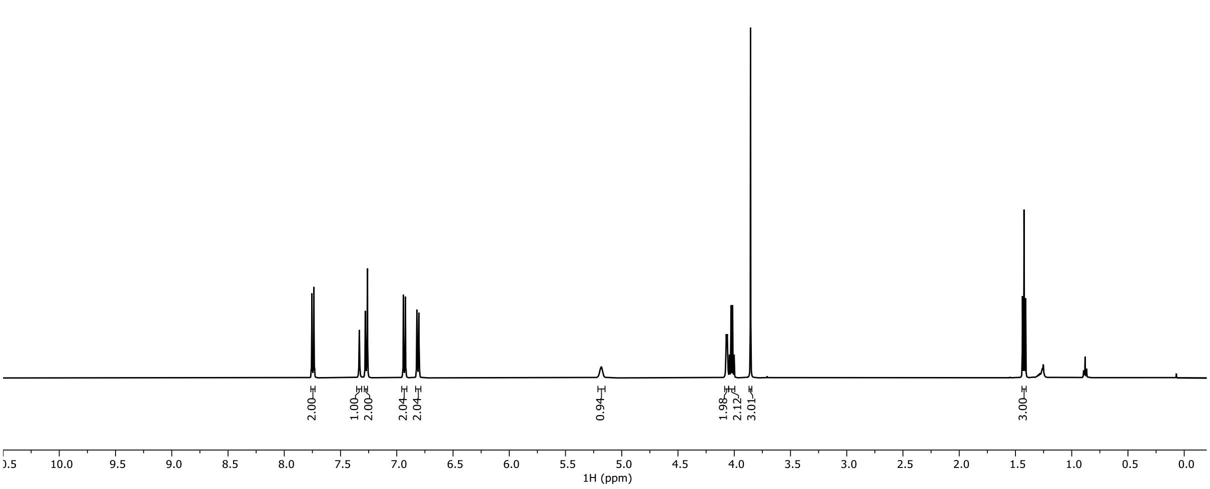
7.75
7.74
7.33
7.28
7.27
7.26
7.26 CDCl3
7.26
6.95
6.94
6.92
6.82
6.80
5.18
4.07
4.06
4.04
4.03
4.01
4.00
3.86
1.44
1.42
1.41
2.00
1.00
2.00
2.04
2.04
0.94
1.98
2.12
3.01
3.00
1H (ppm)

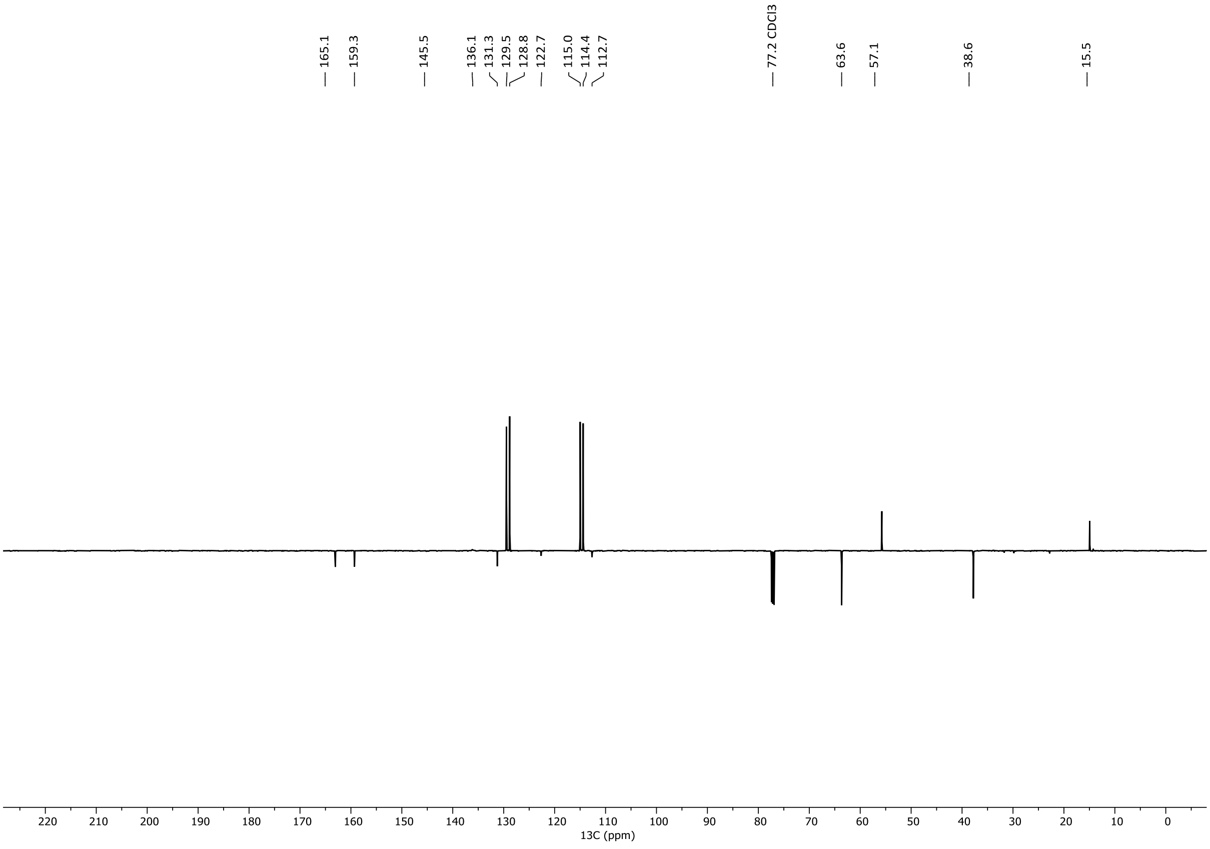
165.1
159.3
145.5
136.1
131.3
129.5
128.8
122.7
115.0
114.4
112.7
77.2 CDCl3
63.6
57.1
38.6
15.5
13C (ppm)

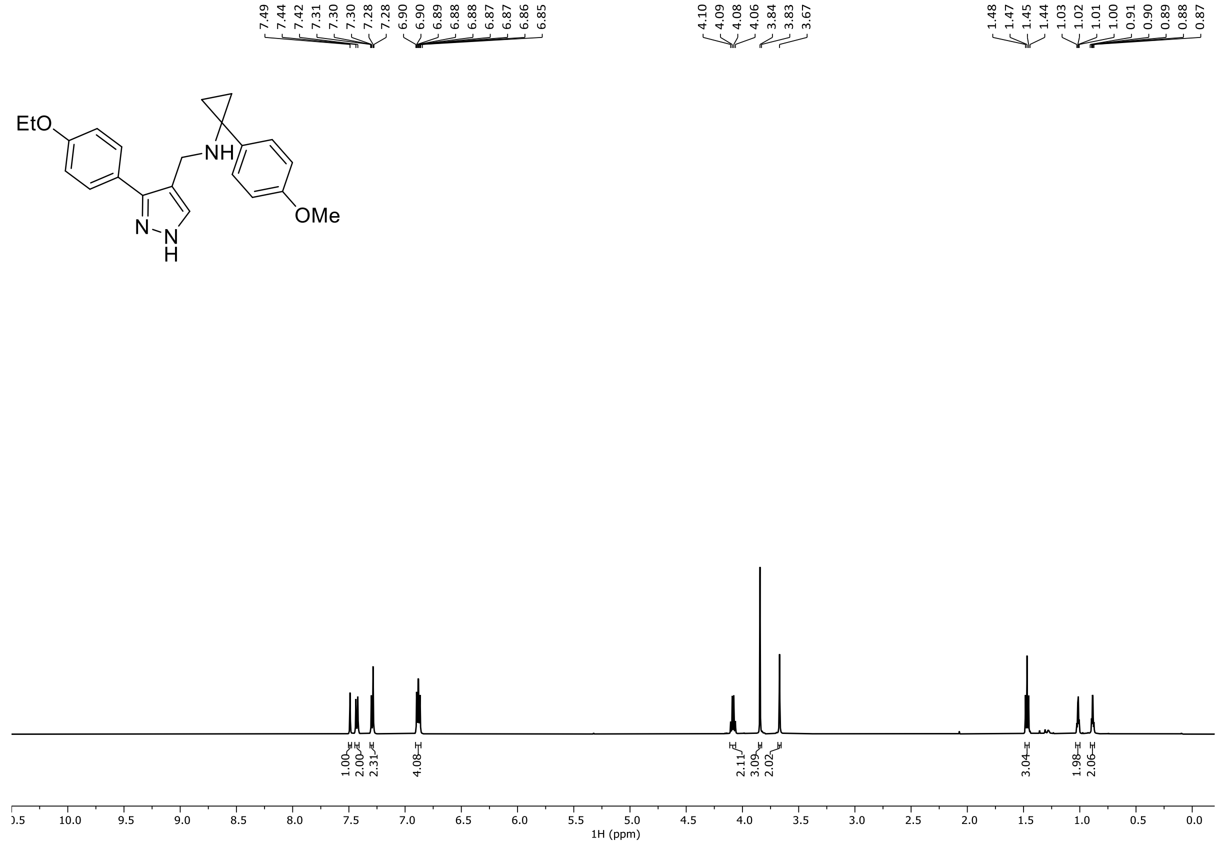
7.49 7.44 7.42 7.31 7.30 7.30 7.28 7.28 6.90 6.90 6.89 6.88 6.88 6.87 6.87 6.86 6.85
4.10 4.09 4.08 4.06 3.84 3.83 3.67
1.48 1.47 1.45 1.44 1.03 1.02 1.01 1.00 0.91 0.90 0.89 0.88 0.87
EtO
NH
N
N
H
OMe
1.00 2.00 2.31 4.08 2.11 3.09 2.02 3.04 1.98 2.06
1H (ppm)

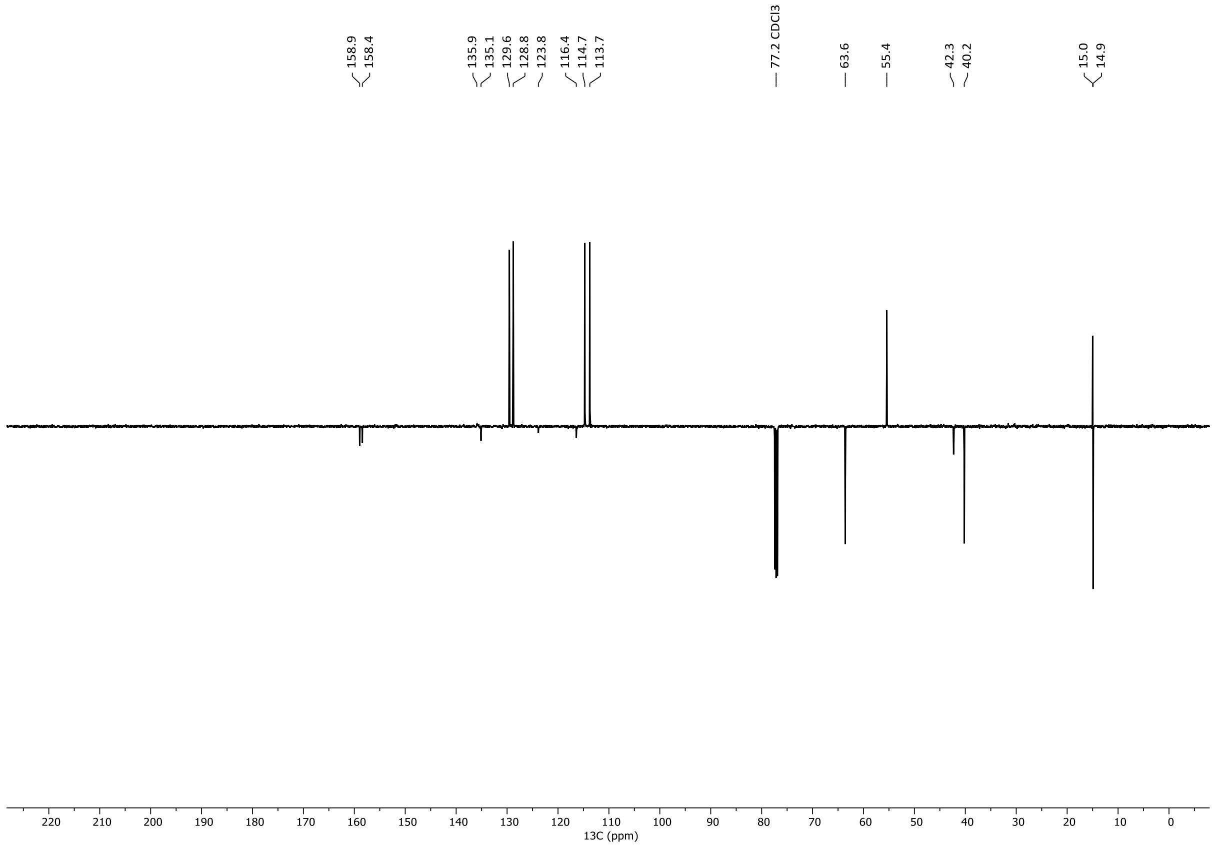
158.9 158.4 135.9 135.1 129.6 128.8 123.8 116.4 114.7 113.7
77.2 CDCl3
63.6 55.4 42.3 40.2 15.0 14.9
13C (ppm)

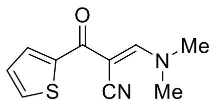
O
N
Me
S
CN
Me

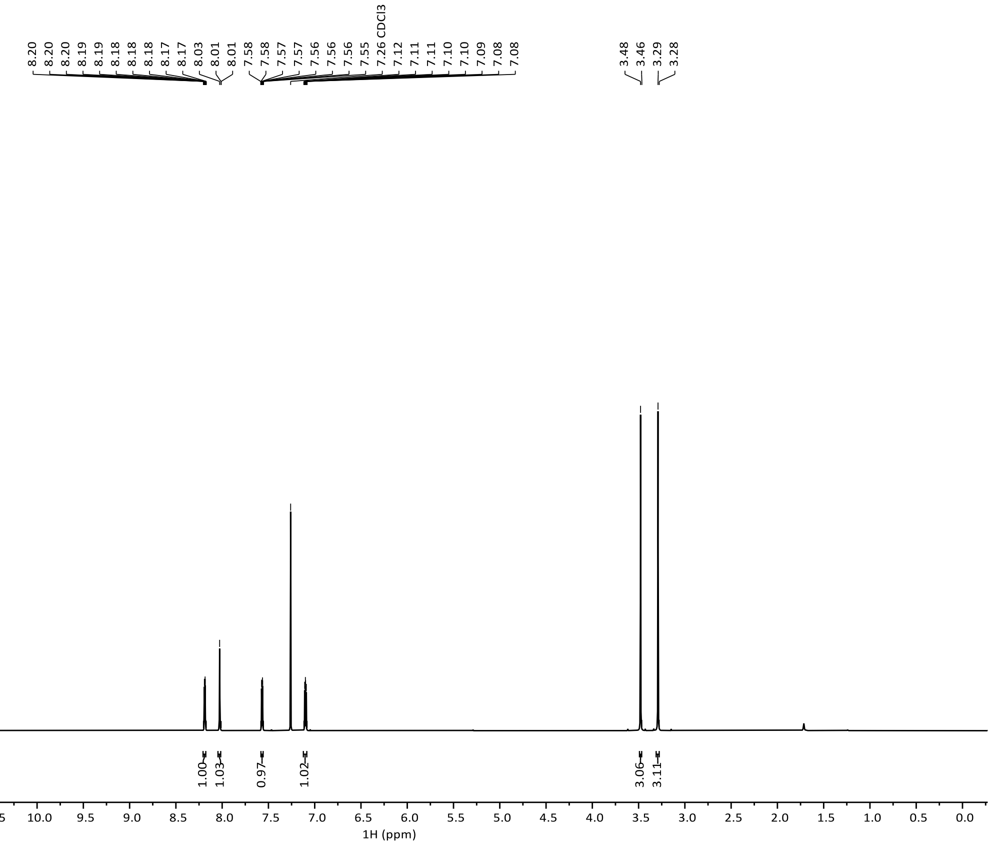
8.20 8.20 8.20 8.19 8.19 8.18 8.18 8.18 8.17 8.17 8.03 8.01 8.01 7.58 7.58 7.57 7.57 7.56 7.56 7.56 7.55 7.26 CDCl3 7.12 7.11 7.11 7.10 7.10 7.09 7.08 7.08
3.48 3.46 3.29 3.28
1.00 1.03 0.97 1.02 3.06 3.11
12.5 12.0 11.5 11.0 10.5 10.0 9.5 9.0 8.5 8.0 7.5 7.0 6.5 6.0 5.5 5.0 4.5 4.0 3.5 3.0 2.5 2.0 1.5 1.0 0.5 0.0
1H (ppm)

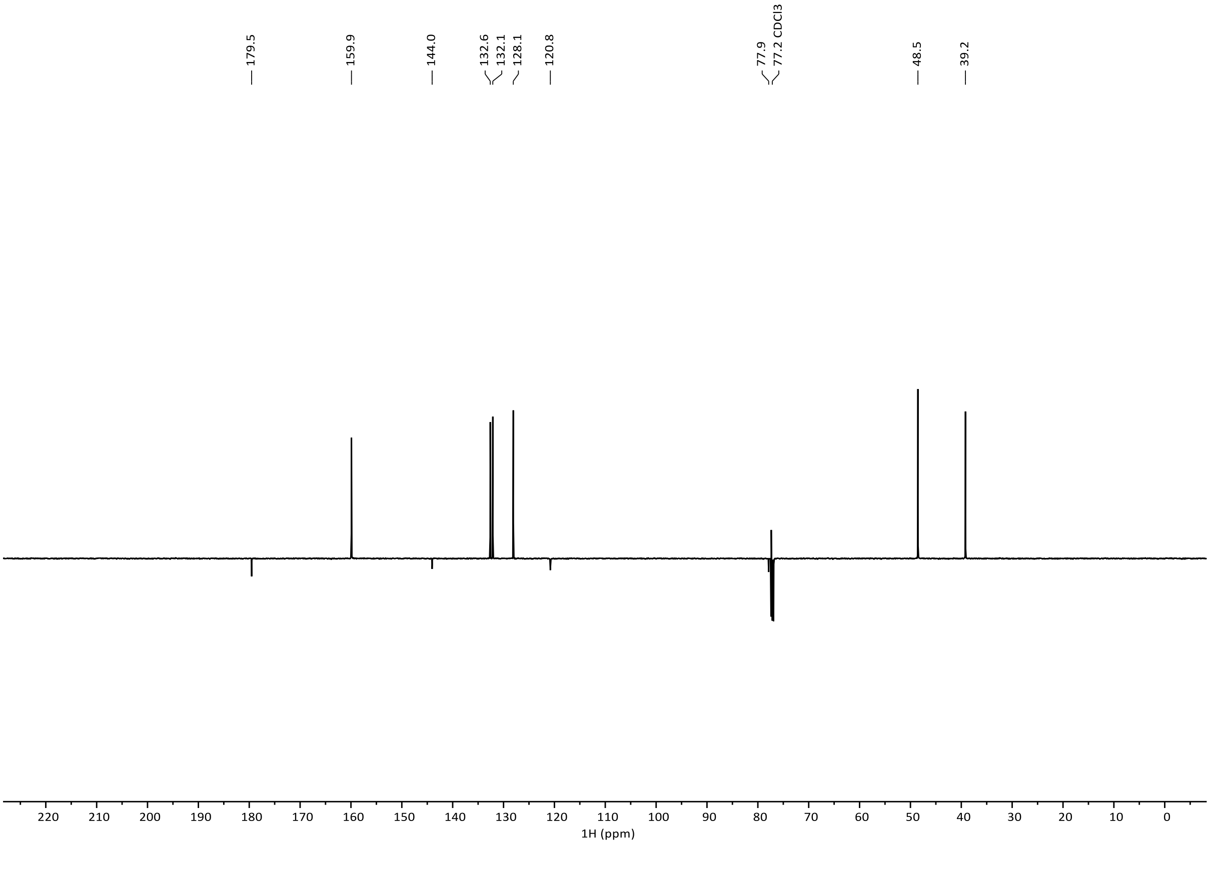
179.5
159.9
144.0
132.6 132.1 128.1
120.8
77.9 77.2 CDCl3
48.5
39.2
220 210 200 190 180 170 160 150 140 130 120 110 100 90 80 70 60 50 40 30 20 10 0
1H (ppm)

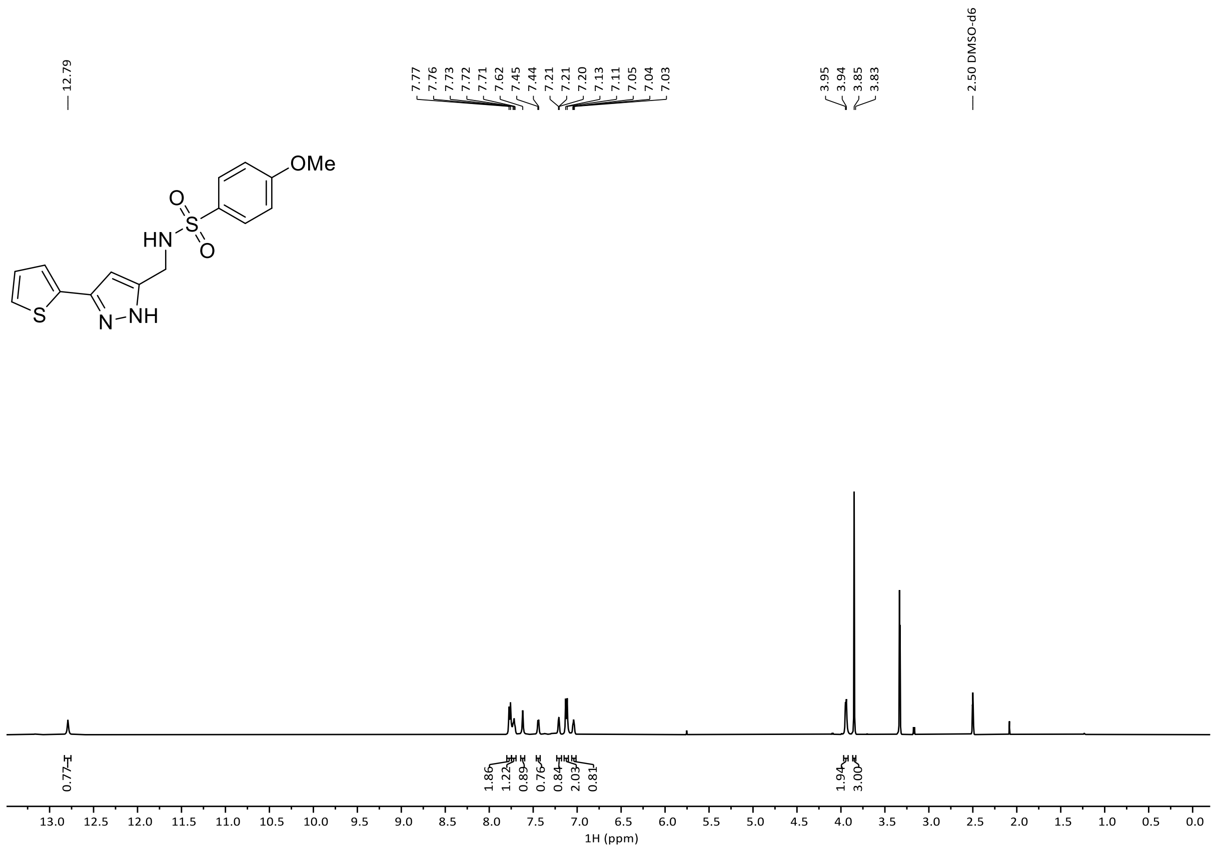
12.79
7.77
7.76
7.73
7.72
7.71
7.62
7.45
7.44
7.21
7.21
7.20
7.13
7.11
7.05
7.04
7.03
3.95
3.94
3.85
3.83
2.50 DMSO-d6
OMe
O
S
O
HN
S
N
NH
0.77
1.86
1.22
0.89
0.76
0.84
2.03
0.81
1.94
3.00
1H (ppm)

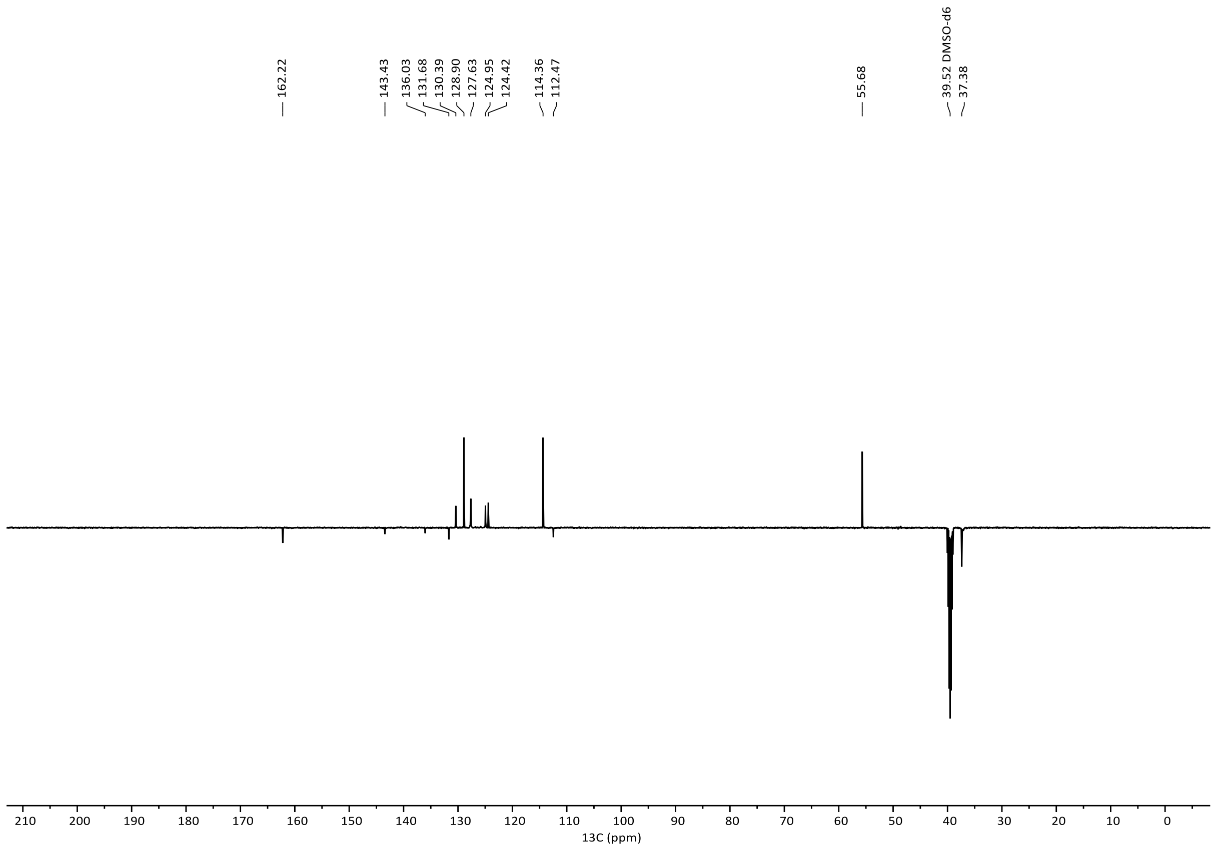
162.22
143.43
136.03
131.68
130.39
128.90
127.63
124.95
124.42
114.36
112.47
55.68
39.52 DMSO-d6
37.38
13C (ppm)

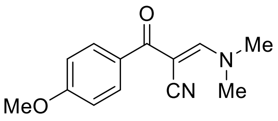
O
Me
N
MeO
CN
Me

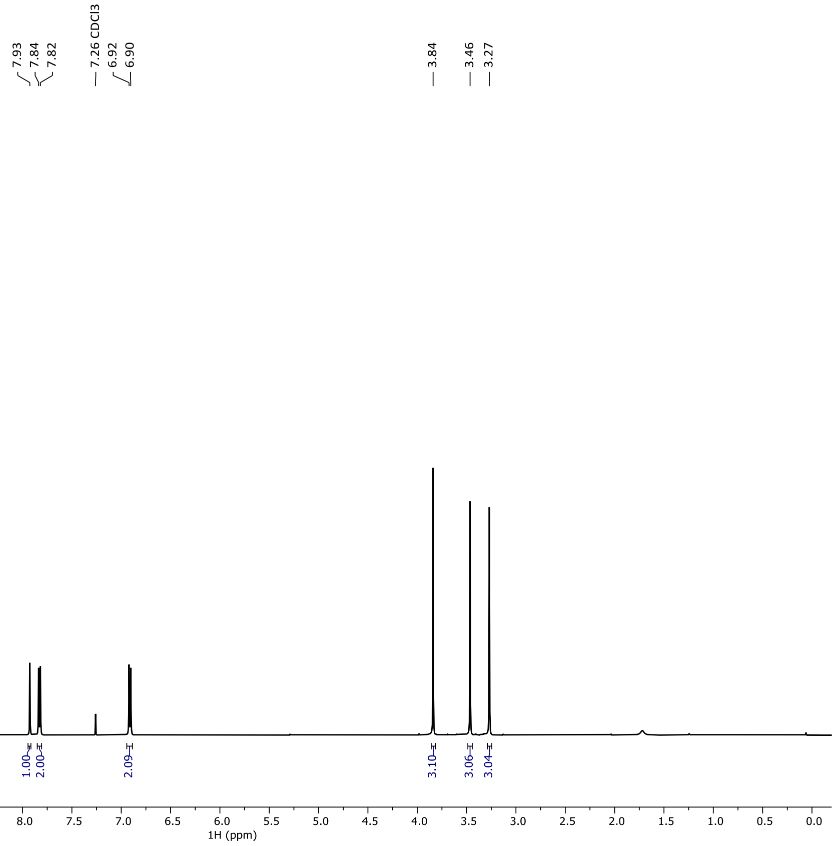
7.93
7.84
7.82
7.26 CDCl3
6.92
6.90
3.84
3.46
3.27
1.00
2.00
2.09
3.10
3.06
3.04
1H (ppm)

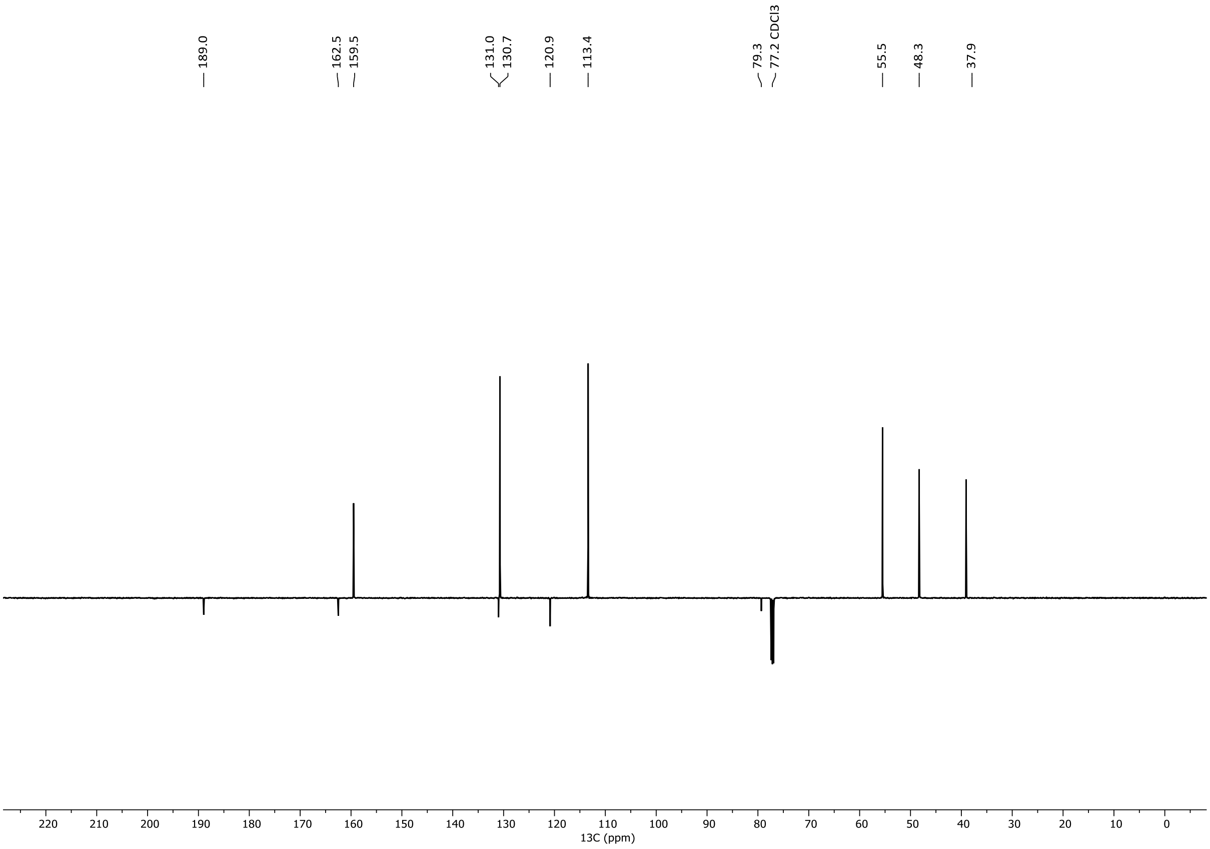
189.0
162.5
159.5
131.0
130.7
120.9
113.4
79.3
77.2 CDCl3
55.5
48.3
37.9
13C (ppm)

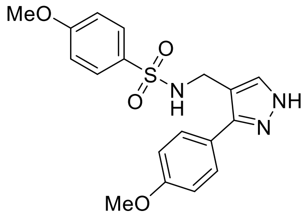
MeO
O
S
O
N
H
NH
N
MeO

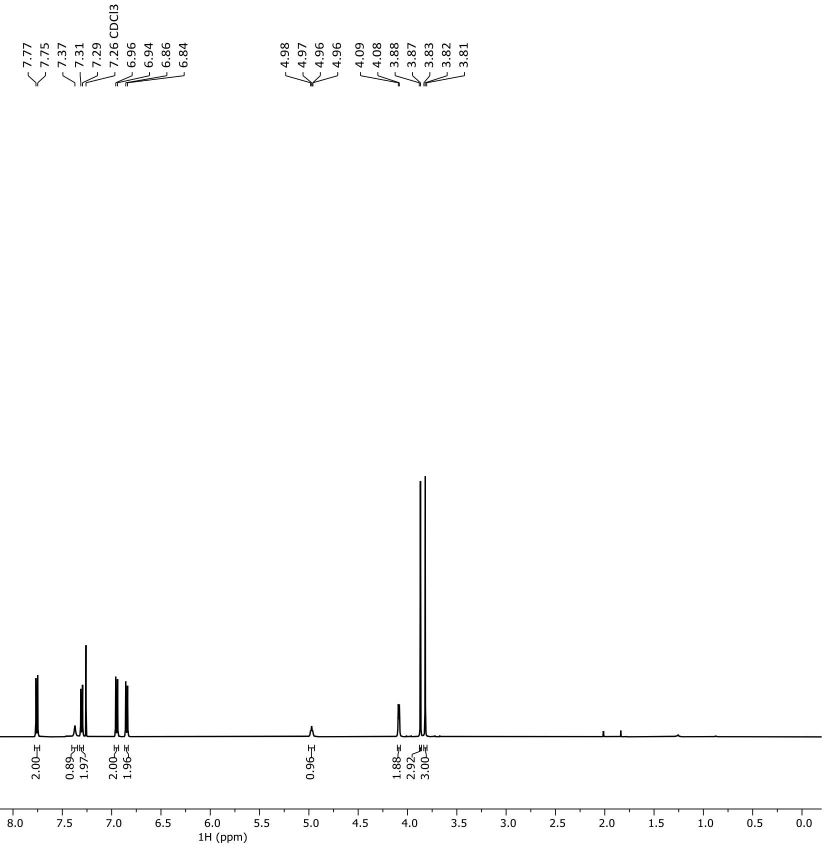
7.77
7.75
7.37
7.31
7.29
7.26 CDCl3
6.96
6.94
6.86
6.84
4.98
4.97
4.96
4.96
4.09
4.08
3.88
3.87
3.83
3.82
3.81
2.00
0.89
1.97
2.00
1.96
0.96
1.88
2.92
3.00
1H (ppm)

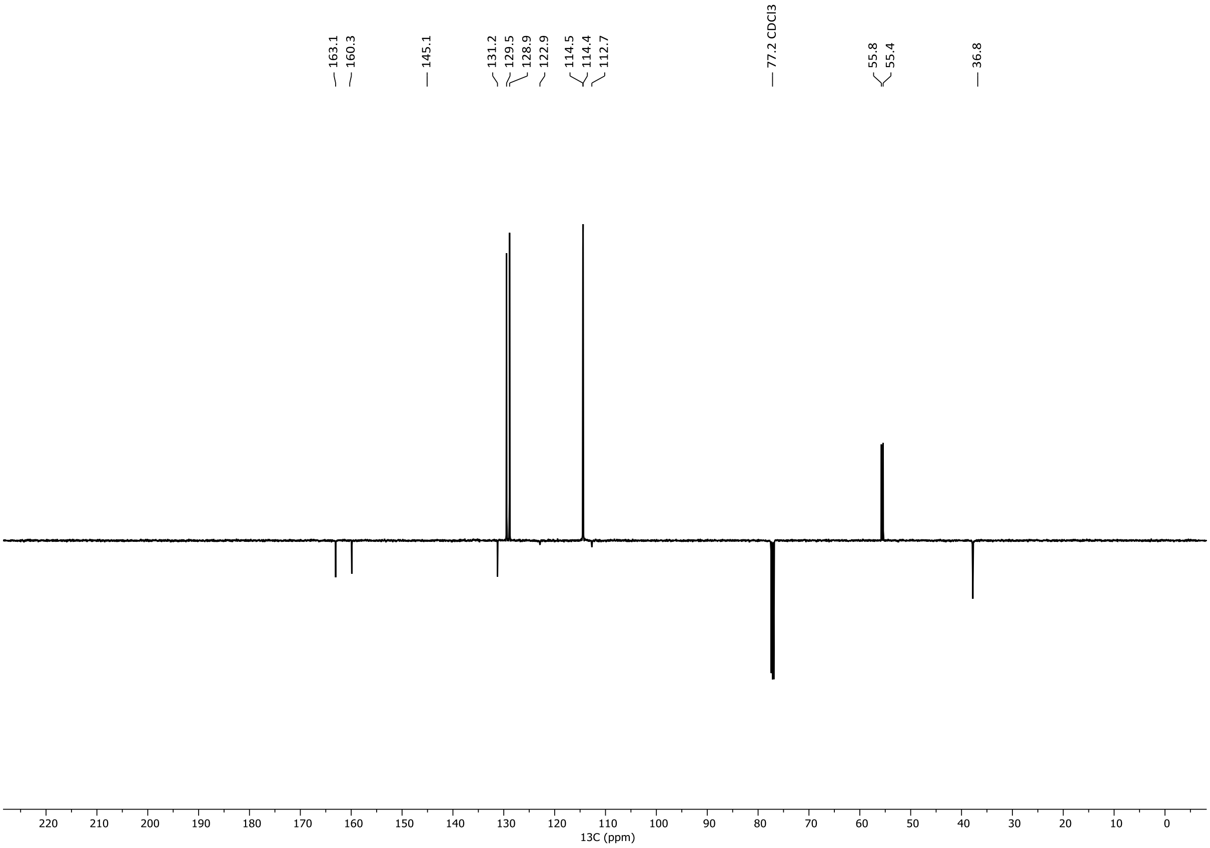
163.1
160.3
145.1
131.2
129.5
128.9
122.9
114.5
114.4
112.7
77.2 CDCl3
55.8
55.4
36.8
13C (ppm)

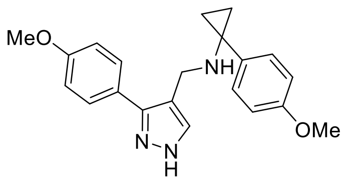
MeO
NH
N
N
H
OMe

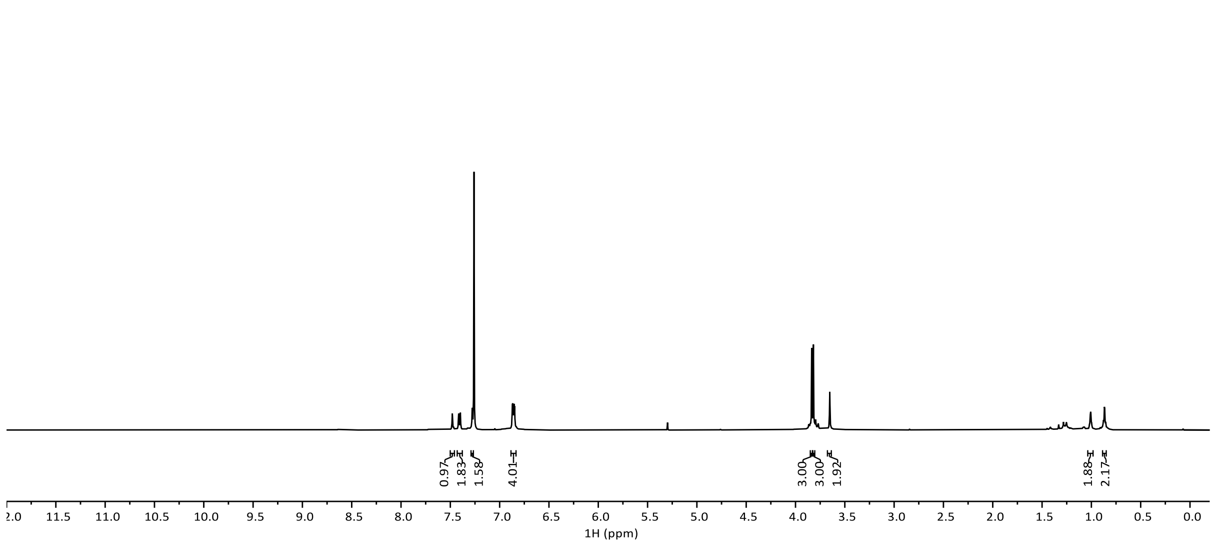
7.48
7.47
7.42
7.40
7.28
7.28
7.27
7.26 CDCl3
6.87
6.87
6.85
6.85
6.84
6.83
6.83
3.84
3.82
3.65
3.64
1.02
1.01
1.01
1.00
0.88
0.88
0.87
0.86
0.85
0.84
0.97
1.83
1.58
4.01
3.00
3.00
1.92
1.88
2.17
1H (ppm)

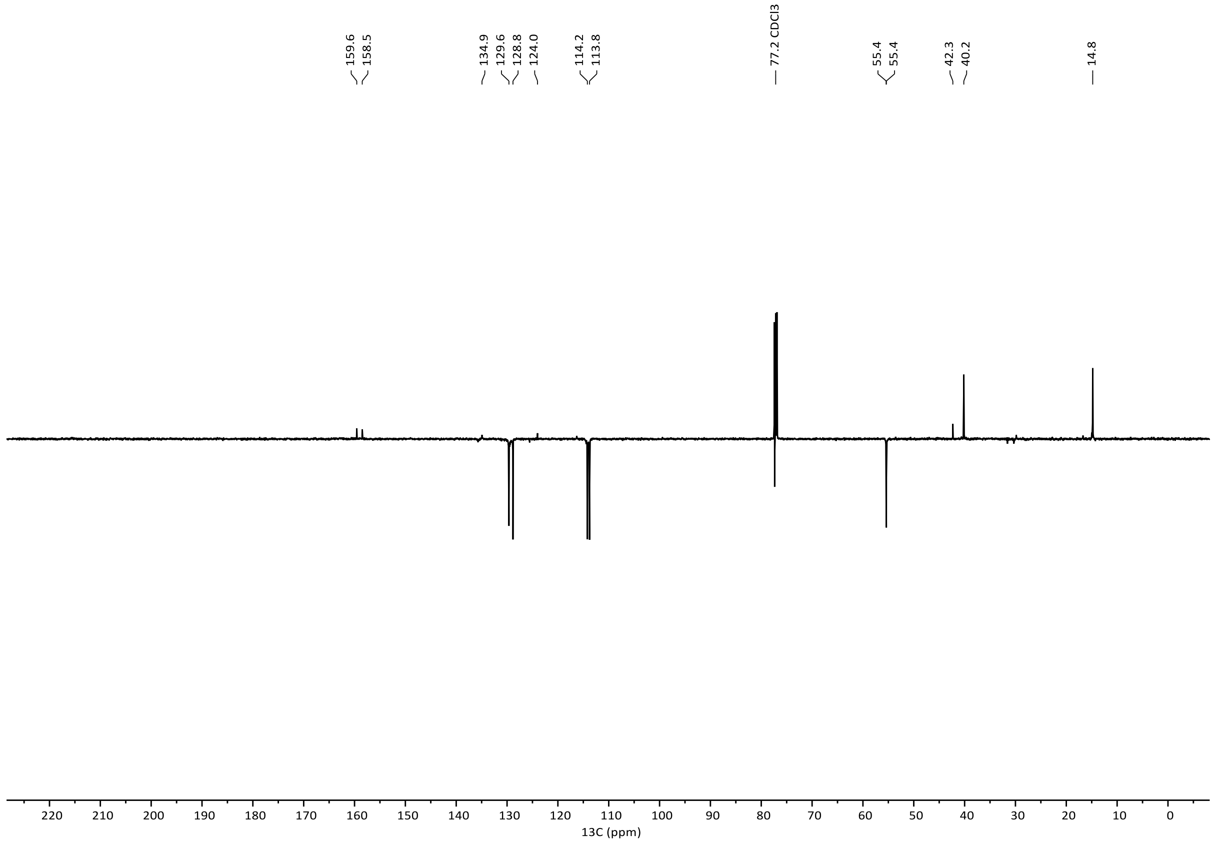
159.6
158.5
134.9
129.6
128.8
124.0
114.2
113.8
77.2 CDCl3
55.4
55.4
42.3
40.2
14.8
13C (ppm)
